# PTEN deficiency exposes a requirement for an ARF GTPase module in integrin-dependent invasion in ovarian cancer

**DOI:** 10.1101/2022.11.29.518198

**Authors:** Konstantina Nikolatou, Emma Sandilands, Alvaro Román-Fernández, Erin M. Cumming, Eva Freckmann, Sergio Lilla, Lori Buetow, Lynn McGarry, Matthew Neilson, Robin Shaw, David Strachan, Crispin Miller, Danny T. Huang, Iain A. McNeish, James C. Norman, Sara Zanivan, David M. Bryant

## Abstract

Dysregulation of the PI3K/AKT pathway is a common occurrence in ovarian carcinomas. Loss of the tumour suppressor *PTEN* in high-grade serous ovarian carcinoma (HGSOC) is associated with a patient subgroup with poor prognosis. The cellular mechanisms of how *PTEN* loss contributes to HGSOC are largely unknown. We utilise long-term time-lapse imaging of HGSOC spheroids coupled to a machine learning approach to classify the phenotype of *PTEN* loss. *PTEN* deficiency does not affect proliferation but rather induces PI(3,4,5)P_3_-rich and -dependent membrane protrusions into the extracellular matrix (ECM), resulting in a collective invasion phenotype. We identify the small GTPase ARF6 as a crucial vulnerability upon *PTEN* loss. Through a functional proteomic CRISPR screen of ARF6 interactors, we identify the ARF GTPase-activating protein (GAP) AGAP1 and the ECM receptor β1-integrin (ITGB1) as key ARF6 interactors regulating the *PTEN* loss-associated invasion phenotype. ARF6 functions to promote invasion by controlling the recycling of internalised, active β1-integrin complexes to maintain invasive activity into the ECM. The expression of the ARF6-centred complex in HGSOC patients is inversely associated with outcome, allowing identification of patient groups with improved versus poor outcome. ARF6 may represent a new therapeutic vulnerability in *PTEN*- depleted HGSOC tumours.

## Main

The tumour suppressor PTEN is a dual specificity phosphatase regulating both protein tyrosine dephosphorylation ^1^ as well as dephosphorylation of the 3- positions of phosphatidyl-inositol-3,4,5-*tris*-phosphate (PI(3,4,5)P_3_, PIP_3_) ^2^ and phosphatidyl-inositol-3,4-*bis*-phosphate (PI(3,4)_2_) ^3^. In a classical view of lipid phosphatase function PTEN acts as a buffer to oppose potential over-production of PIP_3_ or PI(3,4)_2_. This ensures the appropriate level of downstream pathway activation and homeostatic responses to PI3K signalling ^4, 5^. In addition to their well-documented roles in cell signalling, such as to the AKT and mTOR pathways ^6–8^ the spatial distribution of PIP_3_ or PI(3,4)_2_ is integral to their contribution to cell behaviour. Specifically, the location of these two PTEN-regulated PIP species is asymmetric in polarised epithelial cells; PIP_3_ is focally enriched at the basolateral surface ^9^ while PI(3,4)_2_ is located at the apical domain ^10^. PTEN is present at the apico-laterally localised tight junction, which is a boundary point between these asymmetric lipids ^11^.

*PTEN* gene deletion can be found in a number of cancers, particularly high-grade serous ovarian carcinoma (HGSOC) and prostate cancers ^12, 13^. Mutation of *PTEN* also occurs at a modest level in most cancers, with Glioblastoma and Uterine cancers presenting frequent *PTEN* mutation (The Cancer Genome Atlas (TCGA), cBioPortal^14, 15^). Mutation of the PIP_3_-producing *PIK3CA*, in contrast, is a frequent event in a number of cancers ^16^. This emphasises that dysregulation of the PI3K- PTEN axis is a common event in several cancer types ^17^. Despite this, exactly how these lipid kinase and phosphatases enact the cellular changes that contribute to tumourigenesis remains largely unclear. For instance, given the polarised nature of these lipids, does the loss of *PTEN* allow for enhanced signalling function at the normal site of PIP_3_ in the cell (the basolateral domain) or is PIP_3_ produced at ectopic sites, allowing for *de novo* functions? Clarifying such fundamental questions may inform whether targeting classical downstream targets of PI3K-PIP_3_ signalling versus potential dependencies that manifest particularly when *PTEN* is lost, show therapeutic viability.

The spatial distribution of PIP species has been revealed by the use of domains of proteins that show preferential PIP affinity fused to fluorescent proteins as indirect reporters for PIP location^17–20^. For example, fusion to fluorescent proteins (e.g. GFP) of the pleckstrin homology (PH) domain from the cytohesin family of GTP exchange factors (GEFs) for ARF GTPases, such as ARNO/CYTH2, (e.g. GFP-PH-CYTH2) can be an exquisite sensor for PIP_3_ location. Splicing of these cytohesin PH domains alters their lipid specificity, wherein a di-glycine splice variant of the PH domain (PH-CYTH2^2G^) preferentially binds PIP_3_, while a tri-glycine splice variant (PH-CYTH2^3G^) associates with PI(4,5)_2_ ^21^. This illustrates how using lipid- preferential binding domains in such reporters allows detection of PIP distribution.

Although the PH domain of CYTH-type ARF GEFs have been extensively used as probes for PIP_3_ localisation, the extent to which they are required to enact PIP_3_ downstream signalling has mostly been neglected. Recent work identifies that the PIP_3_-specific variant of CYTH1 is required for signalling from c-Met to induce migration ^22^. Moreover, both PI4- and PI5-kinases are effectors of ARF GTPases themselves ^23–26^, highlighting that ARF GTPases are intimately involved in maintaining and effecting PIP homeostasis. ARF GTPases are evolutionarily conserved membrane trafficking regulators, controlling many aspects of cargo trafficking, such as turnover or recycling of receptor tyrosine kinases, cell-cell and cell-matrix adhesion proteins^27–30^. ARF GTPases are therefore well-placed to respond to changes in phospholipid metabolism that occur frequently in cancer and enact the cellular alterations that lead to invasive activity in cancer.

Here, we used the murine-derived model of HGSOC (ID8) to examine the cellular consequences of *Pten* loss on collective cancer cell behaviour using machine learning to detect phenotypic changes across multi-day time-lapse spheroid imaging. We identify that *Pten* loss induces PIP_3_-rich and -driven invasive protrusions into the extracellular matrix (ECM), which leads to invasive activity. We uncover that ARF6 is essential for this process. Through CRISPR-mediated ARF6 interactor screening, we identify that ARF6 acts in concert with the ARFGAP protein AGAP1 to promote recycling of active integrin in protrusions and drive invasion. Levels of this ARF6 module predict clinical outcome in ovarian cancer patients. Our approach therefore uncovers an ARF6 vulnerability upon PTEN loss in collective cancer cell behaviour in ovarian cancer.

## Results

### PTEN loss in the tumour epithelium and association with poor patient survival

To understand how *PTEN* expression levels are altered in ovarian cancer, we examined *PTEN* mRNA in tumour epithelium and stroma. In three independent datasets of laser capture-microdissected (LCMD) ovarian tumours separated into epithelium and stroma, *PTEN* mRNA was significantly decreased in Tumour versus Normal ovarian epithelium, whereas stromal *PTEN* levels were inconsistently altered (Fig 1A-C). Across bulk Ovarian Cancer tumour datasets, which include epithelium and stroma, three of six independent datasets show decreased *PTEN* mRNA in Tumour versus Normal samples, with non-significant datasets all possessing a low number of normal samples (n=4-6) (Fig. 1D). In The Cancer Genome Atlas (TCGA) Ovarian Cancer dataset, 73% of samples possessed *TP53* mutation and consequently *PTEN* alteration occurred frequently with *TP53* alteration, low *PTEN* mRNA was poorly associated with *PTEN* copy number changes and modestly associated with low PTEN protein levels (Fig. 1E). While comparing high levels of *PTEN* mRNA (Quartile 4, Q4) to lower (Q1+2+3) levels did not distinguish overall survival in ovarian cancer patients (Fig. 1F), an 11-month (p=0.0019) increase in survival was observed in high (Q4) versus not (Q1+2+3) PTEN protein levels (Fig. 1G). Accordingly, while low *PTEN* mRNA patients (Q1 versus Q4) display significant, but modest AKT activation (pT308, pS473) (Fig 1H), similar comparisons using PTEN protein levels revealed a significant and robust PI3K-AKT signalling signature in low PTEN protein patients (Fig. 1I). Therefore, *PTEN* loss in ovarian cancer, particularly at the protein level, occurs in the tumour epithelium, and is associated with upregulated AKT signalling and poor overall survival.

**Figure 1.**
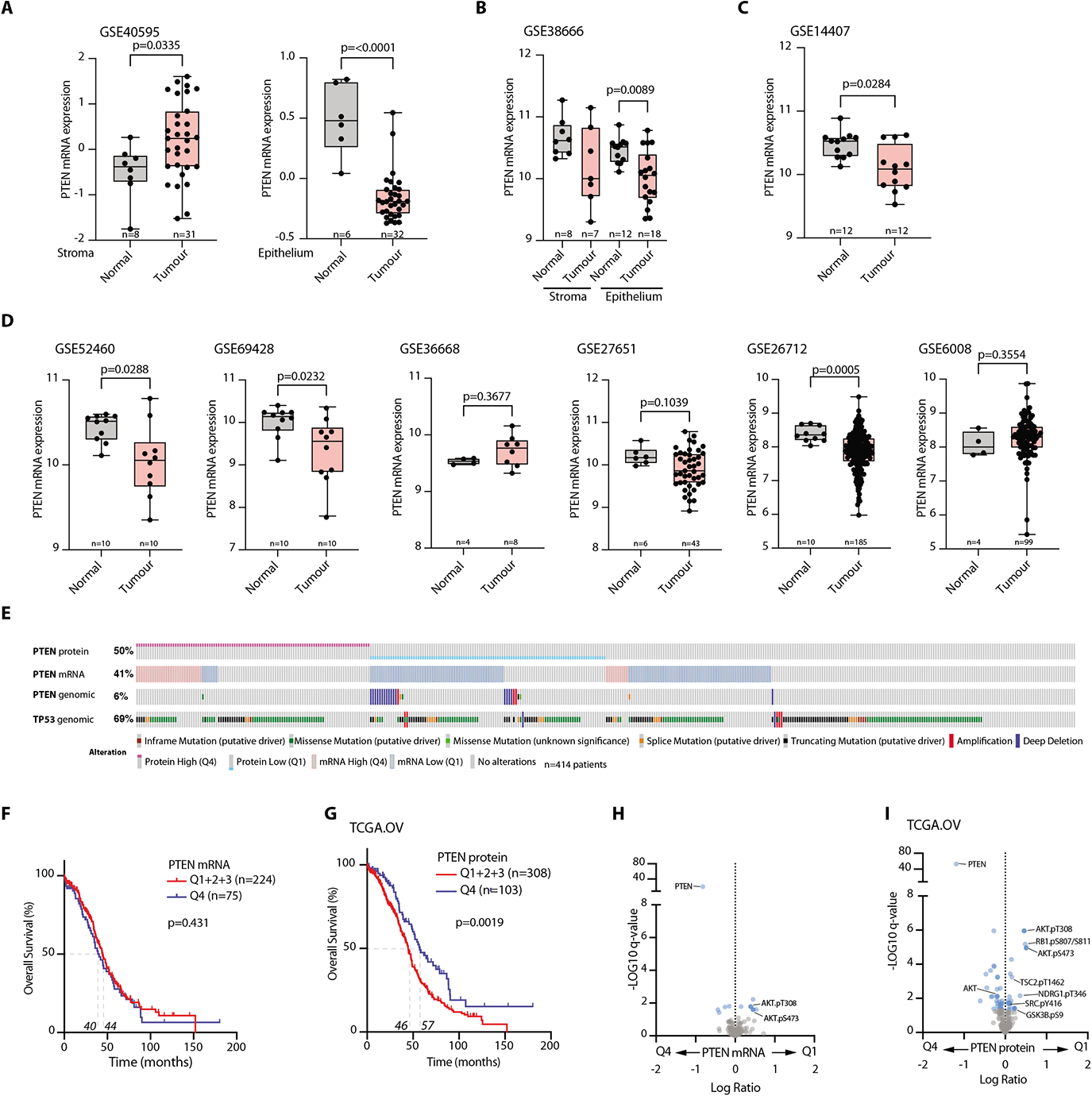
Loss of *Pten* in HGSOC epithelium is associated with poor outcome. **A-C.** *PTEN* mRNA levels in laser-capture micro-dissected normal ovarian surface epithelium versus high-grade serous ovarian cancer (HGSOC) epithelium or normal ovarian stroma versus ovarian cancer-associated stroma. Datasets; **(A)** GSE40595, **(B)** GSE38666, **(C)** Epithelium only, GSE14407. Sample size (n) and p-values, (Mann-Whitney) annotated. **D.** *PTEN* mRNA levels in normal ovarian surface epithelium versus tumour. Dataset ID, sample size (n) and p-values (Mann-Whitney) annotated. **E.** Copy Number, mRNA, protein level changes and mutations identified across *PTEN* and *TP53* in the TCGA dataset of OC. N=414 patients. **F.** Overall survival (% patients, months) of ovarian cancer patients. Highest quartile (Q4) versus combination of quartiles 1-3 (Q1+2+3), *PTEN* mRNA (TCGA, OV). Median survival (40 and 44 months), sample size (n) and p-value, Log-rank test (Mantel-Cox) annotated. **G.** Overall survival (% patients, months) of ovarian cancer patients. Highest quartile (Q4) versus combination of quartiles 1-3 (Q1+2+3), PTEN protein. Reverse Phase Protein Array Data, TCGA OV. Median survival (46 and 57 months), sample size (n) and p-value, Log-rank test (Mantel-Cox). **H.** Differential abundance (x, Log Ratio between conditions; y, -Log_10_ q-values) of Reverse Phase Protein Array data (TCGA, OV) in patient grouped by *PTEN* mRNA, High (Q4) vs Low (Q1). Significant, blue (-Log10 q-values >1.3); AKT signalling pathway, labelled. **I.** Differential abundance (x, Log Ratio between conditions; y, Log_10_ q-values) of proteins in PTEN High (Q4) vs PTEN Low (Q1) protein samples. Reverse Phase Protein Array Data, TCGA OV. Significantly altered components in AKT signalling pathway labelled (-Log10 q-value> 1.3).

### *Pten* loss induces modest effects in 2D culture

We aimed to model how *PTEN* loss in the epithelium affects tumour cell behaviour. A mutant *TP53* is a defining feature of HGSOC and is therefore an almost universal characteristic of the disease. Due to the numerous mutations that may occur on *TP53*, and the pleiotropic effects they may incur on the observed phenotype, we utilised ID8 ovarian cancer cells knocked out (KO) for *Trp53* and *Pten*, alone or in combination (Fig. S1A,B; including multiple clones of the double KO, dKO)^31, 32^. As a control, we made use of a Wild Type (WT) ID8 cell line, derived from Parental ID8 cells upon treatment with CRISPR plasmids containing the gRNA sequence that produced the *Trp53*^-/-^ subline but had in this specific case failed to introduce *Trp53* KO ^31^. *Pten* KO, either alone or in combination with *Trp53* KO, significantly increased AKT activation (pS473) (Fig. S1C-H), which occurred at the cell cortex in cells grown in 2-Dimensional (2D) contexts (Fig. S1I-J arrowheads). ID8 cells in 2D displayed a mixed morphology and could be classified into three categories: Cobblestone, Round, and Elongated (Fig. S1K-L). *Trp53* KO alone did not significantly affect cell shape class compared to parental (WT) cells, while *Pten* co-depletion decreased the Round phenotype, but did not cause a consistent increase in the other classes (Fig. S1M). Examination of proliferation or apoptosis, using puromycin treatment as a control for the latter, revealed that neither *Trp53^-/-^* or *Trp53^-/-^;Pten^-/-^* double KO (dKO) affected global growth or death in 2D (Fig S1N-O). Together, this suggests that p53 and PTEN loss do not manifest in major phenotypes in the examined conditions in 2D culture.

### PTEN loss induces ECM invasion

We next examined whether PTEN loss phenotypes may involve altered collective morphogenesis using multi-day time-lapse imaging of single cells plated into ECM gels that develop into 3-Dimensional spheroids (Fig 2A). While parental ID8 spheroids (WT) underwent proliferation and organisation into spherical multicellular objects with infrequent protrusive activity into the ECM (arrowheads), *Trp53^-/-^* spheroids exhibited modest protrusive activity. In contrast, *Trp53^-/-^;Pten^-/-^*spheroids displayed an enlarged, hyper-protrusive phenotype (Fig 2B, Supplementary Movie 1). This suggests that, in contrast to mild phenotypes in 2D culture, the phenotype of PTEN loss manifests in 3D contexts where ECM is present.

**Figure 2.**
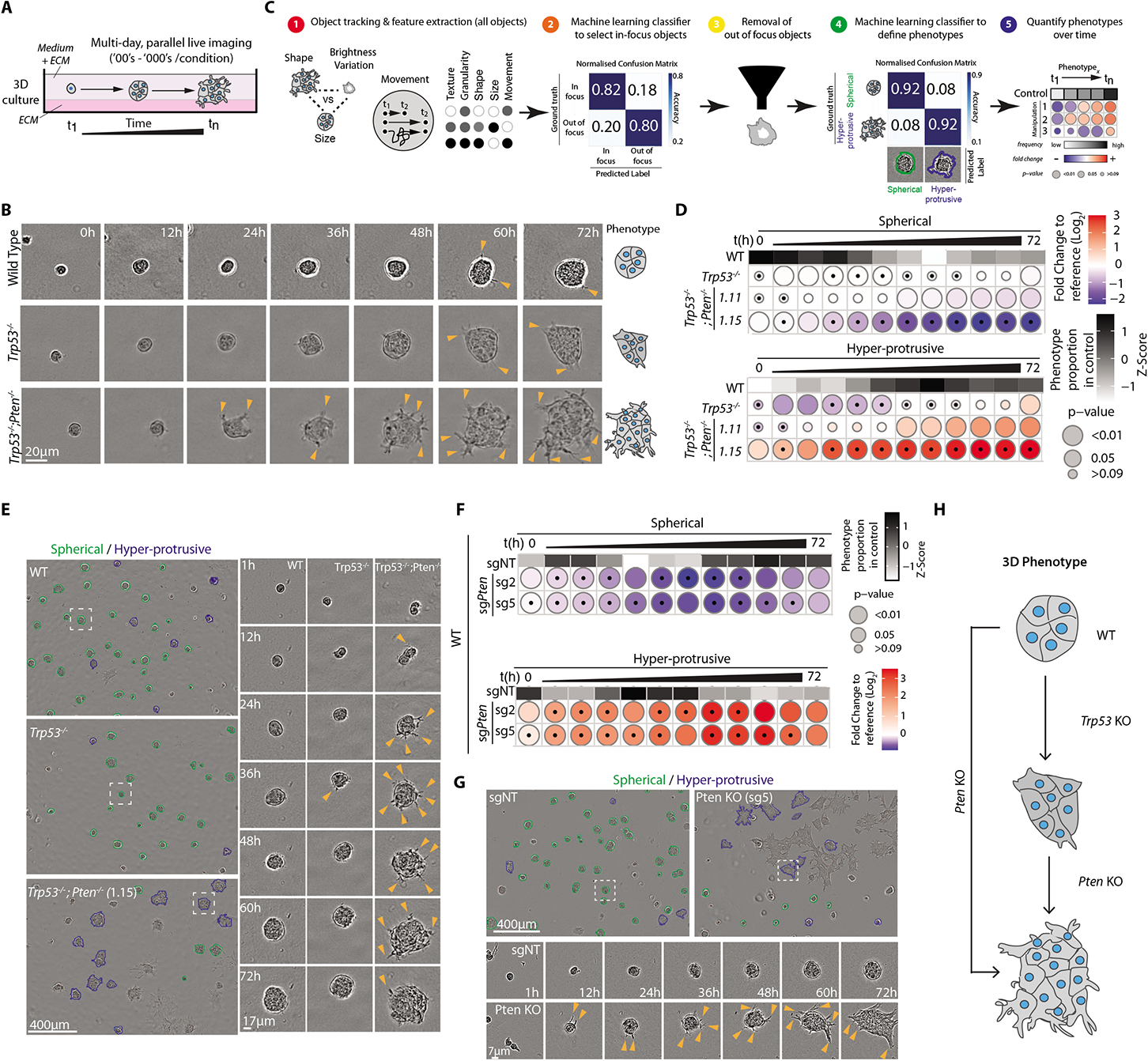
Loss of *Pten* is associated with collective invasion into ECM in a spheroid assay. **A.** Schema, imaging of ID8 spheroids in 3-Dimensional (3D) culture over time. Single cell suspensions were seeded onto and overlayed with ECM diluted in medium and then live-imaged. **B.** Time series, showing a representative spheroid for each genotype, 12 hr intervals. Arrowheads, protrusions into ECM. Scale bar, 20μm. Right, cartoon of phenotype. **C.** Schema, analysis pathway to classify ID8 3D phenotypes. (1) Phase contrast images were segmented using CellProfiler. Shape, size, movement, texture, granularity and brightness measurements were extracted for each object. (2) Machine learning was used to identify ’Out-of-focus’ objects, which (3) were filtered out of the dataset. (4) Additional machine learning was used to classify remaining ‘In-focus’ objects as ’Hyper-protrusive’ or ’Spherical’, with high accuracy. (5) Data analysis pipeline was used to quantify the log_2_ fold-change of each phenotype relative to control for each sub-line. **D.** Frequency, phenotype (Spherical, Hyper-protrusive) during 6 hr intervals, over 72 hr. N=3 experiments set up with repeated cultures of each sub-line, 3-5 technical replicates/subline in each experiment. **E.** Representative phase contrast images of indicated spheroid genotype from **(D)**, outlines pseudocoloured for classification (Hyper-protrusive, blue; Spherical, green) at indicated timepoints. Scale bars, 400μm or 17μm, as indicated. Magnified individual spheroids from boxed regions. Arrowheads, protrusions into ECM. **F.** Frequency, phenotype (Spherical, Hyper-protrusive) in ID8 parental spheroids expressing sgNT, sg2 *Pten* or sg5 *Pten*. N=3 experiments set up with repeated cultures of each sub-line, 2-6 technical replicates/subline in each experiment. **G.** Representative phase contrast images of indicated spheroid genotype from **(F)**, outlines pseudocoloured for classification (Hyper-protrusive, blue; Spherical, green) at indicated timepoints. Scale bars, 400μm or 17μm, as indicated. Magnified individual spheroids from boxed regions. Arrowheads, protrusions into ECM. **H.** Schema, phenotypes of ID8 spheroids with analysed genotypes. For **D** and **G**, Grayscale heatmaps, phenotype proportion (z-score) in control (Wild-type (WT)). Blue-to-red heatmap, log_2_ fold change from control. P-values, bubble size (Cochran-Mantel-Haenszel test with Bonferroni adjustment). Black dot, homogenous effect across independent experiments (Breslow-Day test, Bonferroni adjustment, non-significant p-value). Total spheroid number per sub-line, Supplementary Table 1.

To develop a quantitative measure of altered morphogenesis, we used a CellProfiler and CellProfiler Analyst-based *Fast Gentle Boosting* machine learning pipeline. Upon imaging, this pipeline could classify hundreds-to-thousands of spheroids per condition into Round and Hyper-protrusive ^33^. The steps involved were as follows: 1) phase contrast images of segmented spheroids were measured for texture, granularity, shape, size and movement features in tracked objects over multiple days, 2) a high-accuracy classifier was applied to determine in-focus objects, 3) out of focus objects were removed, 4) a second high-accuracy classification into Spherical and Hyper-protrusive spheroids was applied, and 5) the frequency of phenotypes over time across different manipulations were calculated (Fig. 2C). Application of this approach revealed that KO of *Pten*, whether alone or in combination with *Trp53* loss, and across multiple clones, results in induction of a hyper-protrusive, invasive spheroid phenotype (Fig. 2D-H, arrowheads; Fig. S2A, Supplementary Movie 2 and 3). Confirmation of this increased activity upon *Pten* KO was found in orthogonal invasion assays of wounded monolayers invading into ECM gels (Fig. S2B-D, Supplementary Movie 4), including increased invasion depth persistence (Fig. S2E). Notably, while invasion of parental cells occurred via infrequent chains of cells following a leader cell, upon *Pten* KO most cells at the leading edge displayed leader cell behaviours (Fig. S2C; Fig. S2F arrowheads; Fig. S2G). Therefore, loss of *Pten* is associated with desynchronised leader cell activity into the ECM, leading to a hyper-protrusive, invasive phenotype.

### *Pten* loss-induced invasion is associated with PIP_3_ enrichment at invasive protrusion tips

PI3-kinases (PI3Ks) act by adding a 3-phosphate group to PI(4,5)P_2_, generating PIP_3_. PTEN acts by removing this 3-phosphate group. We thus examined how *Pten* loss controls PI(4,5)P_2_ and PIP_3_ distribution in 3D contexts.

In poorly protrusive *Trp53^-/-^*spheroids, probes for PI(4,5)P_2_ (mNeonGreen [mNG]-tagged PH-PLC™1) and PIP_3_ (mNG-PH-CYTH3^2G^) localised cortically (red arrowheads), as well as in the nucleus in the case of PIP_3_ (yellow arrowheads) (Fig. 3A). In wounded invasive monolayers, PI(4,5)P_2_ and PIP_3_ localised similarly to parental cells, without an obvious enrichment at protrusion tips (Fig. S3A,B). However, in *Trp53^-/-^;Pten^-/-^*dKO cells, a pool of PIP_3_ was prominently located to the tips of protrusions in both spheroids and invasive monolayers (Fig. 3A, S3A,B; Red arrowheads, cell-ECM interface; Black arrowheads, cell-cell contacts; Green arrowheads, protrusion tips). This suggests that the elevated protrusive activity upon PTEN loss is associated with a pool of PIP_3_ at the tip of protrusions.

**Figure 3.**
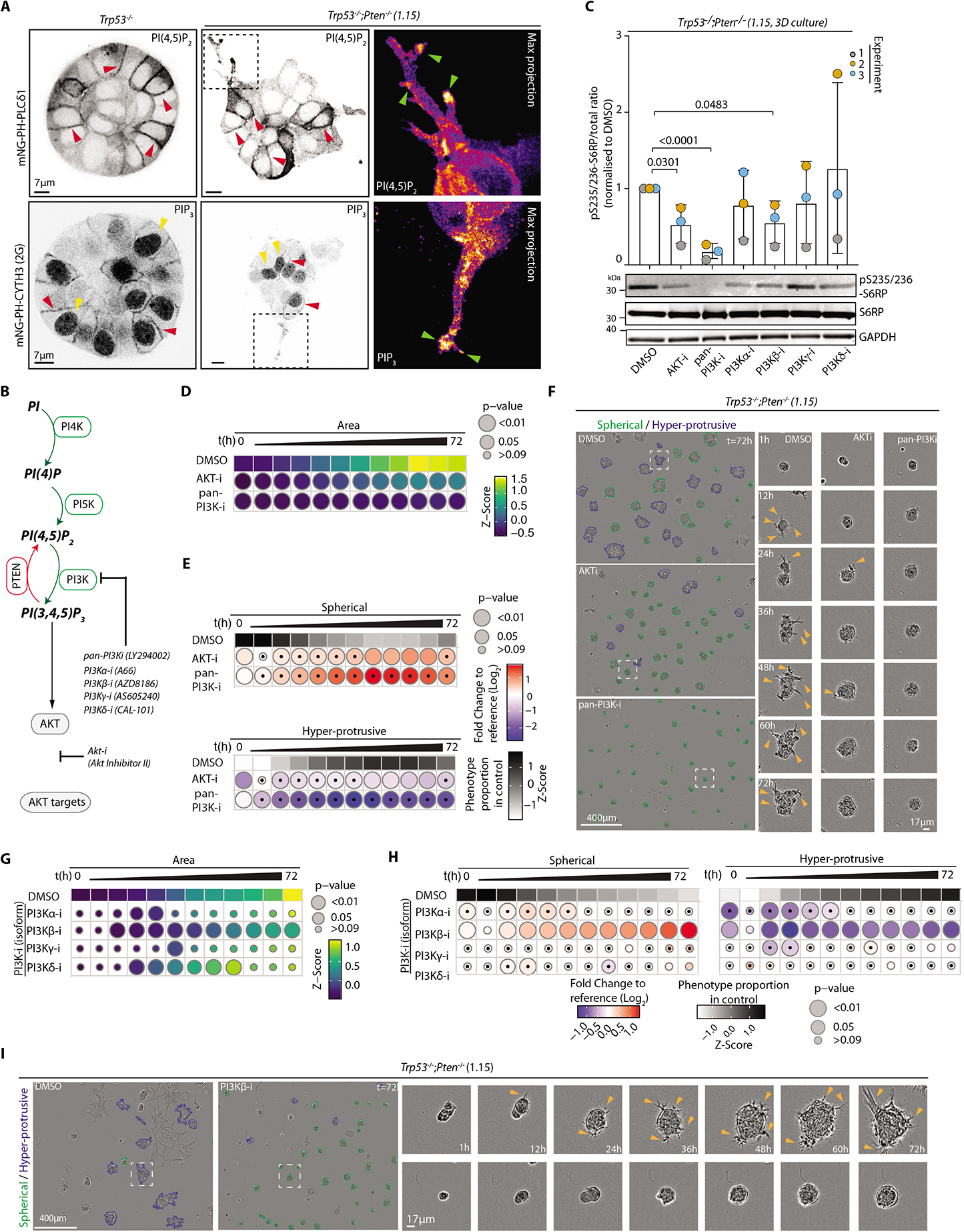
PI3K-AKT dependence of collective invasion. **A.** Confocal images of *Trp53^-/-^* or *Trp53^-/-^;Pten^-/-^* spheroids expressing mNeonGreen-tagged (mNG) biosensors for PI(4,5)P_2_ (PH-PLCδ1) or PIP_3_ (CYTH3^2G^/GRP1). Representative, 8 (*Trp53^-/-^*) or 10 (*Trp53^-/-^;Pten^-/-^*) spheroids imaged across n=2 experiments set up with repeated cultures of each sub-lines. Magnified images from boxed regions, pseudocoloured in FIRE LUT. Arrowheads: red, cell-cell contact; yellow, nucleus; green, protrusion tip. Scale bar, 7μm. **B.** Schema, select PI-kinases and phosphatases and their inhibitors participating in PIP_3_ production and downstream AKT phosphorylation. **C.** Western blotting and quantitation for S6RP pS235/236, S6RP, GAPDH (sample integrity control) in *Trp53^-/-^;Pten^-/-^*1.15 spheroids treated with DMSO or inhibitors annotated in **B** for 2 days. Representative of n=3 lysate preparations taken from spheroids set up with repeated culture of the cell line. Data, mean ± SD of pS235/236:total S6RP ratio, normalised to DMSO. P-values, unpaired, 2-tailed t tests, as annotated. **D-E.** Quantitation of **F**. **(E)** Heatmap (viridis colour scale), Area, mean of Z-score normalised values to control (DMSO). **(F)** Frequency, phenotypes (Spherical, Hyper- protrusive) over 72 hrs. Grayscale heatmaps, phenotype proportion (z-score) in control (DMSO). Heatmap, log_2_ fold change from control (blue to red). **F.** Representative phase contrast images, *Trp53*^-/-^;*Pten*^-/-^ 1.15 spheroids treated with DMSO, AKTi (AKT inhibitor II) or pan-PI3Ki (LY294002). Outlines: Hyper-protrusive, blue and Spherical, green. Magnified individual spheroids shown at select time points. Arrowheads, protrusions into ECM. Scale bar, 400μm or 17μm (indicated). N=2 experiments set up with repeated cultures of the sub-lines, 4-5 technical replicates/condition in each experiment. **G-H.** Quantification over time for **(G)** Area or **(H)** Phenotype frequency (Spherical, Hyper-protrusive) for ID8 *Trp53*^-/-^*;Pten*^-/-^ spheroids treated with PI3K isoform specific inhibitors: A66 (PI3Kα), AZD8186 (PI3Kβ), AS605240 (PI3Kγ) or CAL-101 (PI3Kδ). **(G)** Heatmap (viridis colour scale), mean, Z-score normalised values to control (DMSO). **(H)** Grayscale heatmap, phenotype proportion (z-score) in control (DMSO). Heatmap, log_2_ fold change to control (blue to red). N=2 experiments set up with repeated cultures of the sub-line, 3-5 technical replicates/condition in each experiment. **I.** Representative phase contrast images of ID8 *Trp53^-/-;^Pten^-/-^*1.15 spheroids upon treatment with DMSO or PI3Kβi as described in **G**. Outlines: Hyper-protrusive, blue; Spherical, green. Scale bar, 400μm or 17μm (indicated). Magnified spheroids shown for each condition at select time points. In **E-H**, P-values, bubble size (Area, Student’s t-test, Bonferroni adjustment; Classifications, Cochran-Mantel-Haenszel test with Bonferroni adjustment). Black dot, homogenous effect across independent experiments (Breslow-Day test, Bonferroni adjustment, non-significant p-value). Total spheroid number per condition, Supplementary Table 1.

As low PTEN protein patient tumours displayed a PI3K-AKT substrate phosphorylation activation signature (Fig 1I), we examined the requirement for PI3K- AKT signalling in the hyper-protrusive PTEN KO phenotype. PIP_3_ can be generated from PI(4,5)P_2_ through four Class-I PI3Ks (α,β,γ,δ). Pan inhibition of these PI3Ks (pan-PI3K-I; LY294002) or AKT (AKT-I; AKT Inhibitor II) (Fig. 3B, Supplementary Movie 5) abolished protrusion formation, resulting in smaller spheroids with upregulation of the Spherical phenotype and loss of Hyper-protrusive classification (Fig. 3C-F). Deconvolution of class-I PI3K contribution using isoform-preferential inhibitors revealed a major contribution of PI3Kβ to invasion and growth across the entire imaging period, and a more modest effect of PI3Kα at earlier timepoints (1- 36h) (Fig. 3B, G-I, Supplementary Movie 6). Therefore, in this system, and similar to Ovarian Cancer patients with low *PTEN* (Fig. 1I), loss of *Pten* is associated with PI3K-AKT signalling elevation, which is required for invasive protrusion formation and activity.

### The small GTPase ARF6 is required for *Pten* loss-mediated ECM invasion

We previously reported that in prostate cancer cells the small GTPases ARF5 and ARF6 are required to maintain invasive protrusion formation in 3D culture ^34^. In ID8 *Trp53^-/-^;Pten^-/^*^-^ dKO cells, stable lentiviral shRNA to *Arf5* or *Arf6* (Fig S4A-B) revealed a moderate effect of *Arf5* depletion on Hyper-protrusiveness and spheroid size (Fig. 4A-C), and no effect on invasion (Fig S4C-D). In contrast, *Arf6* stable depletion phenocopied PI3Kβ inhibition, resulting in reduced Area, a near complete loss of Hyper-protrusiveness in spheroids (Fig. 4A-C), and strongly attenuated invasion (Fig S4C-D, White arrowheads, invading cells, Supplementary Movie 7). *Arf5* or *Arf6* depletion did not affect AKT activation (pS473) (Fig S4A, B), suggesting that these GTPases act downstream of PIP_3_ generation. Validation of the *Arf6* depletion effect across five additional *Arf6*-targeting shRNAs revealed that the Hyper-protrusive activity of *Trp53^-/-^;Pten^-/^*^-^ dKO spheroids highly correlated with ARF6 levels (R2=0.7787, p=0.0199; Fig S4E-G). In *Trp53^-/-^*spheroids and tip cells of invasive monolayers, ARF6-mNG localised prominently at cell-cell contacts (white arrowheads), cell-ECM contacts (blue arrowheads), as well as to intracellular pools (yellow arrowheads) (Fig. 4D,E). In contrast, in *Trp53^-/-^;Pten^-/^*^-^ dKO spheroids and invading monolayers, while the cell-cell labelling (white arrowheads) of ARF6-mNG was still present, a new pool of ARF6-mNG could be observed at invasive protrusion tips (yellow arrowheads) (Fig 4D,E), mirroring PIP_3_ location upon *Pten* loss (Fig. 3A; Fig. S3A). Collectively, this suggests a role for ARF6 in regulating invasive protrusion tip formation upon PTEN loss.

**Figure 4.**
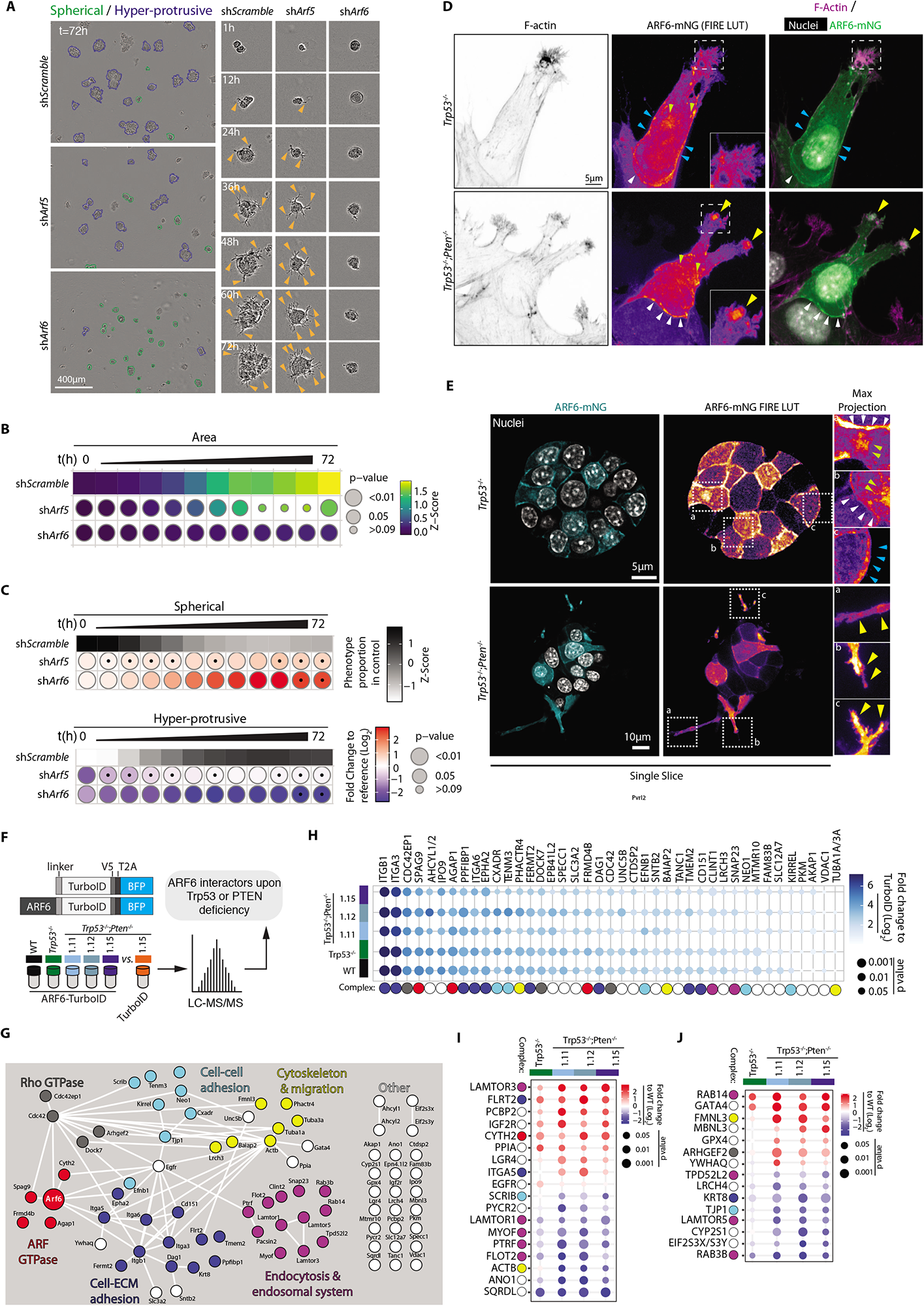
The small GTPase ARF6 is required for *Pten*-loss mediated ECM invasion. **A.** Representative phase contrast images, ID8 *Trp53*^-/-^*;Pten*^-/-^ 1.15 spheroids expressing shScramble, sh*Arf5* or sh*Arf6*. Outlines: Hyper-protrusive, blue; Spherical, green. Magnified spheroids shown at select time points. Arrowheads, protrusions into ECM. Representative of n=3 experiments set up with repeated cultures of each sub-line, 4-5 technical replicates/cell line in each experiment. Total initial number of spheroids in Supplementary Table 1. Scale bar, 400μm or 17μm (indicated). **B-C.** Quantitation of **A**. **(B)** Heatmap (viridis colour scale), area measurements, mean of Z-score normalised values to control (shScramble). **(C)** Frequency, phenotypes (Spherical, Hyper-protrusive). Grayscale heatmaps, phenotype proportion (z-score) in control (shScramble). Heatmap, log_2_ fold change from control (blue to red). P-values, size of bubble **(B),** Student’s t-test, Bonferroni adjustment; **(C)** Cochran-Mantel-Haenszel test with Bonferroni adjustment). Black dot indicates homogenous effect across independent experiments (Breslow-Day test, Bonferroni adjustment, non-significant p-value). **D-E.** Representative confocal images of ID8 *Trp53*^-/-^ and *Trp53*^-/-^*;Pten*^-/-^ 1.15 cells expressing mNeonGreen (mNG)-tagged ARF6 at **(D)** invasive front of wounded monolayer, stained with Hoechst and F-actin or **(E)** spheroids stained with Hoechst. Representative images from **(D)** n=2 experiments set up with repeated cultures of each sub-line, 3-7 fields imaged/sub-line in each experiment or **(E)** n=3 experiments set up with repeated cultures of each sub-line, 4-8 fields imaged/sub-line in each experiment. In **(E)**, boxed regions, magnified maximum projections of ∼10 Z-slices, pseudocoloured by FIRE LUT. Arrowheads: cell-cell contacts, white; cell-ECM contacts, blue; endosomes, grey; protrusion tips, yellow. Scale bar, 5μm. **F.** Schema, mass spectrometry (MS) proteomic-based TurboID approach for detecting ARF6-proximal proteins. **G.** STRING network analysis of interactions visualised using Cytoscape. Nodes manually annotated for known protein complexes. **H.** Heatmap, unchanging ARF6 interactors across genotypes. White to blue colour, ARF6 interaction score, Log_2_Fold Student’s t-test Difference in LFQ intensity compared to control ID8 *Trp53*^-/-^*;Pten*^-/-^ 1.15 TurboID alone. Interactors, sorted, descending order of mean interaction. Circle size, t-test p-value, coloured spots underneath denote the protein complex that each interactor belongs (in **J**), manual annotation. N=4 lysate preparations obtained from repeated cultures of each sub- line. **I-J.** Heatmaps of **(I)** Strong and **(J)** Weaker genotype-preferential interactors interactors (difference in LFQ intensity when compared to control ID8 *Trp53*^-/-^*;Pten*^-/-^ 1.15 TurboID). Coloured by Student’s t-test difference in LFQ intensity value for each protein when compared to ARF6-TurboID in ID8 WT cells. Interactors, sorted, descending order of mean interaction. Circle size, t-test p-value, coloured spots underneath denote the protein complex that each interactor belongs (in **G**), manual annotation. Experiment number described in **H**.

### Identification of ARF6-proximal protein networks

We examined how ARF6 is a vulnerability in *Pten*-null cells. We observed no consistent alteration in global levels of *Arf6* mRNA, protein or GTP-loading in *Trp53^-/-^ or Trp53^-/-^;Pten^-/^*^-^ cells compared to parental cell (WT), including multiple clones of the latter genotype (Fig S4H-J). We therefore examined whether, rather than ARF6 activation or levels being altered upon *Trp53* and *Pte*n loss, ARF6 interaction partners may change. We examined ARF6-proximal proteins through ARF6 fusion to the promiscuous biotin ligase TurboID ^35^ (Fig. 4F, Fig. S4K), in WT, *Trp53^-/-^*and *Trp53^-/-^;Pten^-/^*^-^ cells, including three clones of the latter genotype and across four independently repeated experiments. This allowed robust statistical support of identified ARF6-proximal proteins by mass spectrometry (MS) proteomic analysis. ARF6-TurboID mirrored ARF6-mNG, localising to cell-cell (black arrowheads) and cell-ECM contacts in 2D cells (Fig. S4L) and allowed rapid labelling of ARF6- proximal proteins upon biotin addition (Fig. S4M). Gene Ontology Cell Compartment (GOCC) analysis of ARF6-proximal proteins in *Trp53^-/-^;Pten^-/^*^-^ cells compared to TurboID alone in the same cells, identified significant enrichment for proteins involved in cell projections, filopodia, and ECM interactions (Fig. S4N). Cytoscape and STRING database analysis identified a highly interconnected network of ARF6- proximal proteins (Fig. 4G), including a singular ARF GEF, the PIP_3_-regulated CYTH2/ARNO protein, and a singular ARF GAP, AGAP1 ^36^. In addition, networks centred around proteins with known function of Rho GTPases, cell-ECM adhesion, cell-cell adhesion, endocytosis & endosomal system, and cytoskeleton & migration, as well as others with less reported connections.

We examined how ARF6-proximal proteins changed upon *Trp53* and *Pten* loss, dividing interactors into three categories: those that were largely unchanged across examined genotypes (Fig. 4H), strong interactors (Label-Free Quantitation (LFQ) intensity >1.2) altered in *Trp53^-/-^* and/or *Trp53^-/-^;Pten^-/^*^-^ cells compared to the parental (WT) genotype (Fig. 4I), and weak interactors (LFQ intensity<1.2) altered in *Trp53^-/-^* and/or *Trp53^-/-^;Pten^-/^*^-^ cells compared to the parental (WT) genotype (Fig. 4J, I). We include this third category, for example, to entertain interactors that may only bind in the *Trp53* and/or *Pten* loss conditions, but do not display significant binding in the WT condition.

The majority of prominent ARF6 interactors, such as β1-integrin/*Itgb1* and 〈3- integrin/*Itga3*, or AGAP1, did not change upon *Trp53* or *Pten* loss (Fig. 4H, colour scheme at bottom corresponds to grouping from 4G). When compared to WT ID8 cells, only a subset of ARF6 interactors were altered upon *Trp53* loss or when *Pten* was lost (Fig. 4H-I), such as CYTH2 interaction increasing upon *Trp53* loss irrespective of *Pten* status, or α5-integrin/ *Itga*5 interaction specifically induced upon *Pten* loss. This suggests that rather than large-scale alteration to ARF6 networks, loss of *Pten* may change a small number of key network members or render cells dependent on constitutive ARF6 network members.

### Cytohesin-2 function in invasion and contribution to ovarian cancer

The majority of known ARF GEFs were expressed in ID8 cells, and their expression was not consistently altered upon *Trp53* or *Pten* loss (Fig S5A). However, only a single ARF GEF, Cytohesin-2 (CYTH2), was identified as interacting with ARF6 (Fig. 4G-J). We therefore investigated chemical inhibition of Cytohesin-class GEFs using SecinH3 ^37^. SecinH3 treatment of *Trp53^-/-^;Pten^-/^*^-^ cells resulted in modestly smaller spheroids (Fig. S5B) that displayed less protrusive activity (Fig. S5C,D, arrowheads, Supplementary Movie 8). Accordingly, invasive activity (Fig. S5E, arrowheads), invasion distance, and multiple leader cell formation was strongly reduced upon SecinH3 treatment (Fig. S5F-G). This suggests that CYTH2 may function with ARF6 to regulate invasion.

In ovarian cancer patients, *CYTH2* mRNA was increased in the tumour compared to normal epithelium in both independent datasets of LCM tumours, whereas stromal *CYTH2* levels were inconsistent across datasets (Fig. S5H, I). In bulk tumour sequencing, five of seven datasets indicate increased *CYTH2* mRNA levels in tumours (Fig. S5J-P). Comparison of *CYTH2* mRNA levels based on median split comparing high (M2) versus low (M1) showed no significant different in survival (Fig. S5Q). *CYTH2* however, can be produced as two alternate transcripts based on alternate inclusion of exon 9.1, which encodes for a single additional glycine residue in the PH domain. Exclusion of exon 9.1 results in the CYTH2^2G^ isoform, which is preferential for PIP_3_ binding, whereas inclusion of exon 9.1 results in the PI(4,5)P_2_-binding CYTH2^3G^ isoform ^21, 38, 39^. Therefore, the exon 9.1 Percentage Spliced In (Ex9.1 PSI) ratio can be used to distinguish such alternate PIP-associating CYTH2 isoforms. A modest but significant (3-month, p=0.0262), decrease in overall survival was observed in patients displaying low Ex9.1 PSI (e.g. predominantly the PIP_3_-associating CYTH2^2G^ isoform) (Fig. S5R). Combination of *CYTH2* expression and splicing with *ARF6* expression levels revealed a significant decrease in overall survival in *ARF6*^HI^/*CYTH2*^HI^ patient subgroups (M2/M2) (Fig. S5S), and only when *CYTH2* Ex9.1 PSI was low (i.e. when PIP_3_-binding CYTH2^2G^ is predominant) (Fig. S5T). These data suggest that the PIP_3_-binding *CYTH2* isoform is associated with poor survival when co-expressed with high levels of *ARF6*.

### Identification of ARF6 interactors required for invasive activity

To identify ARF6 additional network proteins required for invasion, we performed a functional proteomic screen of 26 select interactors that represented constitutive ARF6 network members or those altered upon *Trp53* and *Pten* KO compared to WT (Fig. 5A). In this approach, *Trp53^-/-^;Pten^-/^*^-^ ID8 cells were transduced with a lentiviral pool of 5x sgRNAs/gene and Cas9, for each of the 26 interactors. Each transduced and selected cell pool was then plated as 3D cultures, also in an arrayed fashion (single gene per well). Machine learning classification of Spherical and Hyper- protrusive phenotypes was calculated from multi-day time-lapse imaging. To ensure accuracy of plating in 3D culture, sgRNAs were broken into four iterations containing distinct gene targets (Screen Iteration 1-4) and a control (sgNon-targeting, sgNT) per iteration (Fig. 5A). Each iteration contained multiple technical replicates of gene targets and controls, and was performed three independent times. The effect of each pooled sgRNA was calculated as fold-change to control classification.

**Figure 5.**
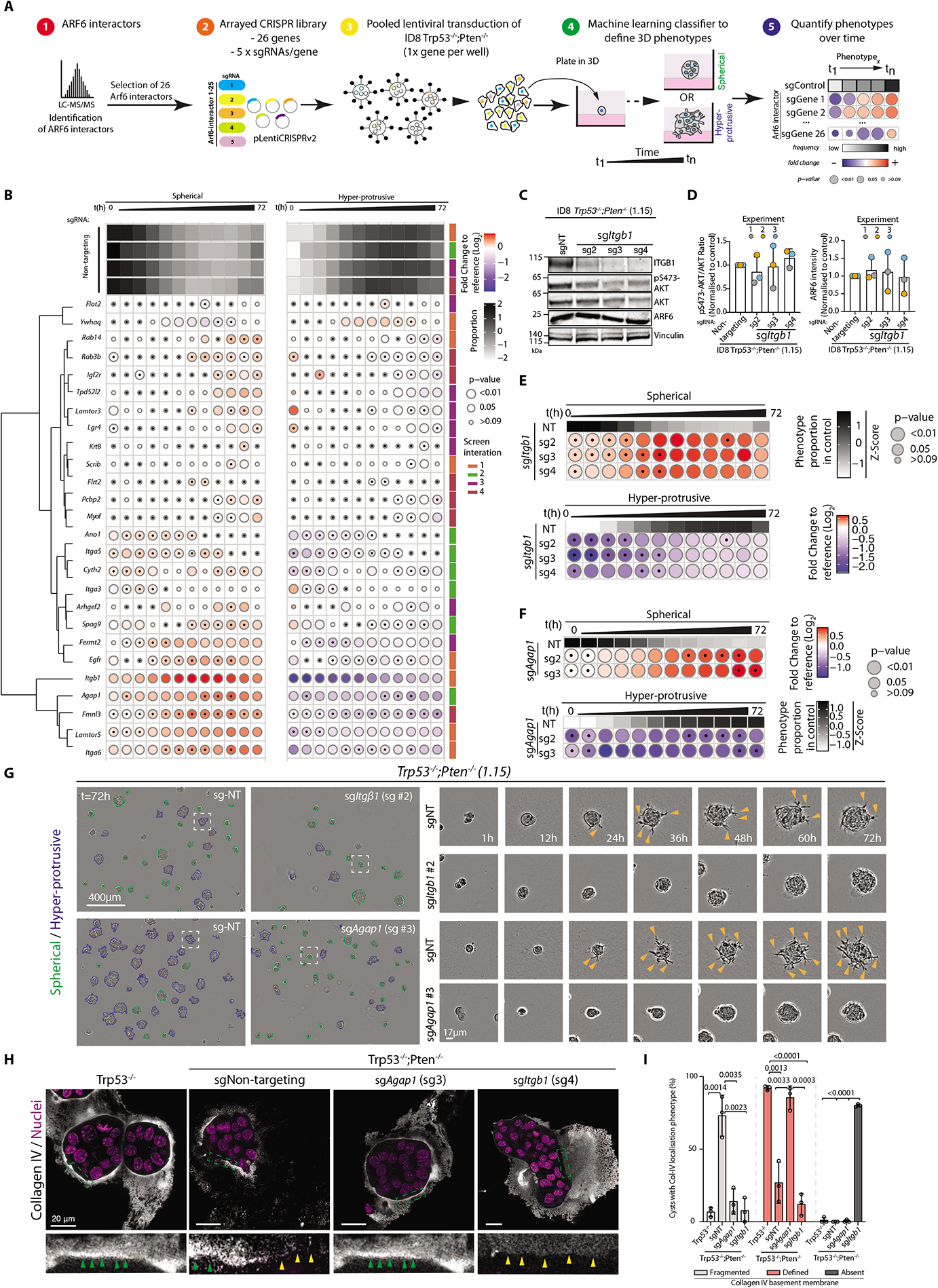
A functional proteomic CRISPR screen for ARF6-proximal proteins controlling collective invasion. **A.** Schema, CRISPR screen. 26 ARF6-proximal proteins from TurboID studies were investigated for their contribution to ARF6-mediated invasion of ID8 Trp53*^-/-^;Pten^-/-^* spheroids: (1) for each interactor, 5 sgRNAs were cloned into lentiviral CRISPR vectors (2). A pooled approach was used, generating a KO cell line with all 5 sgRNAs (3), live imaging performed (4) and the phenotype of each KO was compared to non-targeting sgRNA (5). **B.** Spheroid phenotype (Spherical, Hyper-protrusive) upon pooled gRNA CRISPR of indicated targets (sorted based on hierarchical clustering) in ID8 *Trp53^-/-^;Pten*^-/-^ clone 1.15 cells, performed in 4 parts (Iterations indicated). Grayscale heatmaps, phenotype proportion (z-score) in control per iteration (sgNT). Blue-to-red heatmap, log_2_ fold change from control. N=3-4 experiments per screen iteration set up using repeated cultures of each sub-line 3-6 technical replicates/subline per experiment. **C.** Western blot, β1-integrin (ITGB1), pS473-AKT, AKT, ARF6 from deconvolved ITGB1 sgRNA-expressing cells. VCL, loading control for ITGB1, sample integrity control for other blots. Representative blots of n=3 lysate preparations obtained from repeated cultures of each sub-line. **D.** Quantitation of **(C)**. Data, mean ± SD for pS473-AKT:total AKT band intensity ratio or ARF6 intensity, normalised to control (sgNT ID8 *Trp53^-/-^;Pten*^-/-^ clone 1.15) cells. P-values, unpaired, two-tailed t-test. **E.** Frequency, ID8 *Trp53^-/-^;Pten*^-/-^ clone 1.15 spheroid phenotypes (Spherical, Hyper- protrusive) upon CRISPR-mediated KO of *Itg*β*1*. Grayscale heatmaps, phenotype proportion (z-score) in control (NT). Blue-to-red heatmap, log_2_ fold change from control. N=3 experiments set up using repeated cultures of each cell line, 1-5 technical replicates/condition in each experiment. **F.** Frequency, ID8 *Trp53^-/-^;Pten*^-/-^ clone 1.15 spheroid phenotypes (Spherical, Hyper- protrusive) upon CRISPR-mediated KO of *Agap1*. Grayscale heatmaps, phenotype proportion (z-score) in control (NT). Heatmap, log_2_ fold change from control (blue to red). N=3 experiments set up using repeated cultures of each cell line, 1-5 technical replicates per sub-line in each experiment. **G.** Representative phase contrast images of ID8 *Trp53^-/-^;Pten*^-/-^ clone 1.15 spheroids treated with sgNT, *Itg*β*1*-targeting or *Agap1*-targeting sgRNA. Outlines; Hyper- protrusive, blue; Spherical, green. Magnified spheroids shown at select time points. Arrowheads, protrusions into ECM. Experiment number and sample sizes described in **(E)** and **(G)**. Scale, 400μm or 17μm (indicated). **H.** Representative confocal images of *Trp53^-/-^* and *Trp53^-/-^;Pten^-/-^ clone* 1.15 spheroids expressing sgNT, sg*Agap1* (sg3) or sg*Itgb1* (sg4), stained for collagen IV (grayscale) and Hoechst (magenta). Boxed areas, basement membrane region in higher magnification. Arrowheads, Collagen IV labelling that is: well-defined, green; fragmented/absent, yellow. Scale bar, 20μm. **I.** Quantitation of **(H)**. Collagen IV basement membrane staining as Defined, Fragmented, or Absent in spheroids set up across n=3 experiments set up using repeated cultures of each cell line, 1 technical replicate/experiment, 5-9 fields imaged per technical replicate, 365 spheroids scored in total. Data, mean ± SD of % of spheroids in each phenotype for independent experiments, with circles representing technical replicates. Unpaired t-test, p values annotated. **For B, E, G,** P-value, bubble size (Cochran-Mantel-Haenszel test with Bonferroni adjustment). Black dot, homogenous effect across independent experiments (Breslow-Day test, Bonferroni adjustment, non-significant p-value). Total initial number of spheroids analysed in Supplementary Table 1.

All pooled sgRNAs decreased Hyper-protrusiveness and increased Spherical phenotype to varying degree, except for 14-3-3theta/*Ywhaq*, which showed a modest increase in Hyper-protrusive activity (Fig. 5B). Notably, several constitutive ARF6 interactors (see Fig. 4G), such as ITGB1 and AGAP1, showed robust reduction in Hyper-protrusive activity when depleted (Fig 5B), while reduction in Hyper- protrusiveness could also be seen for sgRNAs against *Trp53* or *Pten* loss-induced interactors, such as *Cyth2* or *Itga5*, respectively (Fig. 4H-I, Fig. 5B).

Deconvolution of sgRNAs to *Itgb1*, *Agap1* and *Itga5* revealed efficient CRISPR editing to each target across multiple independent sgRNAs (Fig. 5C-D, Supplementary Movies 9 and 10; Fig. S6A-F). This occurred without affecting pS473-AKT levels, suggesting that these effects are downstream of PI3K signalling. Each of *Itgb1*, *Agap1* and *Itga5* depletions resulted in spheroids that lacked Hyper- protrusive activity (Fig. 5E-G, arrowheads; Fig. S6G-H), confirming the pooled screen results (Fig. 5B). This revealed that α5β1-integrin may be a major cargo of ARF6 that regulates interaction with the ECM to promote invasion, in conjunction with the GEF, CYTH2, and the GAP, AGAP1. This is particularly notable as although ITGB1 and AGAP1 association occurred across all genotypes (Fig. 4G), ARF6 association with ITGA5 increased specifically in *Pten*-null conditions (Fig. 4H). Notably, there was no change in the mRNA levels of either integrins in LCM HGSOC patient samples, while the comparison of either *ITGA5 or ITGB1* mRNA levels based on median split comparing high (M2) versus low (M1) showed no significant different in survival (Fig S6I-L). Consistently, neither *Itga5 or Itgb1* mRNA levels, or those of their ligand, Fibronectin *(Fn1*) changed across the ID8 sublines (Fig S6M).

To test whether altered interaction with the ECM underpins the *Pten*-null invasive phenotype we examined basement membrane formation around spheroids by staining for Collagen IV (COL4), the expression levels of which did not change upon loss of *Pten* (Fig S6M). The pattern of Collagen IV surrounding the ID8 spheroids could be classified as Fragmented, Defined, or Absent (Fig. 5H-I). In *Trp53^-/-^* spheroids Collagen IV staining was well-defined (92.16% of spheroids; green arrowheads). By contrast, in *Trp53^-/-^;Pten^-/^*^-^ spheroids (expressing a non-targeting sgRNA), the majority (73.1%) of spheroids displayed a fragmented basement membrane, representing clear regions of presence (green arrowheads) and absence (yellow arrowheads) of Collagen IV. Continuous basement membrane formation could be restored in *Trp53^-/-^;Pten^-/^*^-^ spheroids by KO of *Agap1* (85.5%). Notably, basement membrane was largely absent upon *Itgb1* KO (80,1%). This suggests that rather than a lack of basement membrane inducing invasion, fragmented basement membrane, providing asymmetric areas of cell-ECM adhesion, is associated with hyper-protrusive activity upon *Pten* loss. This asymmetry in basement membrane organisation requires the ARF6 interactor, AGAP1.

### AGAP1 regulates collective invasion and is associated with poor survival

Although the majority of ARF GAPs are co-expressed in ID8 cells, and this expression is unaltered across the examined genotypes (Fig. S7A), AGAP1 was the singular ARF GAP identified in the ARF6 interactome (Fig. 4J). *Agap1* isoforms can differ by alternate inclusion of Exon 14, encoding part of the PH domain (Fig. 6A), and resulting in AGAP1 Long (AGAP1-L) and AGAP1 Short (AGAP1-S) isoforms. The consequence of such splicing on AGAP1 is unknown.

**Figure 6.**
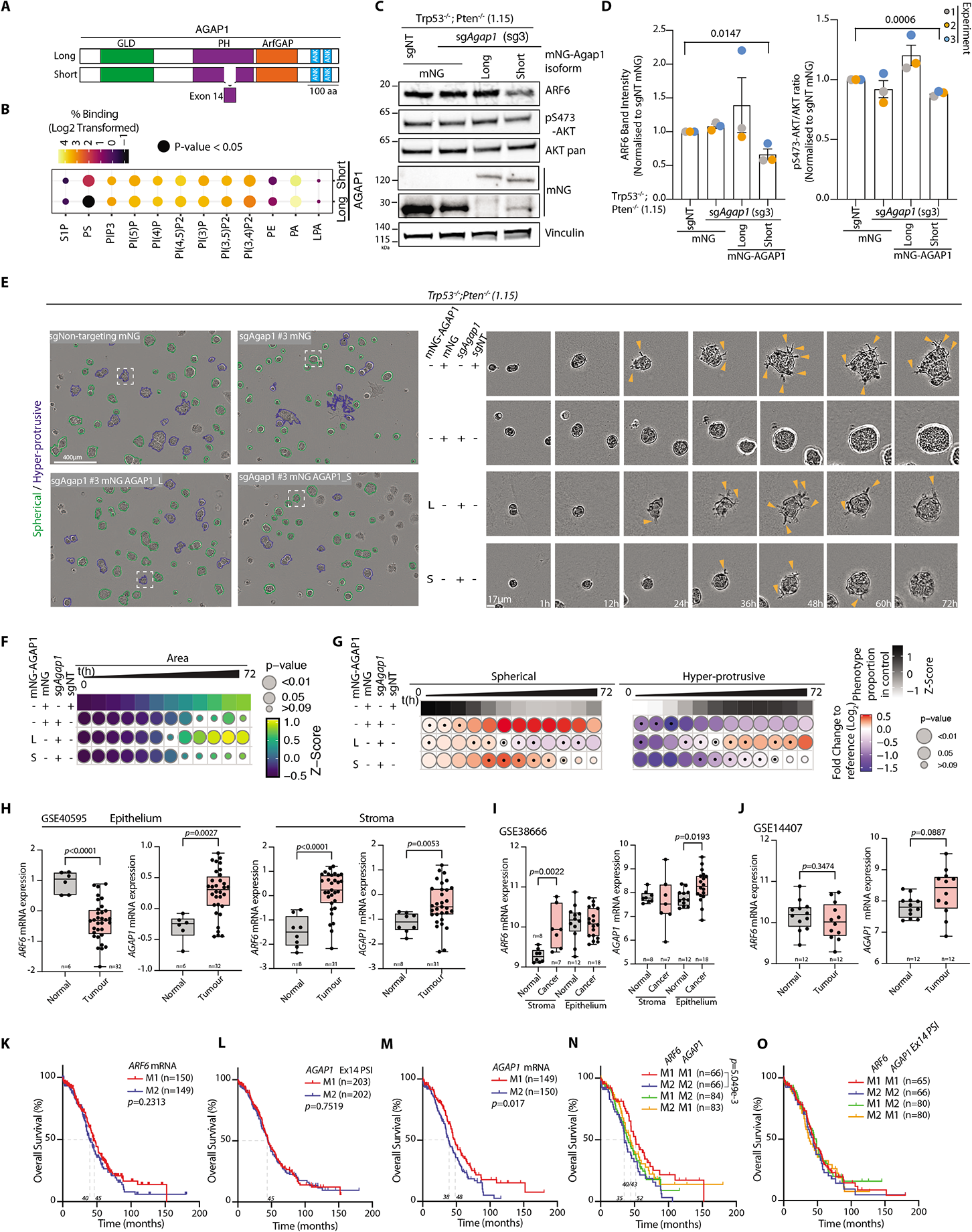
The ARFGAP AGAP1 controls invasion and stratifies survival. **A.** Schema, AGAP1 isoform domains. GLD, GTP binding-like domain; PH, Pleckstrin homology; ANK, Ankyrin; ARF GAP, ARF GTPase-activating Protein. Based on information found in www.ensembl.org67 ((’Long’ isoform, Transcript ID: ENST00000304032.13 for the human genome, and ENSMUST00000027521.15 for the mouse genome) or 804 amino acids (’Short’ isoform, Transcript ID: ENST00000336665.9 for the human and ENSMUST00000190096.7 for the mouse genome) and previously described annotations of AGAP1 domains^36^. **B.** Heatmap, differential association of isoforms with phospholipids. Data, Log2- transformed % of total signal between AGAP1-S versus AGAP1-L GST-tagged PH domain association with each phospholipid. P-value, circles size (unpaired t-test). n=3 blots per condition. **C.** Western blots of ID8 Trp53*^-/^-;Pten^-/-^* 1.15 cells expressing either sgNT or sgAgap1 (sg3) and either mNeonGreen (mNG) or CRISPR-resistant mNG-Agap1_S or -L isoforms. Blotted for ARF6, pS473-AKT, AKT, mNG, and VCL. VCL is loading control for AKT, pS473-AKT and ARF6 and sample integrity control for other blots. Representative blots of n=3 experiments using lysates taken from repeated cultures of the annotated sub-lines. **D.** Quantitation of **C**. Data, mean ± SD for ARF6 and pS473/AKT band intensity ratio, normalised to sgNT. P-values, unpaired two-tailed t-test, annotated when significant. **E.** Representative phase contrast images of ID8 *Trp53*^-/-^*;Pten*^-/-^ 1.15 spheroids treated with sgNT or AGAP1-targeting sg3 and expressing either mNG or mNG-fusion with either isoform of AGAP1. Outlines; Hyper-protrusive, blue and Spherical, green. Magnified spheroids shown at select time points. Arrowheads, protrusions into ECM. Scale bar, 400μm or 17μm (indicated). **F, G.** Quantitation of **(E).** Heatmap **(F)** (viridis colour scale), area, mean of Z-score normalised values to control (sgNT). **(G)** Frequency of ID8 spheroid phenotypes (Spherical, Hyper-protrusive). described in **E**. Grayscale heatmaps, phenotype proportion (z-score) in control (DMSO). Heatmap, log_2_ fold change from control (blue to red). P-values, bubble size **(F)**. Student’s t-test, Bonferroni adjustment. **(G)** Cochran-Mantel-Haenszel test with Bonferroni adjustment). Black dot indicates homogenous effect across independent experiments (Breslow-Day test, Bonferroni adjustment, non-significant p-value). n=3 experiments using repeated cultures of each sub-line, 5-6 technical replicates/sub-line in each experiment. Total number of spheroids analysed per condition in Supplementary Table 1. **H-J.** *ARF6 and AGAP1* mRNA levels in laser-capture micro-dissected normal ovarian surface epithelium versus HGSOC epithelium or normal ovarian stroma versus ovarian cancer-associated stroma. Specific datasets, sample size (n) and p-values (Mann-Whitney) annotated. **K-O.** Overall survival (% patients, months; TCGA OV dataset), of patients grouped by low (M1) versus high (M2) levels, based on a median split, of **(K)** *ARF6* mRNA, **(L)** *AGAP1* mRNA, **(M)** *AGAP1* Exon 14 percentage spliced in ratio (PSI), **(N)** combination of *ARF6* and *AGAP1* mRNA, or **(O)** combination of *ARF6* mRNA and *AGAP1* Ex14 PSI. Median survival, sample size (n) and p-value, Log-rank test (Mantel-Cox) annotated.

Association of purified recombinant AGAP1-L and AGAP1-S PH domains identified that the major difference in lipid binding between isoforms is in phosphatidylserine (PS) association, while broad binding to phosphoinositides and phosphatidic acid (PA) was indistinguishable (Fig. S7B-C; Fig. 6B). We performed reconstitution of sgRNA-resistant mNG-tagged AGAP1 isoforms into AGAP1 KO *Trp53^-/-^;Pten^-/^*^-^ cells (Fig. 6A, C), which were equally expressed (Fig. 6C, Fig S7D). These isoforms did not affect ARF6 levels or AKT activation (Fig. 6C,D). MNG- AGAP1-S-expressing spheroids were initially modestly smaller and strongly deficient in protrusive activity, but this was restored to control levels by later timepoints (Fig. 6E, F). In contrast, mNG-AGAP1-L-expressing spheroids showed increased size and while initially less Hyper-protrusive than control (sgNT) spheroids, AGAP1-L spheroids became more protrusive (arrows) in the second half of the imaging period (Fig. 6E-G). This suggests that both AGAP1 isoforms can support protrusive activity to varying degrees, though this is most prominent for the weakly PS-associating AGAP1-L isoform.

In five of seven bulk tumour datasets, *AGAP1* mRNA expression was elevated in tumour compared to normal ovarian tissue, which occurred in the epithelium in independent LCMD tumour datasets, but not consistently in the stroma. In contrast, *ARF6* showed a less consistent alteration across datasets, with *ARF6* mRNA elevated in only three of seven bulk tumour datasets, and *ARF6* mRNA elevation occurring in the stroma in LCMD datasets (Fig. 6H-J; Fig. S7E-J). While comparison of *ARF6* mRNA (Fig. 6K) or *AGAP1* Exon 14 PSI (Fig. 6L, High versus Low levels based on median split) did not affect overall survival, *AGAP1* mRNA levels strongly segregated survival groups, whether based on median split (Fig. 6M) or comparing Quartile 1 to Quartile 4 (Fig. S7K), in both cases a difference of 10- months survival. Combining *ARF6* and *AGAP1* mRNA levels, but not *AGAP1* Exon 14 PSI, even further separated overall survival, with a robust 17-month increase in overall survival of *ARF6^LO^-AGAP1^LO^*patients (red line), compared to the poor survival of *ARF6*^HI^-*AGAP1*^HI^ patients (blue line) (Fig. 6N). This same effect could not be found when examining splicing of *AGAP1* at Exon 14 (Fig. 6O). Together, these data indicate that AGAP1 is required for invasion in *Pten*-null cells, and that ovarian cancer patients with high *ARF6* and *AGAP1* levels, irrespective of the isoform of the *AGAP1*, have a poor clinical outlook.

### ARF6 regulates active integrin pools to produce invasive protrusions

Our data thus far indicate that a CYTH2-ARF6-AGAP1 module is required for invasion in *Pten*-null cells, and that α5-integrin and β1-integrin are two ARF6- promixal proteins essential for this phenotype. Although the mRNA levels of these two integrins are not altered in normal versus tumour epithelium, nor do they stratify patient survival based on a medium split (Fig. S6K-L), we explored whether the ARF6 module may act by regulating distribution of ECM-adhesion complexes to the tips of protrusions to drive invasion. Two markers of ECM-signalling hubs, pY397- FAK and pY416-Src family kinases (SFK), localised prominently to the tips of protrusions (Fig. 7A,B), in addition to the cell-ECM interface, suggesting active adhesion complexes localised to protrusion tips. In ovarian cancer patients with low levels of PTEN protein, pY416-SFKs protein levels were also elevated (Fig. 1I). Indeed, in Hyper-protrusive *Trp53^-/-^;Pten^-/^*^-^ cells the recycling of internalised total α5- or β1-integrin, the active form of β1-integrin, or a control cargo of Transferrin Receptor (TfnR), was increased at all timepoints examined compared to *Trp53^-/-^* cells, reaching statistical significance (p<0.05) at t=32min for active β1 integrin (Fig. 7C,E; Fig. S8A-C, G-I). *Arf6* depletion in *Trp53^-/-^;Pten^-/^*^-^ cells specifically blunted recycling of active integrin, but not total α5- or β1-integrin or TfnR (Fig. 7D,E; Fig.S8E-F, H-I). This indicates the CYTH2-ARF6-AGAP1 module specifically regulates active β1 -integrin recycling, whilst trafficking of inactive β1-integrins and TfnR is controlled by other signalling modules downstream of PIP_3_.

**Figure 7.**
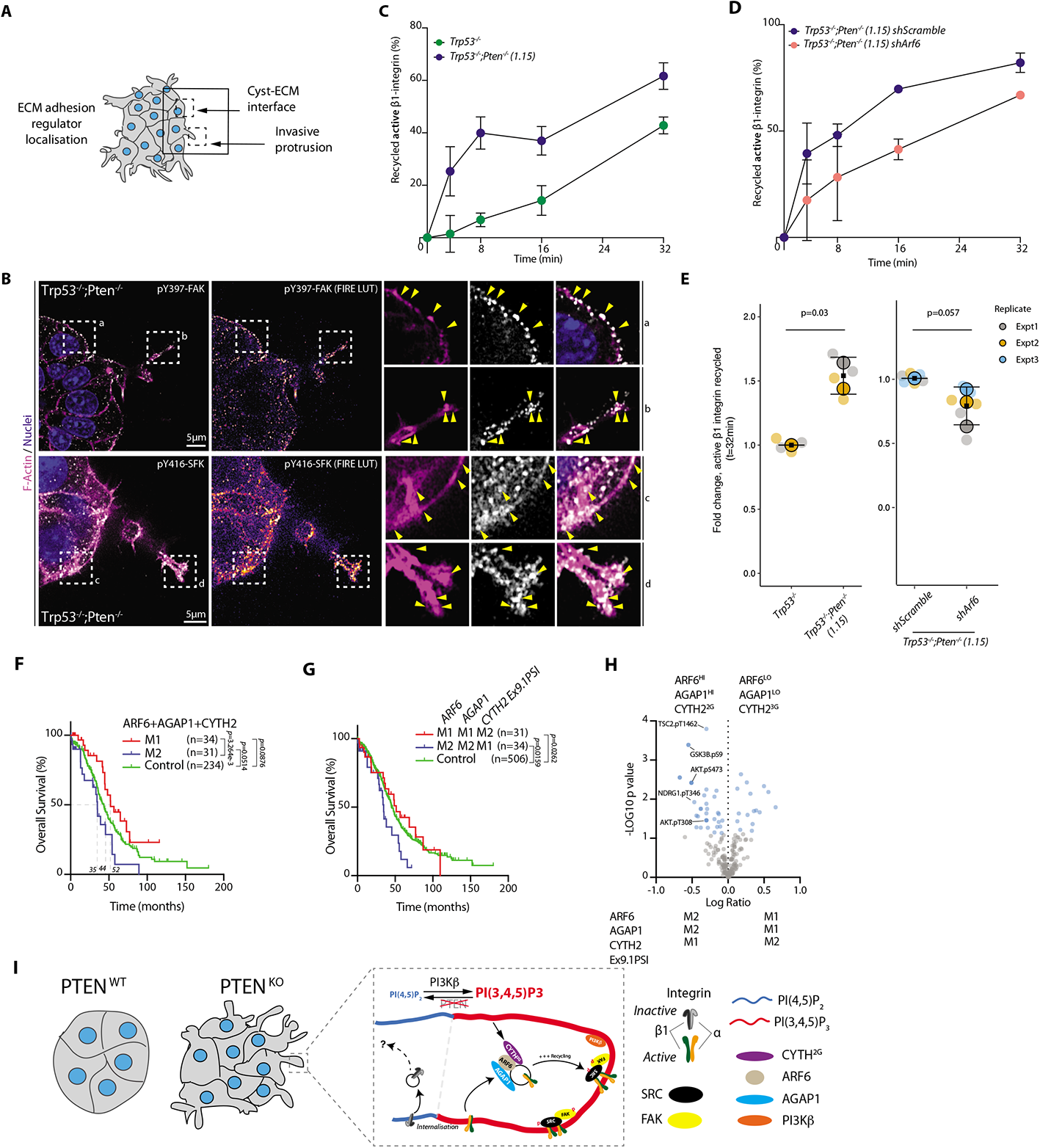
ARF6 controls invasion by regulating recycling of active integrins. **A.** Schema for interpretation of **(B)**, ID8 *Trp53*^-/-^*;Pten*^-/-^ spheroid interaction with the ECM. **B.** Immunofluorescence and confocal imaging of *Trp53^-/-^;Pten^-/-^*1.15 spheroids stained for pFAK (Y379) or pSRC Family Kinases (SFK pY416; grayscale), and with Hoechst (blue) and Phalloidin (F-actin, magenta). Scale bars = 5 μm. Representative of n=5 spheroids imaged per condition. Arrowheads, labelling. **C-E.** Representative capture ELISA graphs **(C,D)** and associated quantitation **(E)** for recycling of internalized cargoes between *Trp53^-/-^* versus *Trp53^-/-^;Pten^-/-^*cells or *Trp53*^-/-^*;Pten*^-/-^ cells expressing sh*Scramble* versus sh*Arf6* for active β1-integrin. Graphs shown are representative of n=2 **(C)** or n=3 **(D)** independent replicates. Data, mean (black square) ± SD for repeated experiments (large circles), 1-3 technical replicates per experiment per timepoint (smaller, semi-transparent circles), two-tailed t-test, p-values are annotated. **F-G.** Overall survival (% patients, months; TCGA OV dataset) of patients grouped into combined expression based on median mRNA split of: **(G)** (red line, M1) low for each of *ARF6, AGAP1, CYTH2*; (blue line, M1), each of *ARF6, AGAP1, CYTH2*; or (Green line), Control, remaining patients. For **(G)**, as for **(F)**, but *CYTH2* Ex9 PSI, rather than total CYTH2 levels. Median survival, sample size (n) and p-value, Log-rank test (Mantel-Cox) annotated. **H.** Differential abundance (x, Log Ratio between conditions; y, Log_10_ q-values) of proteins in PIP_3_-responsive module (ARF6^HI^-AGAP1^HI^-CYTH2^2G^) versus PI(4,5)P_2_- responsive ARF module (ARF6^HI^-AGAP1^HI^-CYTH2^3G^) protein samples. Reverse Phase Protein Array Data, TCGA OV. Significantly altered components in AKT signalling pathway labelled (-Log10 q-value> 1.3). **I.** Schema, molecular model for ARF GTPase regulation of integrin-dependent invasion.

Combined analysis of CYTH2-ARF6-AGAP1 module mRNA levels in ovarian cancer patients indicated that high levels of all three components (blue line; upper grouping based on median split of each gene’s expression, M2) showed a significant, 17-month decrease in survival compared to low levels (red line, M1) (p=3.264e-3; Fig. 7F). This effect could be recapitulated only when considering the PIP_3_-binding *CYTH2^2G^* isoform (i.e. low levels of Ex9.1PSI, blue line; p=0.0159, Fig. 7G). More modest effects could be observed in pairwise comparisons of *CYTH2- ARF6-AGAP1*, and splicing variants of *AGAP1* and *CYTH2* (Fig. S8J-L). A robust protein phosphorylation signature for PI3K-AKT signalling was observed in Ovarian Cancer patients with high levels of *ARF6-AGAP1* and PIP_3_-binding *CYTH2^2G^*, but not PI(4,5)P_2_-binding *CYTH2^3G^* (Fig. 7H). Collectively, this indicates a role for the potentially PIP_3_-responsive CYTH2-ARF6-AGAP1 module in regulating survival in ovarian cancer.

## Discussion

Here we propose a model of how loss of *Pten* can drive invasive behaviours, central to which is PTEN’s function as a phosphatase controlling PIP_3_ levels and localisation (Fig. 7I). In *Pten*-expressing cells, PIP_3_ localises to cell-cell contacts. In *Pten* KO cells, while cell-cell contact PIP_3_ is not lost, a prominent pool of PIP_3_ appears in ECM-invading protrusion tips. The small GTPase ARF6 likely acts directly in PIP_3_- elevated areas through activation by the PIP_3_-associating variant of its cognate GEF, Cyth2^2G^, and via its GAP AGAP1. This ARF6 module functions in the recycling of internalised active pools of integrin, thereby driving invasive protrusions enriched for the adhesion signal-transducing FAK and SFKs. This suggests a model wherein PTEN loss elevates recycling of invasion-promoting cargoes. The cellular consequence of this is altered interaction with the ECM.

It is notable that this CYTH2-ARF6-AGAP1 module was not specifically and only induced in *Pten*-null contexts, but rather that *Pten*-null cells became dependent on the module for enacting the invasive phenotype. Indeed, with the exception of α5- integrin, the majority of ARF6-proximal protein network was largely unchanged across *Trp53* or *Pten* knockout cells. This suggests that ARF6 and interactors likely have a steady-state function in recycling active integrins. It may be that this function is enhanced in *Pten* KO cells, as in our functional proteomic CRISPR screen of ARF6-proximal proteins we identified KINDLIN-2/FERMT2, a PIP_3_-binding regulator of integrin activation. When PIP_3_ levels are high it is possible that, in addition to selectively supporting recycling of previously internalised active integrin cargoes, ARF6 may collaborate with KINDLIN-2 to promote or maintain activation of recycled integrins at the plasma membrane, although this remains to be tested. In addition, a number of additional hits in the screen, such as EGFR, FMNL3, LAMTOR5 and ITGA6, gave strong reductions in Hyper-protrusiveness and may act as additional ARF6 cargoes or effectors in regulating collective invasion.

It is notable that the effects of *Pten* or *Trp53* loss were most prominent in 3D culture. This suggests that the phenotypes of loss of these central tumour suppressors may only manifest when cells are embedded in extracellular matrices and/or when multicellularity is considered. Indeed, the basement membrane around *Trp53^-/-^;Pten^-/^*^-^ spheroids were fragmented. This may explain how *Pten* loss resulted in the hyperactivation of leader-cell function in most cells at the ECM interface, rather than co-ordination of follower cells behind a singular leader cell. The tumour suppressor function of PTEN therefore may normally function to co-ordinate polarisation and cellular position in multicellularity by controlling basement membrane assembly through integrins, structurally influencing where invasive protrusions can occur.

The application of machine-learning approaches to live imaging allowed us to classify hundreds-to-thousands of spheroids tracked over time into robustly statistically supported categories, Spherical and Hyper-protrusive. While these labels were pivotal in identifying molecular perturbation that essentially turn on or off invasive behaviours, they are broad categories. It may be that subtle and important differences occur between perturbations, which could be further segregated with additional phenotype classifications. Indeed, while Hyper-protrusive *Trp53^-/-^;Pten^-/^*^-^ spheroids have fragmented basement membranes, this could be reversed to a fully defined basement membrane upon *Agap1* KO, thereby preventing protrusions. *Itg*β*1* KO spheroids, however, largely lacked an assembled basement membrane but also the ability to interact with the ECM to form protrusions. Both *Agap1* and *Itg*β*1* KO in *Trp53^-/-^;Pten^-/^*^-^ spheroids lack invasive protrusions, suggesting different alterations can result in similar morphogenetic consequences. More refined sub-categorisations may help to detect additional phenotypic variations.

*Pten* knockout alone was sufficient to drive *in vitro* invasion in the absence of *Trp53* depletion. The exact contribution of *Trp53* loss to the invasive phenotypes we examined is unclear. We observed modest alterations to phenotypes upon *Trp53* KO alone, such as increased protrusive tip formation or invasive capacity, but not sufficiently outside the range of normal variation to reach significance. Dissecting the role of *Trp53* is likely more complicated than we have examined as although *TP53* alteration is near-universal in ovarian cancer, many of these are distinct mutation, including some hotspots. Intraperitoneal injection of *Trp53*-null ID8 cells increases tumour growth rate and decreases survival compared to parental cells ^31^, showing that *Trp53* loss alone does cause *in vivo* functional differences to tumourigenesis. This tumourigenesis effect is accelerated *in vivo* by *Pten* co-knockout ^32^. Whether *Trp53* mutation versus loss differently contributes to *Pten*-depleted phenotypes remains to be examined.

The intraperitoneal injection of ID8 cells is an excellent system for *in vivo* examination of tumourigenesis in an immune-competent host. However, the rapid progression to clinical endpoint due to excess ascites production and spread of cells within the peritoneal fluid, rather than *bona fide* invasion, renders it poorly suited to determine contributions to metastasis, particularly in the case of *Pten*-null tumours due to rapid progression (*Trp53^-/-^;Pten^-/^*^-^, 34 days; *Trp53^-/-^*, 47 days; Parental ID8, ∼100 days) ^31, 32^. Validation of the *in vivo* contribution of PTEN loss to metastasis requires the use of approaches that allow metastasis to occur before clinical endpoint is reached. Introduction of such additional models is beyond the mechanistic cell biological studies provided here.

In ovarian cancer patient cohorts, *PTEN* loss is frequent and PTEN protein loss is associated with poor outcome. *ARF6* mRNA levels themselves were not consistently altered across independent datasets, making *ARF6* mRNA alone a likely unsuitable potential biomarker of poor outcome. Both the *CYTH2* GEF and *AGAP1* GAP mRNAs were elevated in tumour tissue in a number of datasets, however CYTH2 contribution is complicated by poor outcome being specifically conferred by the PIP_3_-associating Cyth2^2G^ isoform. Due to this isoform lacking a single amino acid difference to the PI(4,5)P_2_-binding CYTH2^3G^ isoform, this complexity renders CYTH2 analysis alone a poor biomarker candidate. *AGAP1* mRNA levels, in contrast, were a strong stratifier of outcome. Combined high vs low mRNA levels of *CYTH2-ARF6- AGAP1* provided the most robust 17-month different in survival of ovarian cancer patients, which occurred in patients with an PIK3-AKT signature. This emphasises the capacity of *in vitro* 3D cell biology to identify mechanistic insight into tumour suppressor contribution to cancer that can be used to clinically stratify poor and superior patient survival groups.

## Methods

### Cell Culture

All ID8 sublines were maintained in Dulbecco’s Modified Eagle Medium (DMEM) supplemented with 4% heat-inactivated Fetal Bovine Serum (FBS), 2 mM L-Glutamine, 1X Insulin-Transferrin-Selenium (0.01 mg/ml Insulin, 5.5 μg/ml Transferrin, 6.7 ng/ml Selenium) and 10U Penicillin-Streptomycin (all Gibco). HEK293-FT cells were maintained in DMEM with 10% FBS, 2 mM L-Glutamine and 0.1 mM Non-Essential Amino Acids (NEAA) (all Gibco). Cells were incubated at 37°C, 5% CO_2_ and routinely tested for mycoplasma contamination. Inhibitors were added at the following concentrations: 10 μΜ Nutlin3A (Sigma-Aldrich SML0580), 10 μM pan-PI3Ki (LY294002, Merck 440204), 200 nM PI3Kα-i (A66, Selleckchem S2636), 200 nm PI3Kβ-i (AZD8186, AstraZeneca), 200 nm PI3Kγ-i (AS605240, Stratech S1410), 200 nm PI3Kδ-i (Cal-101, Stratech S2226), 20 μM SecinH3 (Tocris 2849).

### Generation of stable cell lines

Lentiviral delivery systems were used for the generation of stable knock down (KD) lines (pLKO.1 puro), CRISPR knock out (KO) (pLentiCRISPR v2 Neo, Addgene #98292) or for overexpression of mNG protein fusions and the TurboID construct (pLX303/304 blast). A list of all sgRNAs is given in Supplementary Table 2, a list of shRNAs used is given in Supplementary Table 3. A list of all constructs is shown in Supplementary Table 4, and they will be made available on Addgene upon publication. HEK293-FT cells at 70% confluence were co-transfected with the plasmid of interest (0.50 μg DNA/ reaction) and lentiviral packaging vectors (pMD2.G Addgene plasmid #12259, 0.05 μg /reaction; and psPAX2, 0.50 μg /reaction Addgene #12260) using 6 μl Lipofectamine^TM^ 2000 (Thermo Fischer Scientific 11668019) in 500 μl Opti-MEM (Gibco)/reaction. Viral supernatants were centrifuged at 300 g for 5 min, filtered through 0.45 μm syringe filters (Starlab) and concentrated using 1/3 volume of Lenti-X concentrator (Clontech) as per the manufacturer’s instructions. ID8 cells were transduced with lentivirus for 3 days before selection (7.5 μg/ml blasticidin, 2.5 μg/ml puromycin, 1.5 mg/ml G418) or FACS sorting.

### Live 3D imaging and analysis

In a 96-well plate (Sigma-Aldrich Corning® 3595), 30μl of 50% Growth Factor Reduced (GFR) Matrigel (Corning® 354230) was used to precoat each well. The plate was centrifuged for 3 min at 1,500 g at 4°C. Cells were seeded as single-cell resuspensions supplemented with 2% GFRM (2000/well). The plate was centrifuged at 200 g for 2 min at room temperature (RT), incubated at 37°C for 4 hrs and scanned every hr for a total of 72 hrs using the Spheroid module of the Incucyte S3 system (Sartorius) and a x10 objective (1 field imaged/well). Images were extracted and aligned using the Fiji plugin “Image stabilizer” and a custom-made FIJI macro. Custom pipelines in CellProfiler (v4.2.0) identified and tracked individual spheroids at each time point, while extracting information on their size, shape, movement, and brightness variation. The generated dataset was used in CellProfiler Analyst (v2.2.0) to apply user-supervised machine learning (FastGentle Boosting algorithm) and classify spheroids as ‘Out of Focus’ or ‘In Focus’ (Accuracy >80% according to generated confusion matrix). The shape, size, and movement measurements of only ‘In Focus’ spheroids were used again in CellProfiler Analyst to construct rules (Supplementary Table 5) and classify them based on their morphology as ‘Hyper- protrusive’ or ‘Spherical’ (Accuracy 92% according to generated confusion matrix). These rules were exported as .txt files and incorporated in a CellProfiler pipeline that would perform prospective classification of new datasets without the need for retraining. A custom KNIME Data Analytics Platform (v3.3.1) pipeline was used to collate data, log2 transform and normalise the proportion of phenotypes across conditions and time points, perform statistical analyses and generate heatmaps. Statistical tests are described in figure legends and p values are annotated on figures. Heatmaps were generated using ggplot2 (v3.3.0)^40^ in the R environment (v3.6.2). Statistical comparison was performed in R using the Cochran-Mantel-Haenszel wherein a comparison is only statistically significant where the effect was present across all biological replicates. Using the DescTools (v0.99.31)^41^ R package, the Breslow-Day statistic was used to test the assumption that the magnitude of effect of a condition is homogeneous across all strata (biological replicates): a non-significant p-value indicates homogeneity. In both statistical tests, a Bonferroni adjustment was applied to correct for multiple testing.

### Cloning

Molecular cloning was performed using either classical ligation or In-Fusion technology. Restriction reactions were performed using High-Fidelity Restriction enzymes from New England Biolabs (NEB), by incubating 2 μg of DNA with 2U of each enzyme in the presence of 10X NEB CutSmart buffer, diluted to the appropriate concentration in nuclease-free water. The restriction reaction was performed at 37 °C (or other appropriate incubation temperatures) for 1 hr. The digested products were stained with 6X DNA loading dye and resolved at 110V for 1 hr in 1% agarose in TAE buffer supplemented with Midori green (Nippon Genetics, MG04). The desired DNA was purified using the QIAquick Gel Extraction Kit (28706X24) as per the manufacturer’s instructions. For ligations using the Rapid DNA Ligation Kit (Roche 11635379001) vector and insert were mixed in a 1:3 molar ratio, supplemented with 1x Dilution buffer, 1x Ligation buffer and 1 μl Ligase in a total volume of 10 μl, and incubated for 5 min at RT. For ligation reactions using the T4 DNA Ligase (NEB, M0202), the same molar ratio was used, supplemented with 2 μl of 10X T4 DNA Ligase buffer, 1 μl T4 DNA in a total of 20 μl. The reaction was performed at RT for 10 min and the ligase was subsequently heat-inactivated at 65°C for a further 10 min. For In-Fusion Cloning, a 1:3 vector to insert molar ratio was combined with 2 μl of 5X In-Fusion Reagent and brought to a total of 10 μl. In-Fusion reaction was performed at 50°C for 15 min. Bacterial transformation was performed using either Stbl3 (Invitrogen, C737303) or Stellar (Takara, 636766) chemically competent cells using the bacteria:DNA ratio as per the manufacturer’s instructions. A 10 min incubation on ice was followed by a heat-shocking step of 45 sec at 42°C. Transformed bacteria were plated on suitable agar plates and incubated overnight at 37°C.

### CRISPR-Knock Out (KO) Screen

All gRNAs used in the screen were generated against the mouse genome (Ensemble v.100) using an online tool (https://portals.broadinstitute.org/gpp/public/analysis-tools/sgrna-design). The top 5 sgRNAs (as determined by the sgRNA Designer tool) were further interrogated at the Integrated DNA Technologies (IDT) website (https://eu.idtdna.com/site/order/designtool/index/CRISPR_SEQUENCE). Only those with high on-target potential and low off-target risk were retained. To complete a set of 4-5 sgRNAs for each target, genes with less than 5 sgRNAs were further interrogated in the IDT website using the sgRNA Design tool and the best designs were added to the list of sgRNAs to be used. All sgRNAs were synthesized by IDT and ordered in a 96- deep well plate format as premixed oligo pairs (all sgRNA sequences available in Supplementary Table 2). The pLentiCRISPRv2 Neo vector was used as a backbone and the cloning procedure followed the steps as described by the Zhang Lab ^42, 43^. From each oligo pair, 2 μl were combined with 1 μl 10X T4 Ligation Buffer (NEB, M0202), 6.5 μl Nuclease-free H2O and 0.5 μl T4 Polynucleotide Kinase (PNK) (NEB, M0201). The oligos were annealed in a thermocycler with gradual T reduction from 95°C to 25°C at a rate of 5°C/min and subsequently diluted 1:20 into Nuclease free water (ThermoFisher Scientific AM9938). The pLentiCRISPRv2 plasmid was digested for 1 hr at 55°C with 1U per μg of DNA BsmBI-v2 (NEB, R0580), in 5 μl Buffer 3.1 and to a final volume of 50 μl. The digested backbone was dephosphorylated with 1U/mg FastAP Thermosensitive Alkaline Phosphatase (ThermoFischer Scientific, EF0651) for 10 min at 37°C. FastAP was inactivated at 75°C for 5 min. For ligation, 50 ng of digested plasmid were combined with 1 μl diluted oligo duplex, 1X Rapid DNA Ligation buffer (Rapid DNA Ligation Kit, Roche 11635379001), 1X Dilution buffer, nuclease-free water to a final volume of 10 μl and 1 μl Ligase. The mixture was incubated at RT for 5 min. Bacterial transformation was performed as described above.

The screen was performed in two phases. The first phase was performed in 4 iterations. The 4-5 gRNAs targeting each of the genes were pooled together and used to transduce transfect HEK-293F cells as described above. The produced viruses were used to transduce *Trp53*^-/-^;*Pten*^-/-^ 1.15 cells and generate a single, stable cell line (Pooled KO) for each gene. The Pooled KO cell lines were imaged with the Incucyte system as described above and compared to a Pooled sgNon-Targeting cell line. Processing of images and data analysis was performed independently for each iteration as described above. The results are presented as fold change to the iteration’s sgNon-Targeting cell line and each iteration has been colour-coded to allow for easier comparison. Heatmaps were generated using ggplot2 (v3.3.0)^40^ in the R environment (v3.6.2). Statistical comparison was performed in R using the Cochran-Mantel-Haenszel wherein a comparison is only statistically significant where the effect was present across all biological replicates. Using the DescTools (0.99.31?)^41^ R package, the Breslow-Day statistic was used to test the assumption that the magnitude of effect of a condition is homogeneous across all strata (biological replicates): a non-significant p-value indicates homogeneity. In both statistical tests, a Bonferroni adjustment was applied to correct for multiple testing. Select interactors were deconvoluted in phase 2, where 4-5 distinct KO cell lines were generated using each individual gRNA and compared against a single Non-targeting gRNA.

### Fixed 3D and 2D imaging and analysis

For 2D samples, ID8 cells were seeded on a black-bottom 96-well plate (Greiner 655090) with 2,000 cells per well and incubated for 24 hrs at 37°C. For 3D samples, ID8 spheroids were set up in 8-well chamber slides using 60 μl of 50% GFRM coating and 4,000 cells were seeded per well as single-cell resuspension supplemented with 2% GFRM and incubated for 48 hrs. Spheroids or cells were washed once with PBS and fixed with the addition of 4% PFA for 15 min at RT. Blocking was achieved with PFS (0.7% fish skin gelatine and 0.025% saponin in PBS). The following antibodies were added at 1:200 dilution in PFS and incubated overnight at 4°C with gentle shaking: Collagen IV (Abcam ab19808), pAKT pS473 (CST, 4060, D9E), pFAK pY397 (CST, 3283), pSRC Family pY416 (CST, 2101), V5-Tag (ABM, G189). Following three PFS washes, secondary antibodies Alexa Fluor® 488 Donkey Anti-Mouse IgG (H+L) and/or Alexa Fluor® 647 Donkey Anti- Mouse IgG (H+L) (Life Technologies A21202 and A31571, respectively) were added diluted in PFS (1:1,000) together with Alexa Fluor® 568 Phalloidin (Life Technologies A12380, 1:200 dilution), HCS CellMask™ Deep Red Stain (1:50,000) and Hoechst 34580 (Life Technologies H21486) (1:1000) and incubated at RT for 45 min. Samples were further washed with PFS (twice) and with PBS (thrice). Invading monolayers and spheroids were imaged using a Zeiss 880 Laser Scanning Microscope with Airyscan or at normal confocal function. Images taken in super resolution mode were processed using the Zeiss proprietary ZEN 3.2 software, exported as TIFF files and processed in Fiji. Invading monolayers and cells on 96-well plates were imaged using an Opera Phenix™ high content analysis system (x20 or x63) and the Columbus High-Content Imaging and Analysis Software (PerkinElmer, Version 2.9.1) was used to generate custom pipelines and perform object segmentation, intensity measurements and machine learning. For 2D morphology assays cells were identified based on nuclear staining (Hoechst) and the shape of each cell was defined by CellMask staining. Machine learning and manual training was used to classify cells as either ’elongated’, ’cobblestone’ or ’round’. Each cell was imaged in one plane. Cells in contact with the image border were discarded. For measurement of pAKT enrichment cells were identified based on nuclear staining (Hoechst), the total cell area was defined by CellMask staining and cells in contact with the image border were discarded. The cell area was split into 3 Regions: Ring Region, or ’Perinuclear’, resized to 35% Outer Border Shift (OBS) and 50% Inner Border Shift (IBS), ’Membrane’, resized to -10% OBS and 10% IBS and ’Cytoplasm’, resized to 10% OBS and 35% IBS. The cells were imaged in three planes with 1 μm distance between planes and processed as maximum projection. The staining intensity of pAKT was measured in each cell, for each individual area and was expressed as a proportion of the total (sum of all areas). The log_2_ transformed values were plotted using a custom R pipeline. Due to the large number of values measured, only the means of each experimental replicate are shown as dot-plots overlaid on violin plots depicting the distribution of the normalised pAKT intensity values of all cells measured. Statistical tests are described in figure legends and p values are annotated on figures.

### Invasion assay

Cell invasion was examined using the Scratch Wound assay method on the IncuCyte® System (Zoom 1 or S3, Satorius). The wells of a 96-well IncuCyte® Image Lock plate were coated with 20 μl of 1% GFR Matrigel (Corning® 354230) overnight and incubated at 37°C. The GFR Matrigel was removed and 6.5×10^4^ cells were added per well and incubated at 37°C for 4 hrs to facilitate attachment. The IncuCyte® Scratch Wound Tool was used as per the manufacturer’s instructions to create the wound. PBS was used to clear cell debris from the wells and 50 μl of 50% GFR Matrigel diluted in cell culture medium was placed on top of the cells followed by a 1-hr incubation at 37°C. If inhibitors/drugs were used; an appropriate volume was added in the GFR Matrigel to achieve the desired concentration. After incubation, 100 μl of cell culture medium (supplemented with inhibitors/drugs when required) were placed on top of the GFR Matrigel and the plate was imaged using the Scratch Wound module. Images were taken every 1 hr using the x10 objective and from a single field per well. Any wells where the wound did not form properly were not included in the analysis. Images were analysed using the dedicated IncuCyte® analysis tool. For each time point, the relative wound density (RWD) was measured. Statistical analyses were performed and graphs were generated using Microsoft Excel and RStudio (v1.4.1717). Data are presented for t=1/2 max of Control condition as bee swarm ’super-plots’ ^44^. Statistical tests are annotated on figure legends and p -values on figures. A similar approach was used for tracking of the leader cells, using the x20 objective. A 1:2,000 dilution of Incucyte® NucLight Red dye (Sartorius 4717) was added to stain nuclei and images were obtained every 15 min. Produced stacks were aligned using the Fiji plugin “Image stabilizer” and a custom FIJI macro. Leader cell tracking was performed using the MTrackJ plugin on FIJI. Spider plots were generated using RStudio (v1.4.1717). For scratch wound assays, fixed and stained for immunofluorescence, the same procedure was followed to set up cells as monolayers on black-bottom 96 well plates (Greiner 655090) and a 20 μl pipette tip was used to manually form the wound. Following an incubation period of 19 hrs at 37°C, the invading monolayers were fixed and stained as described above.

### RNA Extraction and sequencing

RNA extractions were performed using the RNeasy kit (QIAGEN 74106) and the QIAshredder spin columns (QIAGEN 79656). For 2D samples, cells at 70-80% confluence were washed twice with PBS and lysed in 600 μl of buffer RLT with 6 μl β- mercaptoethanol for 2 min. Cells were scraped and homogenised using a QIAshredder spin column centrifuged for 2 min at >8,000 x g. A 1:1 ratio of flow-through to 70% EtOH was mixed well and transferred onto a RNeasy Mini spin and the RNA isolated following the manufacturer’s instructions. The eluted RNA was stored at -80°C. For 3D spheroids, ID8 cells were passaged so they were sparse. The next day, 6-well plates (Falcon 353046) were coated using 180 μl of 50% GFR Matrigel per well and left to set for 60-75 min in an incubator at 37°C. Cells were washed, trypsinised, centrifuged, resuspended in fresh media, counted and adjusted to 8×10^4^ cells /ml. In each well, 1.6 ml of cell resuspension supplemented with 2% GFR Matrigel were added, and spheroids were allowed to develop for two days in a TC incubator at 37 °C with 5% CO_2_. For RNA extraction, cells were washed twice and the protocol for lysis was followed as described above for 2D samples. Adjustments were made to support the disruption of the ECM, by passing the lysates through a 25-27G needle slowly 10 times before homogenisation on the QiaShredder column. Lysis was performed using 350 μl of RLT buffer per well. For subsequent RNA sequencing of both 2D and 3D samples, extracted RNA underwent DNase treatment. An aliquot corresponding to 1.3 μg of RNA was obtained and combined with 1μl 10X DNase I Reaction Buffer and 1.3 μl DNase 1 (1U/μl) (ThermoFischer Scientific, 18068015) to a final volume of 10 μl with RNase-free water. The RNA/DNase mix was incubated at RT for 15 min and the reaction was stopped with addition of 10% v/v EDTA and heat-inactivation at 65°C for 10 min. The DNase treated RNA was placed on ice. 300 ng of RNA were taken and diluted to 50-100 ng/μl and used for TapeStation quality control of samples with a *R*NA *I*ntegrity *S*core (RIN) of >6 considered acceptable. The leftover 1μg of RNA was brought to 50μl volume with RNase-free water.

### RNA Sequencing and analysis

Sequencing was performed at the CRUK Beatson Institute using the Illumina polyAselection (2x#x00D7;36 PE Sequencing) kit without long reads. Quality checks and trimming on the raw fastq RNA-Seq data files were done using FastQC version 0.11.9^45^, FastP version 0.20.1 ^46^ and FastQ Screen version 0.14 ^47^. RNA-Seq paired-end reads were aligned to the GRCm38.101 version of the mouse genome and annotation ^48^, using HiSat2 version 2.2.1 ^49^ and sorted using Samtools version 1.7 ^50^. Aligned genes were identified using Feature Counts from the SubRead package version 2.0.1 ^51^. Expression levels were determined and statistically analysed using the R environment version 4.0.3^52^ and utilizing packages from the Bioconductor data analysis suite ^53^. Differential gene expression was analysed based on the negative binomial distribution using the DESeq2 package version 1.28.1^54^ and adaptive shrinkage using Ashr ^55^. Computational analysis was documented at each stage using MultiQC ^56^, Jupyter Notebooks ^57^ and R Notebooks ^58^. Log2 Transformation of counts and heatmap generation was performed using PRISM.

### Protein Domain-GST fusion purification

The PH domain sequences corresponding to the two isoforms of AGAP1 were ordered as GeneArt String DNA Fragments (ThermoFischer Scientific) and cloned by In-Fusion as GST Fusions in pGEX-4T1 vector. Plasmids encoding GST-Control, GST- hAgap1_PH_L and GST-hAgap1_PH_S were transformed into Rosetta 2(DE3)pLysS (Novagen) and proteins were expressed in Luria Broth based auto-induction medium including trace elements (Formedium) at 37°C for 6.5 hrs followed by 18°C for 12 hrs. Cells were harvested by centrifugation and the resulting pellets were resuspended in 200 mM NaCl, 50 mM Tris-HCl, pH 7.6, 1 mM DTT, 2 mM PMSF prior to lysis with a microfluidizer at ∼15,000 psi. Lysate was clarified by centrifugation and the clarified fraction was incubated with glutathione agarose resin (Agarose Bead Technologies), washed with resuspension buffer without PMSF, and eluted with wash buffer containing 10 mM glutathione and 5 mM DTT. The glutathione agarose eluate was diluted to a concentration of 50 mM NaCl, applied to a 5 mL HiFliQ Q ion exchange FPLC column (Neo Biotech) and eluted with a linear gradient ranging from 50 to 600 mM NaCl in 50 mM Tris-HCl, pH 8.5. Selected fractions were combined and applied to a HiLoad 26/60 Superdex 75 (manufactured by GE Healthcare, now produced by Cytiva Life Sciences) equilibrated in 150 mM NaCl, 25 mM Tris-HCl, pH 7.6, 1 mM DTT. Protein concentration was based on the measured absorbance at 280 nm and calculated molar extinction coefficients ^59^ of 44,350, 73,800 and 66,810 M-1 cm-1 for GST-control, GST- hAGAP1_PH_L and GST-hAGAP1_PH_S, respectively.

### BioID Mass Spectrometry proteomics and data analysis

An improved version of the promiscuous ligase BirA* (TurboID ^35^), was fused to the C-terminus of ARF6, followed by a V5 Tag, a cleavable T2A peptide and BFP and cloned into a lentiviral vector. The construct was stably expressed in ID8 cells as described above. A construct lacking ARF6 but containing BirA*, V5, T2A and BFP was used as a negative control for non-specific labelling. Cells at ∼70- 80% confluence were labelled for 30 min at 37°C by adding 50 μM of Biotin in full medium (Sigma-Aldrich S4501). Cells on Biotin-free medium were used as negative control. Cells were washed five times in ice-cold PBS and lysates were obtained by adding 800 μl of Lysis Buffer (50mM Tris-HCl pH 7.4, 100mM NaCl, 5mM in MS-grade water) supplemented with one each of cOmplete™, Mini Protease Inhibitor (Roche 05892970001) and PhosSTOP™ Phosphatase Inhibitor tablets (Roche 04906837001). The lysates were scrapped, left to incubate on ice for 30 min, sonicated and centrifuged at 13,600 g for 30 min at 4°C. Protein concentration was determined by performing a BCA assay (Pierce™ BCA Protein Assay Kit, Thermo Scientific 23225, used following manufacturer’s instructions). 350 μg of proteins were used per condition. 200 μL of streptavidin sepharose beads (Streptavidin Sepharose High Performance, Merck/Millipore GHC-17-5113-01) were washed thrice in 50mM Tris-HCl pH 7.4. All samples were incubated with 25 μL pre-washed beads in each at 4°C for 2 hrs with rotation. The beads were washed 4 times with 400 μL Washing Buffer (50mM Tris pH 7.4, 100mM NaCl, 5mM EDTA) and each time centrifuged at 1,200 g for 1 min at 4°C. Samples were resuspended in 2M urea in 100 mM ammonium bicarbonate buffer and stored at -20°C until further processing. On-bead digestion was performed from the supernatants. Quadruplicate biological replicates were digested with Lys-C (Alpha Laboratories) and trypsin (Promega) on beads as previously described ^60^. Following trypsin digestion, peptides were separated by means of nanoscale C18 reverse-phase Liquid Chromatography (LC) using an EASY-nLC II 1200 (Thermo Scientific) system directly coupled to a mass spectrometer (Orbitrap Fusion Lumos, Thermo Scientific). Elution was performed using a 50 cm fused silica emitter (New Objective) packed in-house with ReproSil-Pur C18-AQ, 1.9 μm resin (Dr Maisch GmbH). Separation was carried out using a 135 min binary gradient at flow rate of 300nl/min. The packed emitter was maintained at 50°C by means of a column oven (Sonation) integrated into the nanoelectrospray ion source (Thermo Scientific). Air contaminants signal levels were decreased using an Active Background Ion Reduction Device (ABIRD ESI Source Solutions). Data acquisition was performed using the Xcalibur software. A full scan was acquired over a mass range of 350-1400m/z at 60,000 resolution at 200 m/z. The 15 most intense ions underwent higher energy collisional dissociation fragmentation and the fragments generated were analysed in the Orbitrap (15,000 resolution). MaxQuant 1.6.14.0 was used for data processing. Data were processed with MaxQuant software ^61, 62^querying SwissProt (UniProt, 2019) Mus musculus (25198 entries). First and main searches were performed with precursor mass tolerances of 20 ppm and 4.5 ppm, respectively, and MS/MS tolerance of 20 ppm. The minimum peptide length was set to six amino acids and specificity for trypsin cleavage was required. Cysteine carbamidomethylation was set as fixed modification, whereas Methionine oxidation, Phosphorylation on Serine-Threonine-Tyrosine, and N-terminal acetylation were specified as variable modifications. The peptide, protein, and site false discovery rate (FDR) was set to 1 %. All MaxQuant outputs were analysed with Perseus software version 1.6.2.3 ^63^. Protein abundance was measured using label-free quantification (LFQ) intensities, which were calculated according to the label-free quantification algorithm available in MaxQuant ^64^, reported in the ProteinGroups.txt file. Only proteins quantified in all 3 replicates in at least one group were used for further analysis. Missing values were imputed separately for each column (width 0.3, down shift 1.8), and significantly enriched proteins were selected using a permutation-based t-test with FDR set at 5% and s0 = 0. Processed data were filtered using Microsoft Excel to select the hits likely representing true interactions. Typically, proteins with Student’s T-test Difference in their LFQ value of >1.2, when compared to ID8 Trp53-/-;Pten-/- 1.15 TurboID, were considered as true interactors. Protein networks were visualized using Cytoscape (v3.9.1) and bubble heatmaps were generated using RStudio (v1.4.1717).

### Integrin Recycling Assay

96-well ELISA plates were coated with 50 μl of integrin antibody at the optimised concentration diluted in 0.05M Na_2_CO_3_ pH 9.6 at 4°C overnight and blocked with 5% BSA in TBS-T. Cells at 80% confluence were washed with cold PBS and surface labelling was achieved with 0.13 mg/ml sulfo-NHS-SS-Biotin for 1 hr. For internalization, cells were washed with cold PBS and treated with 12-14°C cell medium for 30 min at 37 °C. Medium was removed, cells were washed with pH 8.6 buffer (50mM Tris pH7.5, 100 mM NaCl, adjust pH with 10 M NaOH) and MesNa (95 mM of MesNa in pH 8.6 Buffer) was added to achieve thiol reduction at 4°C for 30 min. Cells were washed with PBS and pre-warmed medium was added to induce recycling at 37°C and for the annotated time points. Cells were washed with PBS and pH 8.6 Buffer, followed by another round of thiol reduction. The reaction was quenched with the addition of 1 ml 20 mM iodoacetamide at 4°C. Cells were lysed using 280 μl of lysis buffer (200 mM NaCl, 75 mM Tris, 15 mM NaF, 1,5 mM Na3VO4, 7.5 mM EDTA, 7.5 mM EGTA, 1.5%

Triton X-100, 0.75% Igepal CA-630, 50 μg/ml leupeptin, 50 μg/ml aprotinin and 1mM AEBSF). Lysates were scraped and syringed once through a 30 G needle, span at 13,000 g for 10 min at 4°C, added in the ELISA 96-well plate and incubated overnight at 4°C . The ELISA plate was extensively washed with PBS-T to remove the unbound material. Streptavidin-conjugated horseradish peroxidase in PBS-T (1:6.666) containing 0.1% BSA was added to each well for 1 hr at 4°C. The plate was extensively washed with PBS-T and then with PBS to remove the Tween. For detection, 50 μl of Citrate/PO4 buffer (4 mM o-Phenylenediamine dihydrochloride corrected to pH 5.5 with H_2_O_2_) were added per well until colour in the total pool was well developed. The reaction was stopped with 50 μl of 8 M H_2_SO_4_ and absorbance was read at 490 nm.

### Proliferation and Cell Death assays

ID8 cells were plated in a 96-well plate (2,000/well), 24 hrs either alone or in the presence of 1:1000 dilution Sytox^TM^ Green (Invitrogen, S7020). Imaging was carried out on IncuCyte® ZOOM or S3 each hr for 48 hrs. Cell area (confluence) and the number of green objects over confluence were measured using the IncuCyte® analysis software.

### PCR Genotyping for AGAP1 KO cell lines

AGAP1 KO cell lines (ID8 *Trp53^-/-;^Pten^-/-^* 1.15 sg*Agap1*_2 and sg*Agap1*_3) and a control cell line (ID8 *Trp53^-/-;^Pten^-/-^*sgNT) were allowed to reach 80% confluence. Genomic DNA isolation was performed using 500 μl Lysis Buffer per well of 6-well plate (100mM Tris-HCl pH 8.5, 5mM EDTA, 0.2% SDS, 200 mM NaCl supplemented with 10 μg/ml Proteinase K) and overnight incubation at 55°C. Extraction was performed using Phenol:Chloroform:Isoamyl Alcohol (25:24:1 v/v) The upper layer was retained, and the extraction repeated twice on the supernatants using 450 μl and 400 μl of chloroform. The final 450 μl were precipitated by adding 35 μl of 4 M Sodium Acetate pH 5.2 and 770 μl of 100% EtOH. The precipitate was spun at 14000 rpm for 1 min, the supernatant removed and the DNA pellet washed twice with 1 ml of 70% EtOH. Following removal of EtOH, the DNA pellet was left to air-dry for 5 min, resuspended in 250 μl TE buffer and incubated at 55°C with gentle shaking for 5 min to dissolve. An empty pUC19 vector was linearised by PCR (pUC 5’: 5’-TCTAGAGGATCCCCGGGTAC-3’, pUC3’: 5-CTGCAGGCATGCAAGCTTGG-3’). The NCB1 Blastn tool was used to find the Mus musculus *Agap1* gene and identify a 500 bp region with the target sequence in the middle for each of the gRNAs. Primers that would amplify the specified genomic regions were designed, including 20 bp complementary edges to the linearised pUC19 backbone. PCR was performed using the Q5 ® Hot Start Master mix (NEB M0491), supplemented with 10 μΜ of each primer and 20 ng of template in a final volume of 25 μl in the following conditions: Initial denaturation: 98°C, 30 sec, denaturation: 98°C, 30 sec, annealing: 3-5°C lower than the Tm of the least stable primer in the reaction, 20 sec, extension: 72°C, 20s-30 sec per kb; repeat Steps1-3 for 30 cycle, final extension: 72°C, 2 min.

The PCR products were purified using a QIAquick PCR purification kit (QIAGEN 28104) and inserted into the linearised pUC19 backbone using the In-fusion reagent as described above. Single colonies were selected, the DNA was extracted (QIAGEN 27106X4) and the PCR products sequenced to identify CRISPR-derived Insertions or Deletions (INDELs). The PCR primers used to amplify the genomic DNA were the following:

sg*Agap1*_2: Forward: 5’-CGGGGATCCTCTAGAGCACAGGTAGAGCCTTGCAT-3’, Reverse: 5’-CTTGCATGCCTGCAGGTGGCAGATGTCTGTCTGAG-3’

sg*Agap1*_3: Forward: 5’- CGGGGATCCTCTAGATGCAGAGTTCAAATTTCAAG-3’, Reverse: 5’- CTTGCATGCCTGCAGGCTCACCCCCCTTTGCCACTC-3’

### Immunoblotting

For 2D samples, cells were plated for 48 hrs and allowed to reach 80% confluence. In the case of Nutlin3A treatment, the inhibitor was added 2 hrs before cell harvesting in fresh medium. For 3D samples, ID8 spheroids were generated in 6-well plates as described in the RNA extraction protocol, and inhibitors were added at the time of plating. Plates/wells were washed with ice cold PBS and lysed using RIPA Buffer (50mM Tris, 150mM NaCl, 1% NP-40 and 0. 25% Na deoxycholate with cOmplete protease inhibitor cocktail and PhosSTOP tablets). Cells were scraped and lysates incubated on ice for 15 min and clarified by centrifugation at 216×g at 4 °C for 15 min. For 3D samples the lysates were also passed through 25-27 G needle. BCA Protein Assay kit (Pierce) was used to determine protein concentration in 2D samples while a control immunoblot using samples of known concentration was used in the case of 3D samples. SDS–PAGE was performed in MES buffer at 160V for 1 hr, using 10 or 12-well Bolt™ 4- 12% Bis-Tris Plus Gels (Invitrogen NW04122BOX, NW0412BOX) and proteins were transferred to PVDF membranes using the iBlot 2 transfer system (Thermo Fisher Scientific). Membranes were incubated for 1 hr in Rockland blocking buffer (Rockland) and primary antibodies added overnight at 4 °C (1:1,000 unless stated otherwise). Antibodies used were: anti-2A peptide (Sigma-Aldrich, MABS2005, 3H4), anti-AKT pan (CST, 2920, 40D4), anti-ARF5 (Novus Biologicals, H0000281-M01, IB4), anti-ARF6 (Sigma Aldrich A5230), anti-GAPDH (CST 2118, 14C10, 1:5,000), anti-α5 integrin (Abcam, ab150361), anti-β5 integrin (Merck Sigma Aldrich, clone MB1.2), anti-AKT phospho S473 (CST 4060, D9E), anti-S6RP phospo S235/236 (CST, 2217, 5G10), anti- S6RP (CST, 2217, 5G10), PTEN (CST, 9552), anti-RFP (used to detect BFP) (Life Technologies R10367), anti-Streptavidin-Horseradish Peroxidase (HRP) Conjugated (Thermo Fischer Scientific SA10001), anti-TP53 (Abcam ab26, diluted in 5% milk in TBS-T), anti-V5 Tag (ABM G189), anti-Vinculin (Sigma Aldrich, V9131, 1:2,000). Secondary antibodies were added for 45 min, membranes were washed three times in TBST and imaged using a ChemiDoc Imager (BioRad) or Odyssey Imaging System (LI-COR Biosciences). Bands were quantified using Image Lab 6.1 (BioRad) or Image Studio Software 6.0 (LI-COR Biosciences). GAPDH or Vinculin were used as loading controls for each immunoblot (representative sample integrity controls are shown in the figures). Statistics were performed using 2-tailed t-test between a treatment and the control sample and all significant p values are annotated on figures.

### PIP strips

PIP strips (Tebu-bio 117P-6001) were used as per the manufacturer’s instructions. The membranes were blocked for 1 hr in PBS-T (0.1%) with 3% BSA, at RT. Each strip was incubated with 1 μg of purified GST or GST-PH Domain fusion in PBS-T with gentle agitation. Strips were washed thrice in PBS-T for 5 min. Anti-GST antibody (Sigma Aldrich, 06-332) was added diluted 1:1,000 in PBS-T with 3% BSA and incubated with gentle agitation in RT for 1 hr. The strips were washed thrice with PBS-T and secondary HRP-conjugated antibody was added (1:5,000 in PBS-T 3%) for 1 hr at RT. Supersignal™ West Pico Plus Chemiluminescent Substrate (Thermo Scientific 34580) was added for 3 min and the strips were scanned using the Bio-Rad ChemiDoc^TM^ Imaging system.

### Patient cohort analyses

Patient data were accessed, analysed and downloaded using in-platform tools from either cBioportal.org (The Cancer Genome Atlas, TCGA Ovarian Cancer Dataset) or the Gene Expression for Normal and Tumour database (GENT2, http://gent2.appex.kr/gent2/). Graphs and statistical analyses were generated in Prism 9 (GraphPad). Datasets can be accessed at cBioportal (TCGA, OV) or the Gene Expression Omnibus (GSE40595, GSE38666, GSE14407, GSE52460, GSE69428, GSE36668, GSE27651, GSE26712, GSE6008).

## Acknowledgements

This work was supported by the following grants: D.M.B. NIH K99CA163535, CRUK (C596/A19481) E.S., AR.-F., D. S. and L. McG., CRUK C596/A17196 and A31287. K.N. was supported by a Cancer Research UK Glasgow Centre PhD studentship (C7932/A25170). E.M.C. was supported by a University of Glasgow MVLS Doctoral Training Programme PhD Studentship. E.C.F. was supported by an industrial PhD studentship from Essen Bioscience and the University of Glasgow. S.Z and S.L were supported by CRUK (A29800 to S.Z.). L.B. and D.T.H, were supported by CRUK core funding to D.T.H. (A23278). M.N. R.S. and C.M were supported by CRUK core funding to C.M. (A29801). I.A.M. was supported by the Rivkin Center for Ovarian Cancer Research (574546) and Ovarian Cancer Action (P76567). J.C.N. was supported by CRUK Core funding (A28291).

We would like to thank the Core Services and Advanced Technologies at the Cancer Research UK Beatson Institute, with particular thanks to the Beatson Advanced Imaging Resource and Molecular Technologies. We would also like to thank Professor Stephen Tait, for provision of the pLenti CRISPR v2 Neo vector and Dr David Stevenson for provision of the pUC19 vector.

## Author Contributions

D.M.B conceived the project. Under supervision of D.M.B. lab work was performed by K.N., E.S., A.R.F., E.C. and J.C.N. E.C.F and L.Mc.G performed computational analyses under supervision of D.M.B. DS wrote a Fiji macro to aid live imaging analysis and MN wrote the R script which generates the spiderplots. Proteomic characterisation of ARF6 interactors was performed by S.L. under supervision of S.Z. Purification of recombinant protein domains was performed by L.B. under supervision of D.T.H. Analysis of the RNA-seq data was performed by R.S. under the supervision of C.M. The ID8 CRISPR series of cells, and guidance in experimental design was provided by I.A.M. The manuscript was written by K.N., E.S. and D.B. All authors read and commented on the manuscript.

## Data availability statement

**Figure S1.**
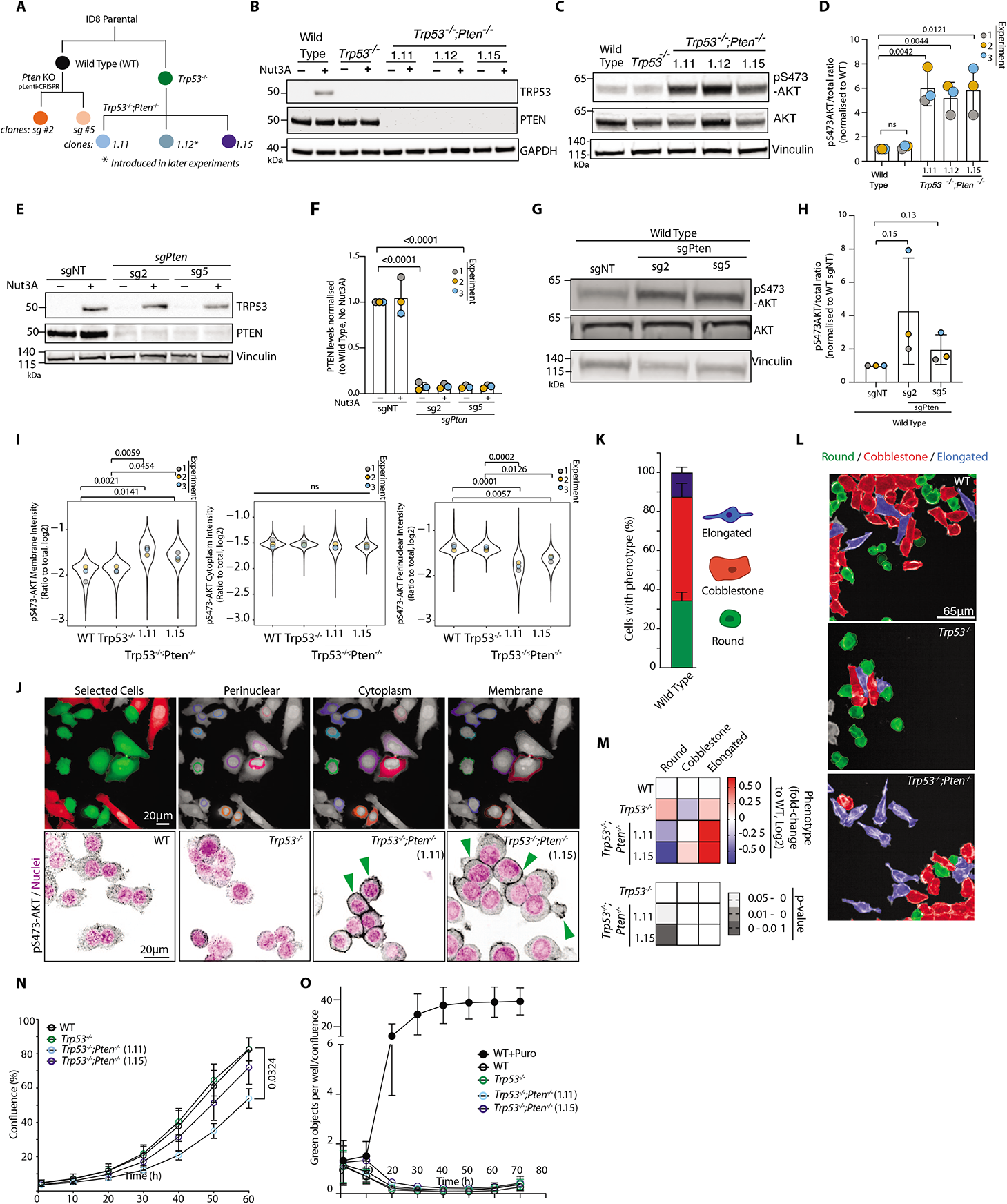
Characterisation of *Pten* loss effect on PI3K-AKT in 2D culture. **A.** Schema, derivation of *Pten* and *Trp53* alterations in ID8 sublines. **B-C.** Western blot, **(B)** TRP53, PTEN, GAPDH expression in ID8 sublines upon Nutlin-3A (MDM2 inhibitor) treatment to stabilise p53 or **(C)** pS473-AKT, AKT, Vinculin (VCL) in ID8 sublines. Each panel is representative of n=3 lysate preparations for each sub-line. GAPDH and VCL are loading controls for each panel. **D.** Quantitation of **C.** Data, mean ± SD of pS473-AKT/total AKT intensity ratio, normalised to WT. Unpaired, two-tailed t-test; p values, annotated. **E.** Western blot, TRP53, PTEN, VCL in ID8 Wild Type cells expressing non-targeting (sgNT) or *Pten*-targeting sgRNA upon Nutlin-3A (MDM2 inhibitor) treatment. Representative of n=3 lysate preparations for each sub-line. VCL is loading control. **F.** Quantitation of PTEN band intensity from **E**. Data, mean ± SD of band intensity, normalised to ID8 Wild Type sgNT. Unpaired, two-tailed t-test; p values, annotated. **G.** Western blot, pAKT(S473), AKT pan, VCL in ID8 Wild Type cells expressing non-targeting (sgNT) or *Pten*-targeting sgRNA. Representative of n=3 lysate preparations for each sub-line. VCL is loading control. **H.** Quantitation of **G,** p:t AKT ratio. Data, mean ± SD of band intensity, normalised to ID8 Wild Type sgNT. Unpaired, two-tailed t-test; p values, annotated. **I.** Quantitation of **J**. Data, ratio pS473-AKT signal at indicated regions to total area. Means, overlaid on plots of all data points as distinctly coloured dots according to culture replicate number. P-values are annotated, ANOVA with Tukey’s HSD test. **J.** ID8 cells plated in 2D, stained with pS473-AKT (Gray) and Hoechst (Magenta) (bottom panels), segmented into indicated regions (perinuclear, cytoplasmic, membrane) (Top panels). Colour in selected cells panel: red, excluded due to touching image edge, green, included for segmentation. N=3 experiments set up with repeated cultures of each sub-line, 4 technical replicates/subline in each experiment. Arrowheads, pS473-AKT at cell membrane. Scale bar, 20μm. **K.** Percentage of cells classified as Round (green), Cobblestone (red) or Elongated (blue) in Wild Type ID8 cells. N= 2 experiments set up with repeated cultures of each sub-line, 4 technical replicates/subline in each experiment. **L.** Representative images, cells classified by shape (Round, green; Cobblestone, red; Elongated, blue) in ID8 WT, *Trp53^-/-^* and *Trp53^-/^-;Pten^-/-^ 1.15* cells grown in 2D. Scale bar, 65μm. **M.** Heatmap, log_2_ fold change, mean proportion across indicated lines. Grayscale heatmap, p-values for each comparison. N=2 experiments set up with repeated cultures of each sub-line, 4 technical replicates/subline in each experiment. P-values, Ordinary one-way ANOVA with multiple comparisons to WT. **N.** Proliferation assay based on well confluence over time. N=3 experiments set up with repeated cultures of each sub-line, 4-5 technical replicates/subline in each experiment. Data are presented as mean ± SD. Unpaired, two-tailed t-test between WT and each of the sublines per time point. Significant p-values annotated. **O.** Cell death assay, green object (Sytox green fluorescence) confluence over time. N=2 experiments set up with repeated cultures of each sub-line, 4 technical replicates/subline in each experiment. Kruskal-Wallis ANOVA was performed at t=10 h and t=20 h, all comparisons are non-significant (p-value >0.05). In **H, I, J**, total number of cells analysed per condition in Supplementary Table 1.

**Figure S2.**
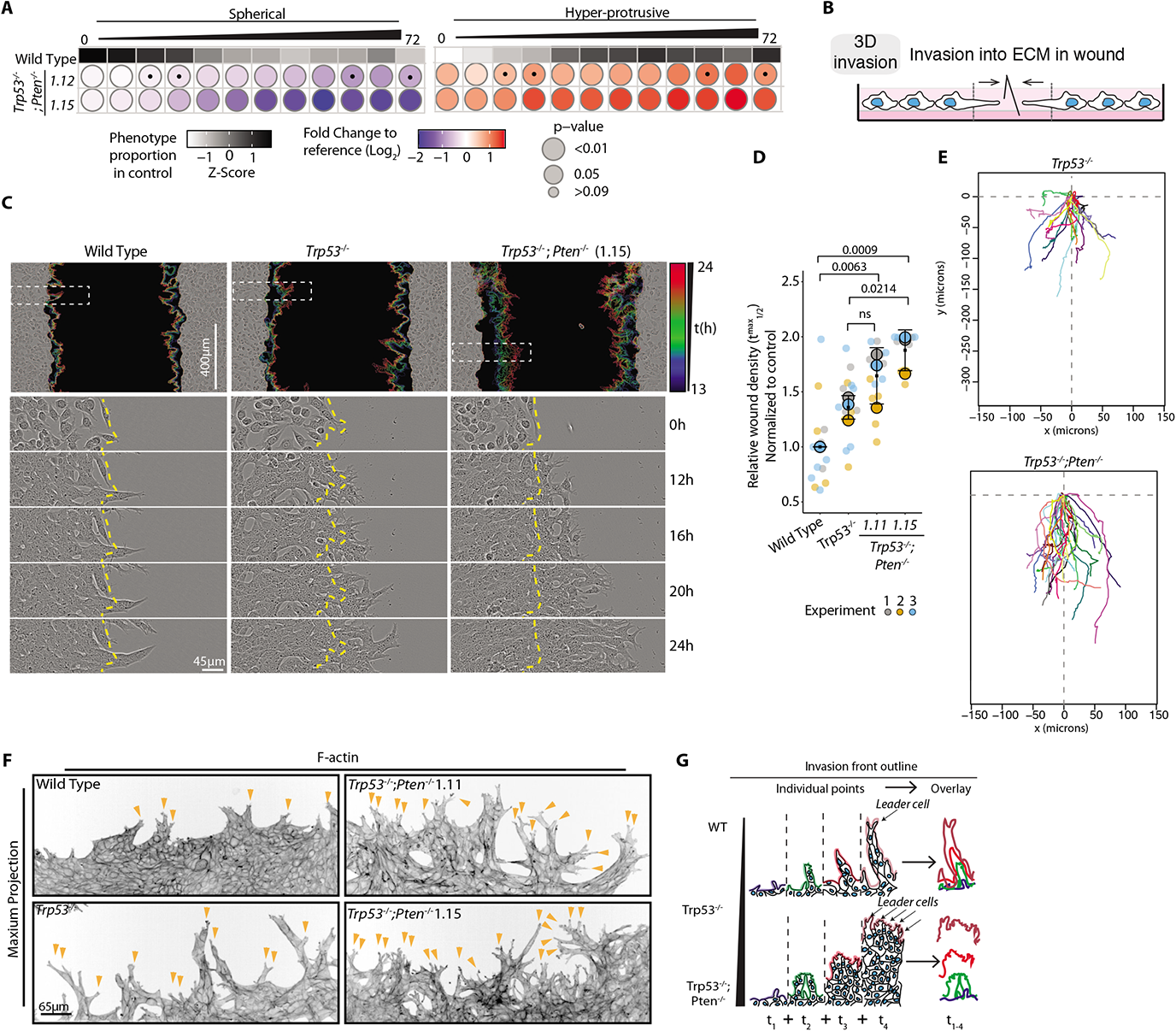
Collective invasion into ECM in orthogonal assays. **A.** Frequency, phenotype (Spherical, Hyper-protrusive), in ID8 WT, *Trp53^-/-^; Pten^-/-^* clones 1.12 and 1.15. spheroids. Grayscale heatmaps, phenotype proportion (z-score) in control (Wild-type (WT)). Blue-to-red heatmap, log_2_ fold change from control. P-values, bubble size (Cochran-Mantel-Haenszel test with Bonferroni adjustment). Black dot, homogenous effect across independent experiments (Breslow-Day test, Bonferroni adjustment, non-significant). N=3 experiments set up with repeated cultures of each sub-line, 3-4 technical replicates/subline in each experiment. Total spheroid number per condition, Supplementary Table 1. **B.** Schema, 3D invasion into ECM of wounded ID8 monolayer. **C.** Representative images, wounded monolayers invading ECM. Outlines of invasive front at different time points, pseudocoloured by time (rainbow legend), overlaid as concatenate on phase image of initial wound. Boxes, regions for different timepoints. Yellow lines, initial wound. Scale bar, 400μm or 45μm (indicated). **D.** Quantification of **C**. Graph, Relative Wound Density (RWD) at t_1/2_ max (time when WT 50% closed). Data, mean (black square) ± SD for 3 experiments set up with repeated cultures of each sub-lines (large circles) with 3-6 technical replicates/sub- line in each experiment (small circles). ANOVA with Tukey’s honest significant difference (HSD) test; exact p-values, annotated; ns, non-significant. **E.** Representative spider plots, leader cell movement in first 19 hr of invasion of *Trp53^-/-^*and *Trp53^-/^-;Pten^-/-^*1.15 ID8 cells. N=2 experiments set up with repeated cultures of each sub-lines, 10-25 leader cells tracked in each, across 4-6 technical replicates per sub-line in each experiment. **F.** Confocal images of wounded monolayer invasive fronts, stained for F-actin (Phalloidin). Representative of 7-12 fields imaged across n=3 experiments set up with repeated cultures of each sub-lines, 4 technical replicates/sub-line in each experiment. Scale bar, 65μm. Arrowheads, protrusion tips. **G.** Schema, loss of PTEN phenotype, a sheet-like mode of invasion with most ECM- abutting cells acting as ‘leader cells’, compared to leader and follower cell chains in WT.

**Figure S3.**
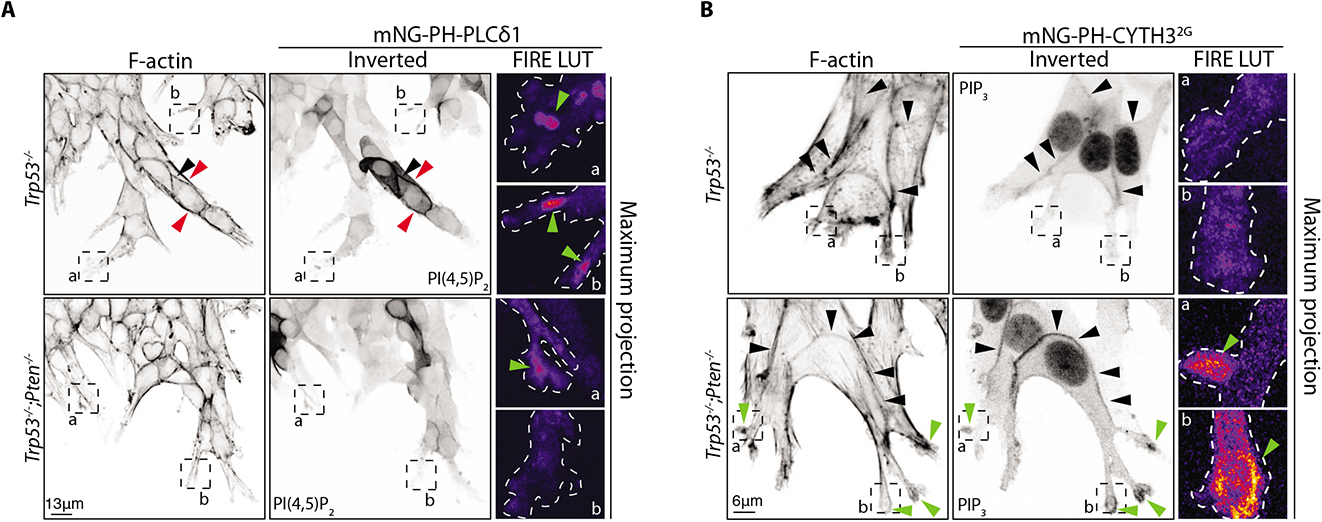
Characterisation of Phosphoinositide enrichment in Invasion assays. **A-B.** Confocal images, *Trp53^-/-^* or *Trp53^-/-^;Pten^-/-^* invasive monolayer fronts with cells expressing mNeonGreen (mNG) tagged biosensors for **(A)** PI(4,5)P_2_ (PH-PLCδ1) or **(B)** PIP_3_ (CYTH3^2G^). Representative of **(A)** 7 (*Trp53-/-*) or 9 (*Trp53^-/-^;Pten^-/-^*) fields or **(B)** 8 (*Trp53^-/-^*) or 9 (*Trp53^-/-^;Pten^-/-^*) fields imaged across n=2 experiments set up with repeated cultures of each sub-line. Magnified boxed regions, pseudocoloured with FIRE LUT. Arrowheads: cell-cell contacts, black; protrusions, green; cell-ECM contacts, red. Scale bar, **(A)**13 μm, **(B)** 6 μm.

**Figure S4.**
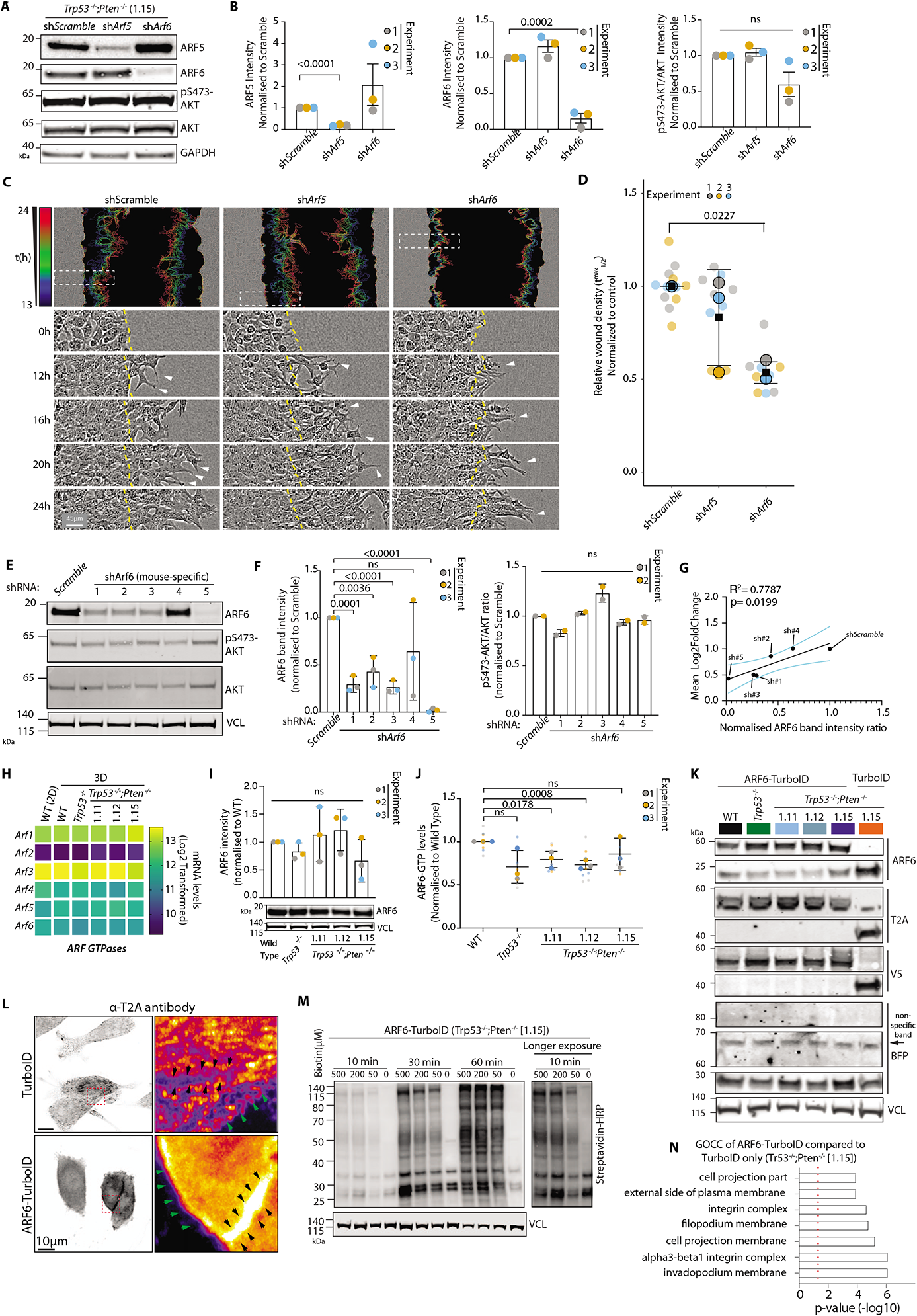
Further characterisation of ARF6 role upon *Trp53* and *Pten* loss. **A-B.** Western blot **(A)** and quantitation **(B)** from pS473-AKT, AKT, ARF5, ARF6, GAPDH in ID8 *Trp53^-/-^;Pten^-/-^*1.15 cell lines expressing shScramble, sh*Arf5* or sh*Arf6*. Representative blots of n=3 lysate preparations obtained from repeated cultures of each sub-line. **(B)** Data, mean ± SD for ARF5, ARF6 and pS473-AKT band intensity ratio, normalised to shScramble. P-values, unpaired, two-tailed t-test; ns, not significant. GAPDH is loading control for all panels. **C.** Representative images, ID8 *Trp53^-/-^;Pten^-/-^* 1.15 cell lines expressing sh*Scramble*, sh*Arf5* or sh*Arf6* in wounded monolayers invading ECM. Yellow lines, initial wound. Arrowheads, invasive protrusions. Outlines of invasive front pseudocoloured by time and overlaid as concatenate over phase image of initial wound. N=3 experiments set up using repeated cultures of each sub-line, 3-6 technical replicates/subline in each condition. Scale bar, 45μm. **D.** Quantitation of **C.** Graph, Relative Wound Density (RWD) at t_1/2_ max (time when shScramble 50% closed). Data, mean (black square) ± SD for 3 experiments set up using repeated cultures of each sub-line (large circles), 3-6 technical replicates/sub- line in each experiment (small circles). P-values, ANOVA with Tukey’s HSD test; annotated when significant. **E-F.** Western blot **(E)** and quantitation **(F)** from pS473-AKT, AKT, ARF6, VCL in ID8 *Trp53^-/-^;Pten^-/-^*1.15 cell lines expressing shScramble or sh*Arf6* (5 individual shRNA sequences). Representative blots of n=3 lysate preparations obtained from repeated cultures of each sub-line (ARF6) or n=2 (pS473-AKT and AKT) lysate preparations obtained from repeated cultures of each sub-line. VCL is loading control for all panels **(F)** Data, mean ± SD for ARF6 and pS473-AKT band intensity normalised to shScramble. P-values, unpaired, 2-tailed t tests; ns, not significant. **G.** Regression analysis. Scatter plot, mean Hyper-protrusive level across all time points versus ARF6 protein levels (determined by western blot). Solid black line, best linear fit and dotted cyan lines, 95% confidence interval. P-value and R^2^, annotated. **H.** Heatmap, Log_2_-transformed RNA-sequencing read counts of each ARF GTPase in ID8 spheroids and 2D monolayers (Wild Type, WT (2D)) across n=4 independent RNA preparations obtained from repeated cultures of each sub-line. **I.** Western blot and quantitation for ARF6 protein in ID8 sub-lines. VCL, loading control. Representative blots of n=3 independent protein isolations obtained from repeated cultures of each sub-line. Quantitation, mean ± SD ARF6 intensity normalised to ID8 WT. P-values, unpaired, two-tailed t-tests; ns, not significant. **J.** ARF6-GTP levels in ID8 sublines. Normalised Optical Density (OD) of Arf6-GTP G-LISA. N=3 lysate preparations obtained from repeated cultures of each sub-line, 3 technical replicates/sub-line in each condition. Data, mean ± SD of independent replicates (large circles) with technical replicates shown (small circles). P-values annotated, student’s T-test; ns, non-significant. **K.** Western blot, ARF6, T2A, V5, BFP and VCL from lysates extracted from TurboID (control) or ARF6-TurboID-expressing cell lines. VCL, loading control for T2A and BFP and sample integrity control for all other blots. N=3 independent lysate preparations. **L.** Confocal images, ID8 *Trp53*^-/-^*;Pten*^-/-^ 1.15 cells expressing ARF6-TurboID or TurboID, stained with T2A. Red box, cell-cell contacts, shown in higher magnification and pseudocoloured with FIRE LUT. Black arrowheads, cell-cell contact; green arrowheads, cell periphery. Representative images from 2 fields (ARF6-TurboID) or 3 fields (TurboID alone) from 1 experiment. Scale, 10μm. **M.** Western blot with Streptavidin HRP in ID8 *Trp53*^-/-^*;Pten*^-/-^ 1.15 cells treated with Biotin for at the times and concentrations indicated. VCL was used as loading control. n=1 lysate preparation. **N.** Gene Ontology Cell Compartment (GOCC) enrichment analysis of interactors identified in ID8 *Trp53*^-/-^*;Pten*^-/-^ 1.15 cells expressing ARF6-TurboID compared to TurboID alone. Data, p-value (-Log10) of enrichment. N=4 independent experiments. Red dotted line, significance threshold.

**Figure S5.**
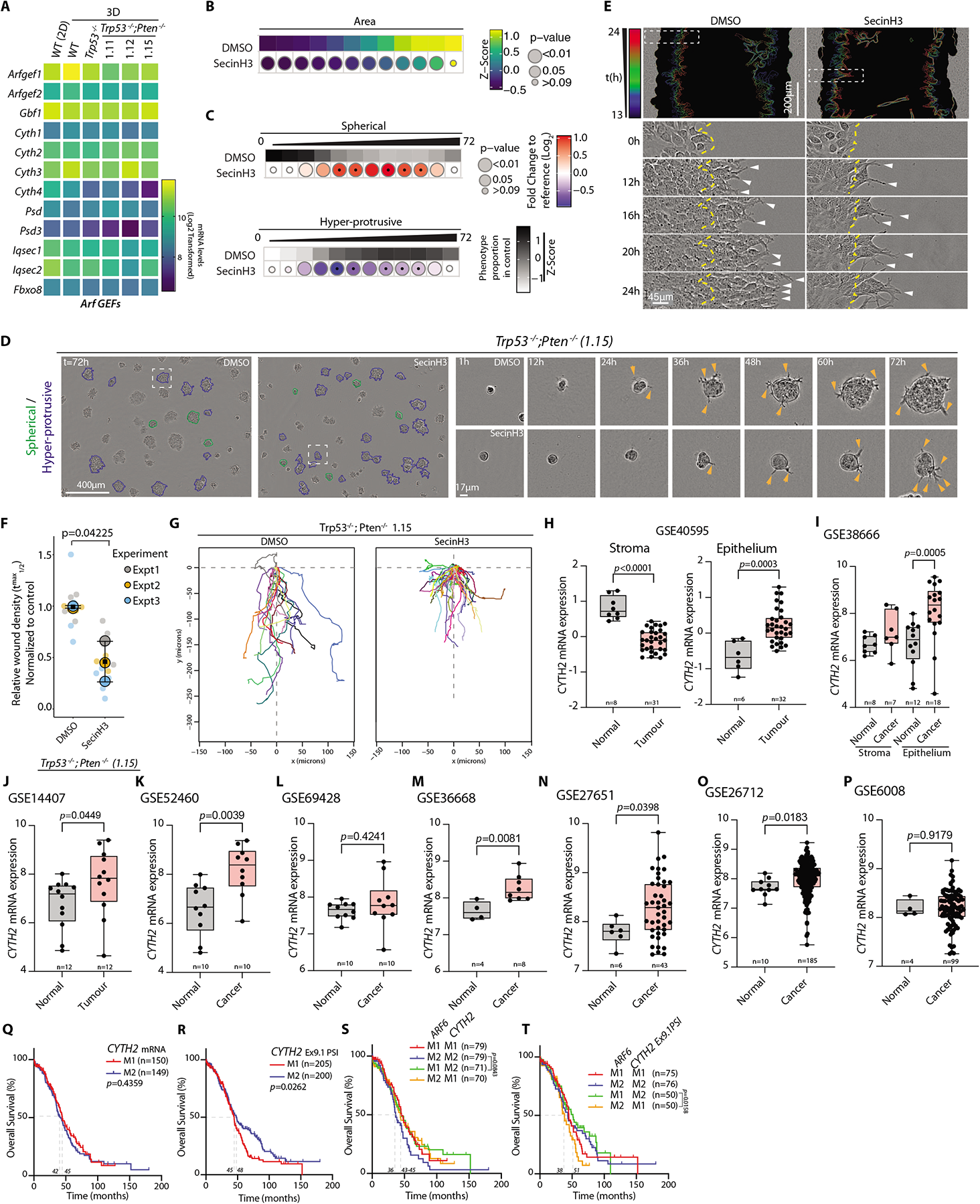
Characterisation of Cytohesin and CYTH2 contribution to invasion. **A.** Heatmap, log_2_-transformed RNA-sequencing read counts of ARF GEFs in ID8 spheroids and 2D monolayers (Wild Type, WT (2D)) across n=4 independent RNA preparations. **B-C.** ID8 *Trp53*^-/-^*;Pten*^-/-^ spheroids treated with 20μΜ SecinH3 quantified for **(B)** Area and **(C)** phenotype frequency (Spherical, Hyper-protrusive). Heatmaps, **(B)** (viridis colour scale) is mean of Z-score normalised values to control (DMSO) **(C)** Grayscale heatmaps, phenotype proportion (z-score) in control (DMSO). Blue-to-read heatmap, log_2_ fold change from control. P-values, bubble size **(B).** Student’s t-test, Bonferroni adjustment; **(C).** Cochran-Mantel-Haenszel test with Bonferroni adjustment). Black dot indicates homogenous effect across experiments (Breslow-Day test, Bonferroni adjustment, non-significant p-value). N=3 experiments set up using repeated cultures of each sub-line, 4-5 technical replicates/sub-line per experiment, total number of spheroids analysed in Supplementary Table 1. **D.** Representative phase contrast images of ID8 *Trp53*^-/-^*;Pten*^-/-^ treated with 20μΜ SecinH3, described in **B-C**. Outlines; Hyper-protrusive, blue; Spherical, green. Magnified spheroids shown for each condition at select time points. Arrowheads, protrusions into ECM. Scale bar, 400μm or 17μm (indicated). **E.** Representative images of ID8 Trp53^-/-^;Pten^-/-^ 1.15 wounded monolayers treated with DMSO or SecinH3 (20μΜ) in wounded monolayers invading ECM. Yellow lines, initial wound. Arrowheads, invasive protrusions. Outlines of invasive front pseudocoloured by time and overlaid as concatenate over phase image of initial wound. N=3 experiments set up using repeated cultures of each sub-line, 3-5 technical replicates per sub-line in each experiment. Scale bar, 200μm or 45μm (zoomed in boxed areas). **F.** Quantitation of **(E).** Graph, Relative Wound Density (RWD) at t_1/2_ max (time when DMSO 50% closed). Data, mean (black square) ± SD for 3 repeated experiments (large circles), 3-6 technical replicates per experiment (small circles). Exact p-value annotated, ANOVA with Tukey’s HSD test. **G.** Spider plots of leader cell movement in the first 19 hrs of invasion of *Trp53*^-/^-*;Pten*^-/-^ 1.15 ID8 cells treated with DMSO or SecinH3. 10-25 leader cells were tracked per experiment (n=2 set up using repeated cultures of each sub-line), across multiple technical replicates per experiment. Representative plots from cells tracked in 1 independent experiment shown. **H-P.** *CYTH2* mRNA levels in **(H, I)** laser-capture micro-dissected normal ovarian surface epithelium versus HGSOC epithelium or normal ovarian stroma versus ovarian cancer-associated stroma, or **(J-P)** bulk sequencing of normal ovary versus tumour. Specific dataset, sample size (n) and p-values (Mann-Whitney) annotated. **Q-T.** Overall survival (% patients, months; TCGA OV dataset), of patients grouped by low (M1) versus high (M2) levels based on a median split of **(Q)** *CYTH2* mRNA, **(R)** *CYTH2* exon 9.1 percentage spliced in ratio (PSI), **(S)** combination of *ARF6* and *CYTH2* mRNA, or **(T)** combination of A*RF6* mRNA and *CYTH2* Ex9.1 PSI. Median survival, sample size (n) and p-value, Log-rank test (Mantel-Cox) annotated.

**Figure S6.**
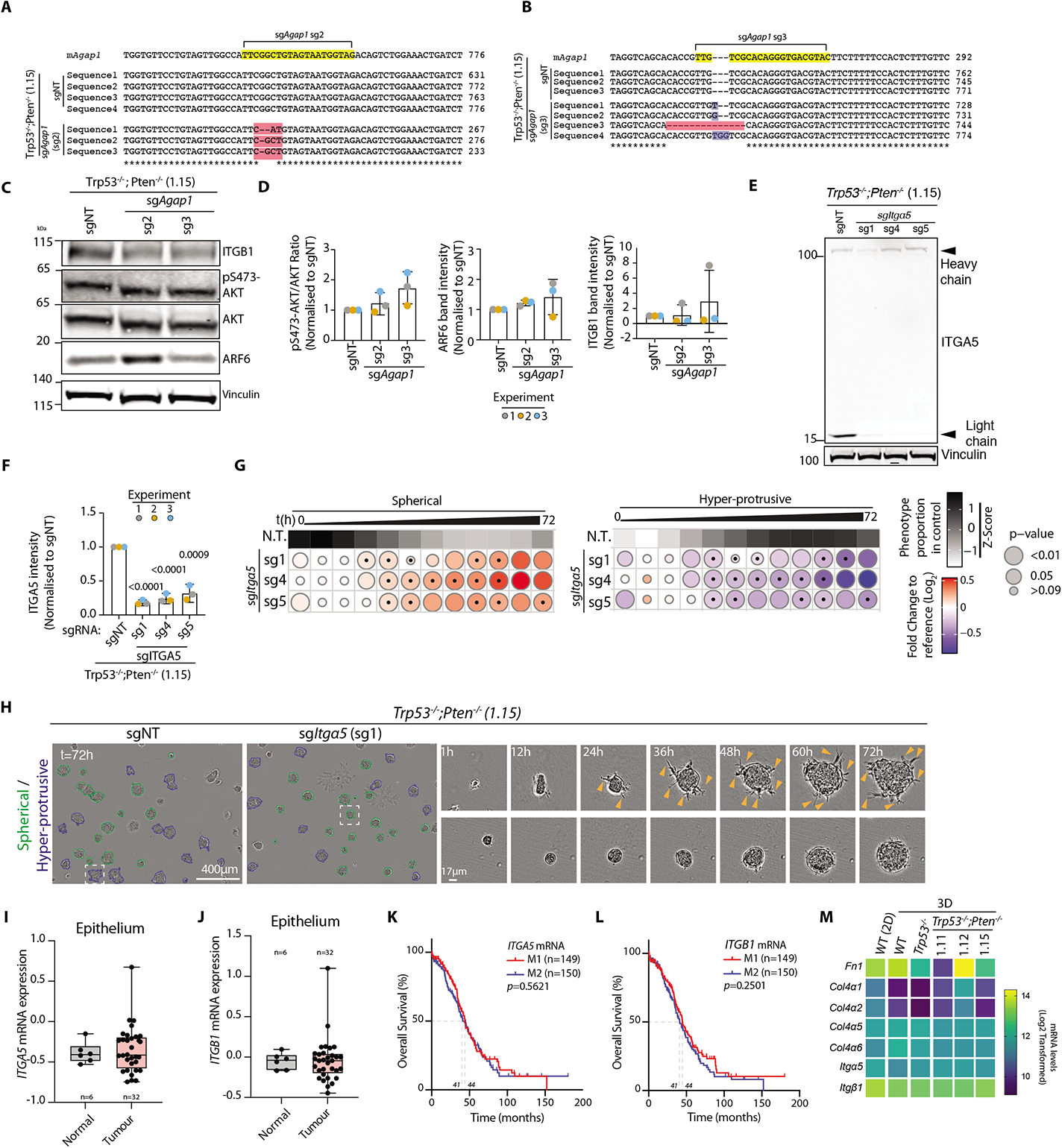
Itga5 contribution to collective invasion. **A-B**. Sequencing of CRISPR editing of the murine *Agap1* locus with sg2 **(A)** or sg3 **(B)** compared to control (sgNon-targeting, sgNT). Yellow, sgRNA sequence; red, deletions; blue, insertions. **C-D.** Western blotting **(C)** and quantitation **(D)** of β1-integrin (ITGB1), pS473-AKT, AKT and ARF6 in lysates from *Agap1* KO cells (sgNT versus sg2 or sg3). VCL is loading control for AKT, pS473-AKT and ARF6 and sample integrity control for ITGB1. Representative blots of n=3 lysate preparations obtained from repeated cultures of the sub-lines. **(D)** Data, mean ± SD for pS473-AKT:total AKT band intensity ratio, normalised to control (sgNT). Unpaired, two-tailed t-test, p values not significant (>0.05). **E-F.** Western blot **(E)** and quantitation (**F**), α5 integrin (ITGA5) in *ITGA5* KO cells (sg1, sg4, sg5 versus sgNT). VCL, loading control. Representative blots of n=3 lysate preparations obtained from repeated cultures of the sub-lines. **(F)** Data, mean ± SD sum intensity of ITGA5 heavy and light chain bands, normalised to ID8 Trp53*^-/^-;Pten^-/-^* 1.15 cells treated with sgNT. Unpaired, two-tailed t-test, p values are annotated. **G.** Frequency, spheroid phenotypes (Spherical, Hyper-protrusive) upon CRISPR-mediated KO of *Itga5*. Grayscale heatmaps, phenotype proportion (z-score) in control (WT). Blue-to-red heatmap, log_2_ fold change from control. P-values, bubble size (Cochran-Mantel-Haenszel test with Bonferroni adjustment). Black dot, homogenous effect across independent experiments (Breslow-Day test, Bonferroni adjustment, non-significant p-value). N=3 independent experiments set up using repeated cultures of the sub-lines, 3-6 technical replicates per sub-line in each condition. Number of spheroids analysed per condition in Supplementary Table 1. **H.** Representative phase contrast images of ID8 spheroids expressing sgNT or sg*Itga5* (sg1). Outlines: Hyper-protrusive, blue; Spherical, green. Magnified spheroids are shown at select time points. Arrowheads, protrusions into ECM. Scale bar 400μm or 17μm (indicated). **I, J.** *ITGA5* **(I)** and *ITGB1* **(J)** mRNA levels in laser-capture micro-dissected normal ovarian surface epithelium versus HGSOC epithelium. Dataset, GSE40595, sample size (n) and p-values (Mann-Whitney) annotated. **K, L.** Overall survival (% patients, months; TCGA OV dataset), of patients grouped by low (M1) versus high (M2) mRNA levels, based on a median split, of **(K)** *ITGA5* or **(L)** *ITGB1*. Median survival, sample size (n) and p-value, Log-rank test (Mantel-Cox) annotated. **M.** Heatmap, Log_2_-transformed RNA-sequencing read counts of Collagen IV, Fibronectin *Itg*α*5* and *Itgb1* in ID8 spheroids and 2D monolayers (Wild Type, WT (2D)) across n=4 independent RNA preparations.

**Figure S7.**
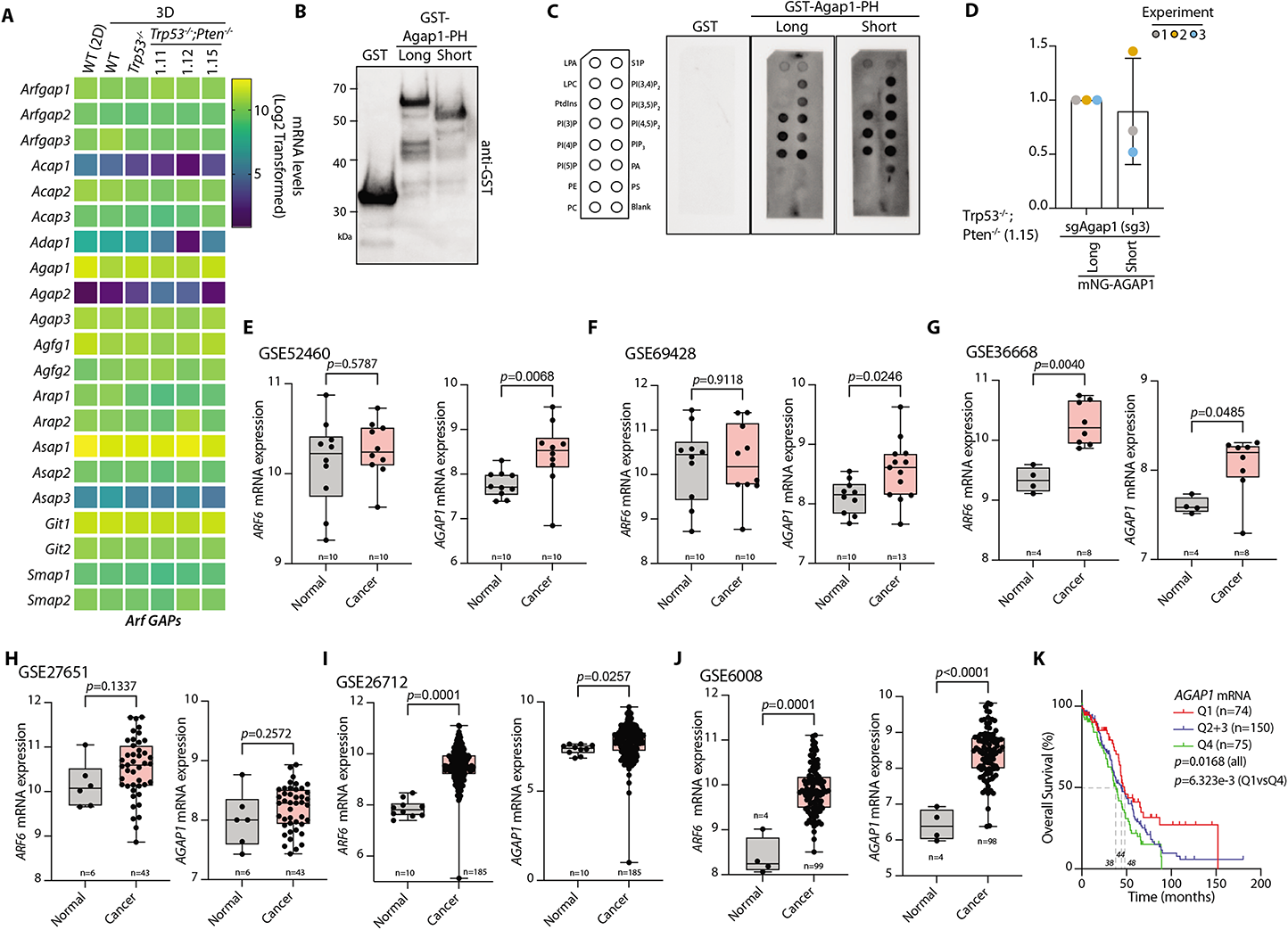
Further characterization of AGAP1 biological and clinical features. **A.** Heatmap, log_2_-transformed RNA-sequencing read counts of ARF GAPs in ID8 spheroids and 2D monolayers (Wild Type, WT (2D)) across n=4 independent RNA preparations. **B.** Western blot for GST performed on purified recombinant GST alone, or GST-fusion of both isoforms of AGAP1 PH domains. Representative of n=2 blots performed. **C.** Binding of the GST (Control), GST-AGAP1 PH Long or GST-AGAP1 PH Short fusion proteins to phospholipids immobilised on a cellulose membrane (cartoon on left denotes lipid position. Probed using a-GST antibody for visualization. Representative of n=3 experiments performed using an equal number of different aliquots of the purified recombinant proteins. LPA, Lysophosphatidic acid; LPC, Lysophosphocholine; PE, Phosphatidylethanolamine; PC, Phosphatidylcholine; S1P, Sphingosine-1-phosphate; PA, Phosphatidic Acid; PS, Phosphatidylserine. **D.** Quantitation of Figure **(6C)**. Data, mean ± SD of mNG band intensity between mNG-AGAP1-Long and mNG-AGAP1-Short isoforms. Unpaired, two-tailed t-test, p values ns, non-significant (>0.05). **E-J.** *ARF6 and AGAP1* mRNA levels in normal ovary versus tumour. Specific datasets, sample size (n) and p-values (Mann-Whitney) annotated. **K.** Overall survival (% patients, months; TCGA OV dataset), of patients grouped by *AGAP1* mRNA levels based on quartile. Median survival, sample size (n) and p-value, Log-rank test (Mantel-Cox) annotated.

**Figure S8.**
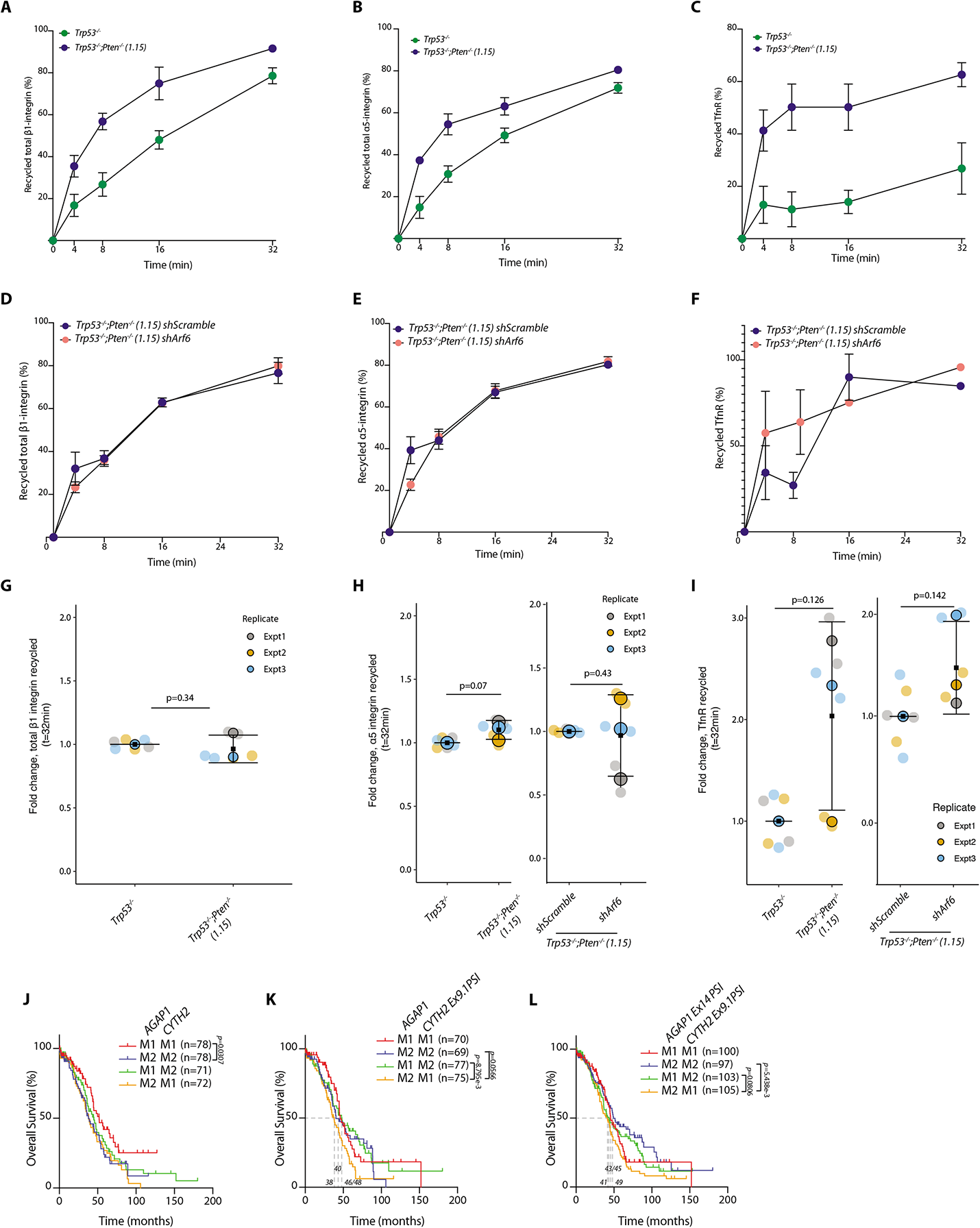
The ARF6 module does not affect recycling of other cargos. **A-I.** Representative capture ELISA graphs (A-F) and associated quantitation (**G-I**) for recycling of internalized cargoes. *Trp53^-/-^* versus *Trp53^-/-^;Pten^-/-^* cells (**A-C**) or *Trp53*^-/-^*;Pten*^-/-^ cells expressing sh*Scramble* versus sh*Arf6* (**D-F**). Graphs are shown for total β1 integrin (**A,D**), α5 integrin **(B,E)** or Transferrin receptor (TrnR) **(C,F)**. Graphs shown are representative of n=3 independent replicates apart from total β1 integrin in *Trp53*^-/-^*;Pten*^-/-^ cells expressing sh*Scramble* versus sh*Arf6,* for which *o*nly one experiment was performed. Data, mean (black square) ± SD for 3 repeated experiments (large circles), 1-3 technical replicates per experiment per timepoint (small circles), two-tailed t-test, p-values are annotated. **J-L.** Overall survival (% patients, months; TCGA OV dataset) of patients grouped in combinations of a median split of (**J**) *AGAP1* mRNA and *CYTH2* mRNA, (**K**) *AGAP1* mRNA and *CYTH2* Ex9.1 PSI, (**L**) *AGAP1* Ex14 PSI and *CYTH2* Ex9.1 PS. Median survival, sample size (n) and p-value, Log-rank test (Mantel-Cox) annotated.

The raw files and the MaxQuant search results files of the Mass Spectrometry experiment have been deposited to the ProteomeXchange Consortium ^65^ via the PRIDE partner repository ^66^ with the dataset identifier PXD038305. The RNA seq data have been deposited on the Sequence Read Archive (SRA), with ID Number PRJNA904679.

**Supplementary Table 1. Spheroid numbers per experimental condition.**

The number of spheroids (object) analysed for each condition in each Experimental replicate (live cyst imaging or 2D analysis of cell shape and pAKT intensity) using the Incucyte or the Opera Phenix systems respectively.

**Supplementary Table 2. Sequences of sgRNAs utilised.**

CRISPR guide RNA sequences used (written 5’-3’, Top and Bottom strand)

**Supplementary Table 3. Sequences of shRNAs utilised.**

shRNAs used, Target, species, target sequence and RNAi Consortium (TRC) clone ID provided.

**Supplementary Table 4. Details of molecular cloning.**

Cloning strategies for all constructs included in the study, including InFusion primer sequences where appropriate.

**Supplementary Table 5. Rules from CellProfiler Analyst for Classifier**

Rules used by CellProfiler to classify objects into “In Focus” or “Out of Focus” and then in “Smooth” or “Hyper-protrusive” spheroids.

**Supplementary Movie 1 Example of developing Smooth and Hyperprotrusive cysts**

Example of developing Smooth (left) and Hyperprotrusive (right) spheroids. Phase, live imaging, imaged every hour.

**Supplementary Movie 2 Developing cysts from ID8 CRISPR-derived cell lines**

Developing spheroids from ID8 CRISPR-derived cell lines, Wild Type (top left), *Trp53*^-/-^, (top right) *Trp53*^-/-^;*Pten*^-/-^ 1.11 (bottom left) and 1.15 (bottom right). Phase, live imaging, imaged every hour.

**Supplementary Movie 3 Developing cysts from ID8 CRISPR-derived Wild Type and *Trp53*^-/-^;*Pten*^-/-^ 1.12 cell lines**

Developing spheroids from ID8 CRISPR-derived cell lines, Wild Type (left), *Trp53*^-/-^, *Trp53*^-/-^;*Pten*^-/-^ 1.12 (right). Phase, live imaging, imaged every hour.

**Supplementary Movie 4 Invading monolayers of ID8 CRISPR-derived cell lines**

Developing monolayers from ID8 CRISPR-derived cell lines, Wild Type (top left), *Trp53*^-/-^, (top right) *Trp53*^-/-^;*Pten*^-/-^ 1.11 (bottom left) and 1.15 (bottom right). Phase, live imaging, imaged every hour.

**Supplementary Movie 5 Example ID8 *Trp53*^-/-^;*Pten*^-/-^ 1.15 spheroids treated with DMSO, AKT Inhibitor II or pan PI3K inhibitor (LY294002)**

Developing spheroids from *Trp53*^-/-^;*Pten*^-/-^ 1.5 cells treated with DMSO (left), AKT inhibitor II (middle), or PI3K pan-isoform inhibitor (LY294002, right). Phase, live imaging, imaged every hour.

**Supplementary Movie 6 Example ID8 *Trp53*^-/-^;*Pten*^-/-^ 1.15 spheroids treated with DMSO or PI3K isoform β inhibitor (AZD8186)**

Developing spheroids from *Trp53*^-/-^;*Pten*^-/-^ 1.15 cells treated with DMSO (left) or PI3K-β isoform inhibitor (AZD8186, right). Phase, live imaging, imaged every hour.

**Supplementary Movie 7 Example ID8 *Trp53*^-/-^;*Pten*^-/-^ 1.15 spheroids from cells stably expressing Scramble, *Arf5*- or *Arf6*-targeting shRNAs**

Developing spheroids from *Trp53*^-/-^;*Pten*^-/-^ 1.15 cells expressing sh*Scramble*, (left) or sh*Arf5* (middle), or sh*Arf6* (right). Phase, live imaging, imaged every hour.

**Supplementary Movie 8 Example ID8 *Trp53*^-/-^;*Pten*^-/-^ 1.15 spheroids with DMSO or SecinH3 treatment**

Developing spheroids from *Trp53*^-/-^;*Pten*^-/-^ 1.15 cells treated with DMSO (left) or SecinH3. Phase, live imaging, imaged every hour.

**Supplementary Movie 9 Example ID8 *Trp53*^-/-^;*Pten*^-/-^ 1.15 spheroids with sgNon-Targeting or sg*Agap1* #3**

Developing spheroids from *Trp53*^-/-^;*Pten*^-/-^ 1.15 cells expressing *Non-targeting*, (left) or *Agap1-targeting* (right) gRNAs. Phase, live imaging, imaged every hour.

**Supplementary Movie 10 Example ID8 *Trp53*^-/-^;*Pten*^-/-^ 1.15 sgNon-Targeting or sg*Itgb1* #4**

Developing spheroids from *Trp53*^-/-^;*Pten*^-/-^ 1.15 cells expressing *Non-targeting*, (left) or *Itgb1-targeting* (right) gRNAs. Phase, live imaging, imaged every hour.

