## Supplementary Table 1 for "PTEN deficiency exposes a requirement for an ARF GTPase module in integrin-dependent invasion in ovarian cancer"

**Supplementary Table 1_Spheroid Numbers per experiment**

The number of spheroids (object) analysed for each condition in each Experimental replicate (live cyst imaging or 2D analysis of cell shape and pAKT intensity) using the Incucyte or the Opera Phenix systems repectively.

| | | | |
|---|---|---|---|
| **Figure 1 Panels J-K** | | | |
| | **ID8 PTEN Expt1** | **ID8 PTEN Expt2** | **ID8 PTEN Expt3** |
| Condition | Spheroid number | Spheroid number | Spheroid number |
| **Wild Type** | 365 | 312 | 1468 |
| **Trp53-/-** | 127 | 677 | 1168 |
| **Trp53-/-;Pten-/- 1.11** | 663 | 439 | 801 |
| **Trp53-/-;Pten-/- 1.15** | 270 | 293 | 872 |
| **Figure 1 Panels L-M** | | | |
| | **PTENKO on WT Expt1** | **PTENKO on WT Expt2** | **PTENKO on WT Expt3** |
| Condition | Spheroid number | Spheroid number | Spheroid number |
| **PTEN_2** | 403 | 724 | 1617 |
| **PTEN_5** | 506 | 1333 | 1384 |
| **SCR2** | 490 | 1087 | 1786 |
| **Figure S1 Panel J** | | | |
| | **ID8 PTEN 112 Expt1** | **ID8 PTEN 112 Expt2** | **ID8 PTEN 112 Expt3** |
| Condition | Spheroid number | Spheroid number | Spheroid number |
| **PTEN 112** | 663 | 158 | 903 |
| **PTEN 115** | 712 | 762 | 829 |
| **Wild Type** | 1133 | 597 | 766 |
| **Figure 2 Panel D-F** | | | |
| | **ID8 PTEN 115 Inhibitors AKT/LY Expt1** | **ID8 PTEN 115 Inhibitors AKT/LY Expt2** | |
| Condition | Spheroid number | Spheroid number | |
| 10uM LY294002 | 701 | 1558 | |
| 15uM AKT Inhibitor II | 897 | 1824 | |
| DMSO | 544 | 1217 | |
| **Figure 2 Panel G-I** | | | |
| | **PI3K Isoform Inhibitors Expt1** | **PI3K Isoform Inhibitors Expt2** | |
| Condition | Spheroid number | Spheroid number | |

| | | | | |
|---|---|---|---|---|
| **DMSO** | 415 | 620 | | |
| **PI3Kalpha** | 676 | 877 | | |
| **PI3Kbeta** | 926 | 689 | | |
| **PI3Kdelta** | 811 | 723 | | |
| **PI3Kgamma** | 305 | 762 | | |
| **Figure S2 Panel A-B** | | | | |
| | **pAKT Intensity Expt1** | **pAKT Intensity Expt2** | **pAKT Intensity Expt3** | |
| Condition | Cell Number | Cell Number | Cell Number | |
| **Wild Type** | 1622 | 1622 | 1749 | |
| **Trp53-/-** | 1471 | 1144 | 1891 | |
| **Trp53-/-;Pten-/- 1.11** | 1274 | 1300 | 1749 | |
| **Trp53-/-;Pten-/- 1.15** | 3150 | 1096 | 1468 | |
| **Figure S2 Panel E-G** | | | | |
| | **2D Shape Expt1** | **2D Shape Expt2** | | |
| Condition | Cell Number | Cell Number | | |
| **Wild Type** | 7786 | 6594 | | |
| **Trp53-/-** | 6366 | 9840 | | |
| **Trp53-/-;Pten-/- 1.11** | 5972 | 4256 | | |
| **Trp53-/-;Pten-/- 1.15** | 7442 | 5933 | | |
| **Figure 3 Panel A-C** | | | | |
| | **Arf6 KD Expt1** | **Arf6 KD Expt2** | **Arf6 KD Expt3** | |
| Condition | Spheroid number | Spheroid number | Spheroid number | |
| **ARF5_3** | 1774 | 659 | 1361 | |
| **ARF6_3** | 1497 | 1014 | 667 | |
| **Scramble** | 1233 | 776 | 958 | |
| **Figure 3 Panel B** | | | | |
| | **CRISPR Screen Iteration 1 Expt1** | **CRISPR Screen Iteration 1 Expt2** | **CRISPR Screen Iteration 1 Expt3** | **CRISPR Screen Iteration 1 Expt4** |
| Condition | Spheroid number | Spheroid number | Spheroid number | Spheroid number |
| **EGFR** | 416 | 674 | 370 | 676 |
| **ITGA6** | 625 | 1239 | 711 | 913 |
| **ITGB1** | 586 | 1850 | 1650 | 860 |
| **LAMTOR5** | 342 | 1099 | 1230 | 1199 |
| **RAB14** | 514 | 847 | 763 | 987 |
| **SCR** | 607 | 1197 | 595 | 1037 |

| | | | | |
|---|---|---|---|---|
| **SCRIB** | 373 | 1516 | 1459 | 1072 |
| **YWHAQ** | 564 | 1167 | 1242 | 916 |
| | **CRISPR Screen Iteration 2 Expt1** | **CRISPR Screen Iteration 2 Expt2** | **CRISPR Screen Iteration 2 Expt3** | |
| Condition | Spheroid number | Spheroid number | Spheroid number | |
| **AGAP1** | 1327 | 1598 | 839 | |
| **ANO1** | 818 | 513 | 954 | |
| **CYTH2** | 1180 | 515 | 1228 | |
| **INPPL1** | 938 | 1602 | 1069 | |
| **ITGA3** | 1197 | 1468 | 929 | |
| **ITGA5** | 957 | 531 | 996 | |
| **SCR** | 1259 | 909 | 1182 | |
| **SPAG9** | 1157 | 1284 | 941 | |
| | CRISPR Screen Iteration 3 Expt1 | CRISPR Screen Iteration 3 Expt2 | CRISPR Screen Iteration 3 Expt3 | |
| Condition | Spheroid number | Spheroid number | Spheroid number | |
| **ARHGEF2** | 770 | 849 | 637 | |
| **FERMT2** | 1533 | 926 | 812 | |
| **FLOT2** | 1103 | 844 | 595 | |
| **KRT8** | 810 | 279 | 517 | |
| **LAMTOR3** | 1914 | 895 | 752 | |
| **LGR4** | 1039 | 1255 | 1135 | |
| **SCR** | 560 | 1084 | 916 | |
| **TPD52L2** | 1313 | 1423 | 982 | |
| | **CRISPR Screen Iteration 4 Expt1** | **CRISPR Screen Iteration 4 Expt2** | **CRISPR Screen Iteration 4 Expt3** | |
| Condition | Spheroid number | Spheroid number | Spheroid number | |
| **FLRT2** | 399 | 436 | 862 | |
| **FMNL3** | 491 | 719 | 858 | |
| **IGF2R** | 440 | 676 | 1144 | |
| **MYOF** | 157 | 1120 | 1179 | |
| **PCBP2** | 433 | 1083 | 655 | |
| **RAB3B** | 261 | 848 | 716 | |
| **SCR** | 35 | 885 | 1114 | |
| **Figure 4 Panel E** | | | | |
| | **ITGB1 Deco Expt1** | **ITGB1 Deco Expt2** | **ITGB1 Deco Expt3** | |
| Condition | Spheroid number | Spheroid number | Spheroid number | |
| **ITGB1_2** | 617 | 599 | 1977 | |
| **ITGB1_3** | 1075 | 504 | 649 | |
| **ITGB1_4** | 764 | 665 | 1530 | |
| **SCR2** | 587 | 751 | 378 | |

| **Figure 4 Panel G** | | | |
|---|---|---|---|
| | **Agap1 Deco Expt1** | **Agap1 Deco Expt2** | **Agap1 Deco Expt3** |
| Condition | Spheroid number | Spheroid number | Spheroid number |
| **AGAP1_2** | 1272 | 952 | 341 |
| **AGAP1_3** | 1786 | 1159 | 171 |
| **SCR2** | 1379 | 890 | 299 |
| **Figure S4 Panel B-C** | | | |
| | **SecinH3 Expt1** | **SecinH3 Expt2** | **SecinH3 Expt3** |
| Condition | Spheroid number | Spheroid number | Spheroid number |
| **115 DMSO** | 924 | 584 | 664 |
| **115 SecinH3** | 197 | 856 | 569 |
| **Figure 5S Panel D** | | | |
| | **ID8 Itgr a5 KO Expt 1** | **ID8 Itgr a5 KO Expt 2** | **ID8 Itgr a5 KO Expt 3** |
| Condition | Spheroid number | Spheroid number | Spheroid number |
| **Itgr a5_1 KO** | 1129 | 1498 | 1178 |
| **Itgr a5_4 KO** | 977 | 932 | 951 |
| **Itgr a5_5 KO** | 1407 | 1422 | 749 |
| **Scr** | 1350 | 838 | 834 |
