## Supplementary Table 2 for "PTEN deficiency exposes a requirement for an ARF GTPase module in integrin-dependent invasion in ovarian cancer"

**Supplementary Table 2 gRNA Sequences**
CRISPR gRNA sequences used in the CRISPR Screen (b is bottom oligo strand) and throughout the paper.

| Sequence Name | Sequence (5'-3') |
|---|---|
| Non_Targeting_2 | CAC CGC CGC GCC GTT AGG GAA CGA G |
| Non_Targeting_2b | AAA CCT CGT TCC CTA ACG GCG CGG C |
| Non_Targeting_3 | CAC CGC GGG CGT CAC CTG CTA GTA A |
| Non_Targeting_3b | AAA CTT ACT AGC AGG TGA CGC CCG C |
| Non_Targeting_4 | CAC CGT ACA GTT ATA CGT CGC GGT G |
| Non_Targeting_4b | AAA CCA CCG CGA CGT ATA ACT GTA C |
| Non_Targeting_6 | CAC CGC TCG GGC TAT TCA GCG ATA G |
| Non_Targeting_6b | AAA CCT ATC GCT GAA TAG CCC GAG C |
| Pten_2 | CAC CGG GTT TGA TAA GTT CTA GCT G |
| Pten_2b | AAA CCA GCT AGA ACT TAT CAA ACC |
| Pten_5 | CAC CGT CAC CTG GAT TAC AGA CCC G |
| Pten_5b | AAA CCG GGT CTG TAA TCC AGG TGA |
| Itga6_1 | CAC CGG AGT ATA TAT TTG ACG GAG A |
| Itga6_1b | AAA CTC TCC GTC AAA TAT ATA CTC C |
| Itga6_2 | CAC CGT CAA ATT CAA TCC GTG TAC A |
| Itga6_2b | AAA CTG TAC ACG GAT TGA ATT TGA C |
| Itga6_3 | CAC CGC TCC TGA GTA TAC TTT GGC T |
| Itga6_3b | AAA CAG CCA AAG TAT ACT CAG GAG C |
| Itga6_4 | CAC CGC AAC ACG AAG CAG GAG TCG C |
| Itga6_4b | AAA CGC GAC TCC TGC TTC GTG TTG C |
| Itga6_5 | CAC CGC AAT CCG TGT ACA AGG TCC C |
| Itga6_5b | AAA CGG GAC CTT GTA CAC GGA TTG C |
| Itgb1_1 | CAC CGG TCA CAG GAT CGA CCT GCA C |
| Itgb1_1b | AAA CGT GCA GGT CGA TCC TGT GAC C |
| Itgb1_2 | CAC CGA AGA CAT GGA CGC TTA CTG C |
| Itgb1_2b | AAA CGC AGT AAG CGT CCA TGT CTT C |
| Itgb1_3 | CAC CGG AGG AAT GTA ACA CGA CTG C |
| Itgb1_3b | AAA CGC AGT CGT GTT ACA TTC CTC C |
| Itgb1_4 | CAC CGG CGG AGA ATG TAT ACA AGC A |
| Itgb1_4b | AAA CTG CTT GTA TAC ATT CTC CGC C |
| Itgb1_5 | CAC CGG AGG AAT GTA ACA CGA CTG C |
| Itgb1_5b | AAA CGC AGT CGT GTT ACA TTC CTC C |
| Krt8_1 | CAC CGC CAG CTT CAA GGG GCT CAA C |
| Krt8_1b | AAA CGT TGA GCC CCT TGA AGC TGG C |
| Krt8_2 | CAC CGG CTA CAT CAA CAA CCT CCG C |
| Krt8_2b | AAA CGC GGA GGT TGT TGA TGT AGC C |

| | |
|---|---|
| Krt8_3 | CAC CGC CGA CGA GAT CAA CTT CCT C |
| Krt8_3b | AAA CGA GGA AGT TGA TCT CGT CGG C |
| Krt8_4 | CAC CGT GAA GTT CGT GCC CAG TAC G |
| Krt8_4b | AAA CCG TAC TGG GCA CGA ACT TCA C |
| Krt8_5 | CAC CGC CTC CGG CAG ATC CAT GAA G |
| Krt8_5b | AAA CCT TCA TGG ATC TGC CGG AGG C |
| Lamtor3_1 | CAC CGA CAT TAC TCG TGA CCT CCG C |
| Lamtor3_1b | AAA CGC GGA GGT CAC GAG TAA TGT C |
| Lamtor3_2 | CAC CGC ACA TTA CTC GTG ACC TCC G |
| Lamtor3_2b | AAA CCG GAG GTC ACG AGT AAT GTG C |
| Lamtor3_3 | CAC CGG ACT GAA CTG CGG GAA CAC C |
| Lamtor3_3b | AAA CGG TGT TCC CGC AGT TCA GTC C |
| Lamtor3_4 | CAC CGA CTG AAC TGC GGG AAC ACC G |
| Lamtor3_4b | AAA CCG GTG TTC CCG CAG TTC AGT C |
| Lamtor3_5 | CAC CGA CGG AAT ATA CAG ACC TGG T |
| Lamtor3_5b | AAA CAC CAG GTC TGT ATA TTC CGT C |
| Lamtor5_1 | CAC CGA GTC ACG TGA TCG AGG TTT G |
| Lamtor5_1b | AAA CCA AAC CTC GAT CAC GTG ACT C |
| Lamtor5_2 | CAC CGA CCC TGT CGG ATG AGC ACG C |
| Lamtor5_2b | AAA CGC GTG CTC ATC CGA CAG GGT C |
| Lamtor5_3 | CAC CGC ATA CCA CAG GGA TGT CGG T |
| Lamtor5_3b | AAA CAC CGA CAT CCC TGT GGT ATG C |
| Lamtor5_4 | CAC CGT CGA TCA CGT GAC TTG ACC G |
| Lamtor5_4b | AAA CCG GTC AAG TCA CGT GAT CGA C |
| Lgr4_1 | CAC CGT AAG TTG ATT GAG CCC ATT C |
| Lgr4_1b | AAA CGA ATG GGC TCA ATC AAC TTA C |
| Lgr4_2 | CAC CGT TAA CCT CAT TTC CTA CGG A |
| Lgr4_2b | AAA CTC CGT AGG AAA TGA GGT TAA C |
| Lgr4_3 | CAC CGA TTA AGT GTT TGC ACG ATA C |
| Lgr4_3b | AAA CGT ATC GTG CAA ACA CTT AAT C |
| Lgr4_4 | CAC CGC AGG ATT TCT ACT ACG ACT G |
| Lgr4_4b | AAA CCA GTC GTA GTA GAA ATC CTG C |
| Lgr4_5 | CAC CGC AAC CAT ATT ACC TCA GTC C |
| Lgr4_5b | AAA CGG ACT GAG GTA ATA TGG TTG C |
| Myof_1 | CAC CGC GGA GTC CAC GCA GGC CCA C |
| Myof_1b | AAA CGT GGG CCT GCG TGG ACT CCG C |
| Myof_2 | CAC CGT GGC TCG GAG GAT CAC CAA A |
| Myof_2b | AAA CTT TGG TGA TCC TCC GAG CCA C |
| Myof_3 | CAC CGC TGG CTC GGA GGA TCA CCA A |
| Myof_3b | AAA CTT GGT GAT CCT CCG AGC CAG C |
| Myof_4 | CAC CGA GAG GAC ACC TAT ACG GAT G |
| Myof_4b | AAA CCA TCC GTA TAG GTG TCC TCT C |

| | |
|---|---|
| Myof_5 | CAC CGG TTT CTC TCC CAG ATA CAC G |
| Myof_5b | AAA CCG TGT ATC TGG GAG AGA AAC C |
| Pcbp2_1 | CAC CGG ACA CCG GTG TGA TTG AAG G |
| Pcbp2_1b | AAA CCC TTC AAT CAC ACC GGT GTC C |
| Pcbp2_2 | CAC CGA TGG ACA CCG GTG TGA TTG A |
| Pcbp2_2b | AAA CTC AAT CAC ACC GGT GTC CAT C |
| Pcbp2_3 | CAC CGG CAC GTA TCA ACA TCT CAG A |
| Pcbp2_3b | AAA CTC TGA GAT GTT GAT ACG TGC C |
| Pcbp2_4 | CAC CGA CTG AAT CCG GTG TTG CCA T |
| Pcbp2_4b | AAA CAT GGC AAC ACC GGA TTC AGT C |
| Rab14_1 | CAC CGG TTA CAC GGA GCT ACT ATA G |
| Rab14_1b | AAA CCT ATA GTA GCT CCG TGT AAC C |
| Rab14_2 | CAC CGC ATA TAA CCA CTT AAG CAG C |
| Rab14_2b | AAA CGC TGC TTA AGT GGT TAT ATG C |
| Rab14_3 | CAC CGG GGC CGG ACC ACA AAA GAA G |
| Rab14_3b | AAA CCT TCT TTT GTG GTC CGG CCC C |
| Rab14_4 | CAC CGT GGG CCG GAC CAC AAA AGA A |
| Rab14_4b | AAA CTT CTT TTG TGG TCC GGC CCA C |
| Rab14_5 | CAC CGG CCG GAC CAC AAA AGA AGG G |
| Rab14_5b | AAA CCC CTT CTT TTG TGG TCC GGC C |
| Rab3b_1 | CAC CGA GCA TTG AAG GAC TCC TCG T |
| Rab3b_1b | AAA CAC GAG GAG TCC TTC AAT GCT C |
| Rab3b_2 | CAC CGC ATG TAT GAC ATC ACC AAC G |
| Rab3b_2b | AAA CCG TTG GTG ATG TCA TAC ATG C |
| Rab3b_3 | CAC CGT CAG ATT AAG ACC TAC TCC T |
| Rab3b_3b | AAA CAG GAG TAG GTC TTA ATC TGA C |
| Rab3b_4 | CAC CGC ATG TAT GAC ATC ACC AAC G |
| Rab3b_4b | AAA CCG TTG GTG ATG TCA TAC ATG C |
| Rab3b_5 | CAC CGG TAG TCA AAG TTC TGG TCG G |
| Rab3b_5b | AAA CCC GAC CAG AAC TTT GAC TAC C |
| Scrib_1 | CAC CGT ACA GAA GAC GAC TAT AAC G |
| Scrib_1b | AAA CCG TTA TAG TCG TCT TCT GTA C |
| Scrib_2 | CAC CGC TTG GAA GGA CCA TAC CCT G |
| Scrib_2b | AAA CCA GGG TAT GGT CCT TCC AAG C |
| Scrib_3 | CAC CGT TGT AAC CGG CAC GTG GAG T |
| Scrib_3b | AAA CAC TCC ACG TGC CGG TTA CAA C |
| Scrib_4 | CAC CGA CGT GGA GTC GGT GGA TAA G |
| Scrib_4b | AAA CCT TAT CCA CCG ACT CCA CGT C |
| Scrib_5 | CAC CGC CTG AAT GAC GTG TCC CTG C |
| Scrib_5b | AAA CGC AGG GAC ACG TCA TTC AGG C |
| Spag9_1 | CAC CGG TCC ATA TCC AAT TCA GGA G |
| Spag9_1b | AAA CCT CCT GAA TTG GAT ATG GAC C |

| | |
|---|---|
| Spag9_2 | CAC CGC ATT GAA TCC ACT CCT GAA T |
| Spag9_2b | AAA CAT TCA GGA GTG GAT TCA ATG C |
| Spag9_3 | CAC CGT TGA ACC TTT ATA TCC ACT G |
| Spag9_3b | AAA CCA GTG GAT ATA AAG GTT CAA C |
| Spag9_4 | CAC CGA AAG GAT TTA CAG ACG CGA G |
| Spag9_4b | AAA CCT CGC GTC TGT AAA TCC TTT C |
| Spag9_5 | CAC CGA ATC ACA GGA GAA GCA CCG T |
| Spag9_5b | AAA CAC GGT GCT TCT CCT GTG ATT C |
| Tpd52l2_1 | CAC CGC TTC TAC AAC TGG AGT CCG G |
| Tpd52l2_1b | AAA CCC GGA CTC CAG TTG TAG AAG C |
| Tpd52l2_2 | CAC CGG AGT CCG GTG GAC CAC ACC T |
| Tpd52l2_2b | AAA CAG GTG TGG TCC ACC GGA CTC C |
| Tpd52l2_3 | CAC CGG GAG TCC GGT GGA CCA CAC C |
| Tpd52l2_3b | AAA CGG TGT GGT CCA CCG GAC TCC C |
| Tpd52l2_4 | CAC CGT TGA CCC AGG TGT GGT CCA C |
| Tpd52l2_4b | AAA CGT GGA CCA CAC CTG GGT CAA C |
| Tpd52l2_5 | CAC CGT GGA CCA CAC CTG GGT CAA C |
| Tpd52l2_5b | AAA CGT TGA CCC AGG TGT GGT CCA C |
| Ywhaq_1 | CAC CGA CGG GAA GGT GCG GCA CCA A |
| Ywhaq_1b | AAA CTT GGT GCC GCA CCT TCC CGT C |
| Ywhaq_2 | CAC CGA CCA ACG GCC GCT TCC ATC T |
| Ywhaq_2b | AAA CAG ATG GAA GCG GCC GTT GGT C |
| Ywhaq_3 | CAC CGC CCA AGA TGG AAG CGG CCG T |
| Ywhaq_3b | AAA CAC GGC CGC TTC CAT CTT GGG C |
| Ywhaq_4 | CAC CGC CTC GTG CGC TGA GAC AAA G |
| Ywhaq_4b | AAA CCT TTG TCT CAG CGC ACG AGG C |
| Ywhaq_5 | CAC CGC ACC CGG CCC AAG ATG GAA G |
| Ywhaq_5b | AAA CCT TCC ATC TTG GGC CGG GTG C |
| Agap1_1 | CAC CGT GTA GCG TCA CTC CAC TGG T |
| Agap1_1b | AAA CAC CAG TGG AGT GAC GCT ACA C |
| Agap1_2 | CAC CGC TAC CAT TAC TAC AGC CGA A |
| Agap1_2b | AAA CTT CGG CTG TAG TAA TGG TAG C |
| Agap1_3 | CAC CGT ACG TCA CCC TGT GCG ACA A |
| Agap1_3b | AAA CTT GTC GCA CAG GGT GAC GTA C |
| Agap1_4 | CAC CGC ATA AGT TCC ACC AAC CCG A |
| Agap1_4b | AAA CTC GGG TTG GTG GAA CTT ATG C |
| Agap1_5 | CAC CGC ATA GTA CGT GCA CCT CTT C |
| Agap1_5b | AAA CGA AGA GGT GCA CGT ACT ATG C |
| Ano1_1 | CAC CGC TGC CCT TCT AAG TAA ACG G |
| Ano1_1b | AAA CCC GTT TAC TTA GAA GGG CAG C |
| Ano1_2 | CAC CGG ATG TTG GAC CGC ACA GAC G |
| Ano1_2b | AAA CCG TCT GTG CGG TCC AAC ATC C |

| | |
|---|---|
| Ano1_3 | CAC CGG ATG AGG CTC AAC TAC CGA T |
| Ano1_3b | AAA CAT CGG TAG TTG AGC CTC ATC C |
| Ano1_4 | CAC CGT GAC AAA CGC TTC AGA CGG G |
| Ano1_4b | AAA CCC CGT CTG AAG CGT TTG TCA C |
| Ano1_5 | CAC CGA CGG AAG GTG GAC TAC ATC T |
| Ano1_5b | AAA CAG ATG TAG TCC ACC TTC CGT C |
| Arhgef2_1 | CAC CGG CTC ACG CTG GAT TAT GTC T |
| Arhgef2_1b | AAA CAG ACA TAA TCC AGC GTG AGC C |
| Arhgef2_2 | CAC CGC AAT AAG GGA TTC CAC GGA C |
| Arhgef2_2b | AAA CGT CCG TGG AAT CCC TTA TTG C |
| Arhgef2_3 | CAC CGT GAG CAA GTC ACC CAA ACG A |
| Arhgef2_3b | AAA CTC GTT TGG GTG ACT TGC TCA C |
| Arhgef2_4 | CAC CGT GGA TGG TCA CGT TAC ATG C |
| Arhgef2_4b | AAA CGC ATG TAA CGT GAC CAT CCA C |
| Arhgef2_5 | CAC CGT GGC TAT GTC TCT TAC GAT G |
| Arhgef2_5b | AAA CCA TCG TAA GAG ACA TAG CCA C |
| Cyth2_1 | CAC CGG TAG ACA CCG TCC TCC ATG G |
| Cyth2_1b | AAA CCC ATG GAG GAC GGT GTC TAC C |
| Cyth2_2 | CAC CGG GCA AAG GAC AAC ACG TAG C |
| Cyth2_2b | AAA CGC TAC GTG TTG TCC TTT GCC C |
| Cyth2_3 | CAC CGA TGA ACC GGG GCA TCA ACG A |
| Cyth2_3b | AAA CTC GTT GAT GCC CCG GTT CAT C |
| Cyth2_4 | CAC CGC TGA GAT TCG GTA CAC CAT G |
| Cyth2_4b | AAA CCA TGG TGT ACC GAA TCT CAG C |
| Cyth2_5 | CAC CGA GCC ATT GGG GAC TAC CTA G |
| Cyth2_5b | AAA CCT AGG TAG TCC CCA ATG GCT C |
| Egfr_1 | CAC CGA CCA GAC AGT CAC TCT CTC G |
| Egfr_1b | AAA CCG AGA GAG TGA CTG TCT GGT C |
| Egfr_2 | CAC CGC ATG AAT AGG CCA ATC CCA A |
| Egfr_2b | AAA CTT GGG ATT GGC CTA TTC ATG C |
| Egfr_3 | CAC CGG AGA ACC TAG AAA TAA TAC G |
| Egfr_3b | AAA CCG TAT TAT TTC TAG GTT CTC C |
| Egfr_4 | CAC CGT GGG CCT GAC TAC TAC GAA G |
| Egfr_4b | AAA CCT TCG TAG TAG TCA GGC CCA C |
| Egfr_5 | CAC CGG ACC GCG AGA ACC ACA CTG C |
| Egfr_5b | AAA CGC AGT GTG GTT CTC GCG GTC C |
| Fermt2_1 | CAC CGA TCA TGA GCG TCG TAA GTA G |
| Fermt2_1b | AAA CCT ACT TAC GAC GCT CAT GAT C |
| Fermt2_2 | CAC CGA TTG AAG TAA TGT CAC CCT A |
| Fermt2_2b | AAA CTA GGG TGA CAT TAC TTC AAT C |
| Fermt2_3 | CAC CGA ATC AAT CAG CTT TAC GAG C |
| Fermt2_3b | AAA CGC TCG TAA AGC TGA TTG ATT C |

| | |
|---|---|
| Fermt2_4 | CAC CGC CAT CAT GAG CGT CGT AAG T |
| Fermt2_4b | AAA CAC TTA CGA CGC TCA TGA TGG C |
| Fermt2_5 | CAC CGC TGA TGA GCA GGT GTA CCA C |
| Fermt2_5b | AAA CGT GGT ACA CCT GCT CAT CAG C |
| Flot2_1 | CAC CGA TGG GCA ACT GCC ACA CGG T |
| Flot2_1b | AAA CAC CGT GTG GCA GTT GCC CAT C |
| Flot2_2 | CAC CGA GTG TCC GAG ATA CAC CAC C |
| Flot2_2b | AAA CGG TGG TGT ATC TCG GAC ACT C |
| Flot2_3 | CAC CGC CAC ACG GTG GGC CCC AAC G |
| Flot2_3b | AAA CCG TTG GGG CCC ACC GTG TGG C |
| Flot2_4 | CAC CGC CGA GAC CAG TTT GCC AAG C |
| Flot2_4b | AAA CGC TTG GCA AAC TGG TCT CGG C |
| Flot2_5 | CAC CGG TTC CTG GGC AAG AAC GTA C |
| Flot2_5b | AAA CGT ACG TTC TTG CCC AGG AAC C |
| Flrt2_1 | CAC CGC ACC ACC AAG GAA TGT GCG C |
| Flrt2_1b | AAA CGC GCA CAT TCC TTG GTG GTG C |
| Flrt2_2 | CAC CGA AGG TCC TGA GCA AGT CCG C |
| Flrt2_2b | AAA CGC GGA CTT GCT CAG GAC CTT C |
| Flrt2_3 | CAC CGC ACA ATA GTC GTC TTT CCG T |
| Flrt2_3b | AAA CAC GGA AAG ACG ACT ATT GTG C |
| Flrt2_4 | CAC CGA CAA CAT TCA GAC CAT CTC G |
| Flrt2_4b | AAA CCG AGA TGG TCT GAA TGT TGT C |
| Flrt2_5 | CAC CGC AGC TTA CGG AGG TTT GCG A |
| Flrt2_5b | AAA CTC GCA AAC CTC CGT AAG CTG C |
| Fmnl3_1 | CAC CGA GAC ATG GAC GTC ATC CTT C |
| Fmnl3_1b | AAA CGA AGG ATG ACG TCC ATG TCT C |
| Fmnl3_2 | CAC CGC ACT AAA GTC CTA CGG GAG C |
| Fmnl3_2b | AAA CGC TCC CGT AGG ACT TTA GTG C |
| Fmnl3_3 | CAC CGG GAG TCC ACT AAA GTC CTA C |
| Fmnl3_3b | AAA CGT AGG ACT TTA GTG GAC TCC C |
| Fmnl3_4 | CAC CGC ATT TGA CAA GCT CCG GTC C |
| Fmnl3_4b | AAA CGG ACC GGA GCT TGT CAA ATG C |
| Fmnl3_5 | CAC CGA CGG TGC ATT TGA CAA GCT C |
| Fmnl3_5b | AAA CGA GCT TGT CAA ATG CAC CGT C |
| Igf2r_1 | CAC CGG TTT ATA AGA TCA ACG TCT G |
| Igf2r_1b | AAA CCA GAC GTT GAT CTT ATA AAC C |
| Igf2r_2 | CAC CGA CGT TGG CCG ACC CAA GGA G |
| Igf2r_2b | AAA CCT CCT TGG GTC GGC CAA CGT C |
| Igf2r_3 | CAC CGT GAC GTT GGC CGA CCC AAG G |
| Igf2r_3b | AAA CCC TTG GGT CGG CCA ACG TCA C |
| Igf2r_4 | CAC CGA GGA CAT TGA CTC CAC ACG A |
| Igf2r_4b | AAA CTC GTG TGG AGT CAA TGT CCT C |

| | |
|---|---|
| Igf2r_5 | CAC CGC CAC TGA TCA AGC TAA ATG G |
| Igf2r_5b | AAA CCC ATT TAG CTT GAT CAG TGG C |
| Itga3_1 | CAC CGA TCG GTA CAC CAA GGT GCT G |
| Itga3_1b | AAA CCA GCA CCT TGG TGT ACC GAT C |
| Itga3_2 | CAC CGA CCC AGA ACA CCG TAT ACT T |
| Itga3_2b | AAA CAA GTA TAC GGT GTT CTG GGT C |
| Itga3_3 | CAC CGC TAC TAC TTC GAA CGG AAA G |
| Itga3_3b | AAA CCT TTC CGT TCG AAG TAG TAG C |
| Itga3_4 | CAC CGA TCG GTG ACA TCA ACC AGG A |
| Itga3_4b | AAA CTC CTG GTT GAT GTC ACC GAT C |
| Itga3_5 | CAC CGT AGT TCA TGG CAA TGA CGA T |
| Itga3_5b | AAA CAT CGT CAT TGC CAT GAA CTA C |
| Itga5_1 | CAC CGG TCT TAC ATA GCC ATA GGT G |
| Itga5_1b | AAA CCA CCT ATG GCT ATG TAA GAC C |
| Itga5_2 | CAC CGG TGT CAA GGA GCT GCC GAA T |
| Itga5_2b | AAA CAT TCG GCA GCT CCT TGA CAC C |
| Itga5_3 | CAC CGG GGC TTC AAC CTA GAC GCG G |
| Itga5_3b | AAA CCC GCG TCT AGG TTG AAG CCC C |
| Itga5_4 | CAC CGT GGG GGC TTC AAC CTA GAC G |
| Itga5_4b | AAA CCG TCT AGG TTG AAG CCC CCA C |
| Itga5_5 | CAC CGC TTC GAG GTG GAA CTC CAA C |
| Itga5_5b | AAA CGT TGG AGT TCC ACC TCG AAG C |
