## Supplementary Table 3 for "PTEN deficiency exposes a requirement for an ARF GTPase module in integrin-dependent invasion in ovarian cancer"

**Supplementary Table 3 shRNA Sequences**

shRNAs used, Target, species, target sequence and RNAi Consortium (TRC) clone ID provided.

| mRNA Target | Target Sequence | TRC ID |
|---|---|---|
| Scramble | CCGCAGGTATGCACGCGT | |
| Arf6_3 (human and mouse) | GCTCACATGGTTAACCTCTAA | TRCN0000048005 |
| Arf5_3 (human and mouse) | TGCTGATGAACTCCAGAAGAT | TRCN0000381650 |
| mARF6_1 (Mouse) | CCGGAAGGAGAGAAATCCAAA | TRCN0000100335 |
| mARF6_2 (Mouse) | CGGCAAGACAACGATCCTGTA | TRCN0000100336 |
| mARF6_3 (Mouse) | GCATTACTACACCGGGACCCA | TRCN0000100338 |
| mARF6_4 (Mouse) | CAACGTGGAGACGGTGACTTA | TRCN0000100339 |
| mARF6_5 (Mouse) | CAACGATCCTGTACAAGTTGA | TRCN0000381041 |
