## Supplementary Table 4 for "PTEN deficiency exposes a requirement for an ARF GTPase module in integrin-dependent invasion in ovarian cancer"

**Supplementary Table 4 Details of molecular cloning.**
Cloning strategies for all constructs included in the study, including InFusion primer sequences where appropriate.

| **Plasmid** | |
|---|---|
| CRISPR plasmids | All gRNA containing plasmids were constructed as described in Materials and Methods in a pLentiCRISPR V2 Neo backbone. The following sequences will be made available on Addgene upon publication: PTEN sgRNA2, PTEN sgRNA5, sgNon-targeting, sgAGAP1_2, sgAGAP1_3, sgITGB1_2, sgITGB1_3, sgITGB1_4, sgITGA5_1, sgITGA5_4, sgITGA5_5. All sequences are available in Supplementary Table 2. |
| shRNAs | All shRNAs were ordered in in the pLKO.puro backbone. All sequences and shRNA IDs are available in Supplementary Table 3. |
| pLX303 new MCS | A custom oligo was inserted beween the NheI and AgeI restriction sites of pLX303 |
| pLX304 mNG PH PLCδ1 | pLX303 new MCS was digested with NheI and AgeI. mNG was excised from mNeon-Green C1 using the AgeI and BsrGI cutting sites and PH domain of PLCδ1 was excised from pEGFP-NI-PH-PLCδ1 using NheI and AgeII. pLX303 mNG PH PLCδ1 was made through double ligation. |
| pLX303 mNG PH CYTH3(2G) | pLX303 new MCS was digested with NheI and BamHI. mNG was excised from mNeon-Green C1 using the NheI and BsrGI cutting sites and PH domain of CYTH3 (GRP1) was excised from pEGFP-CI-PH-GRP1 using BsrgI and BamHI. pLX303 mNG PH CYTH3 (2G) was made through double ligation. |
| pLX304 ARF6-R-WT mNG | pLX304 mNG-N1 was digested with NheI and BamHI and ARF6 was excised via PCR from pQCXIH-hARF6-R 2x myc WT (Fragment.FOR: 5' CGTCAGATCCGCTAGCGCCACCATGGGGAAGGTGCTATCC 3', Fragment.REV: 5' GGCGACCGGTGGATCCTCAGATTTGTAGTTGCTGGTCAGCC 3') and inserted to pLX304 mNG-N1 via InFusion reaction. |
| hARF6-R WT TurboID V5 T2A BFP | The V5-T2A-BFP sequence was ordered from Gene Art as an 899bp sequence. That and the pLX303 new MCS backbone were cut with NheI and AgeI and ligated together. The product was digested with ClaI and BsiWI and the TurboID sequence was ordered from Gene Art as a 990bp sequence and inserted to the backbone via InFusion cloning. The product was digested with NheI and BamHI and ARF6 was excised via PCR from pQCXIH-hARF6-R 2x myc WT (Arf6-R for Fragment.FOR 5'CGTCAGATCCGCTAGCGCCACCATGGGG 3', Fragment.REV 5'TTCCTGATCCGGATCCAGATTTGTAGTTGCTGGTCAGCCAGGT 3') and inserted into the digested backbon via InFusion reaction |
| Start Stuffer TurboID V5 T2A BFP | The steps were followed exactly as in the case of hARF6-R WT TurboID V5 T2A BFP, minus the last step in which a custom oligo was inserted instead of hARF6. (Top Strand 5' CTAGCAGCCACCATGGCCG 3', Bottom strand 5'GATCCGGCCATGGTGGCTG3'). |
| pGEX-4T1-TEV_Stop Codon | pGEX-4T1-TEV was polymerised mutagenesis primers to insert an end codon downstream the TEV site (Primer forward 5' TCAGGGATCCtgaGAATTCCCGG 3', Primer reverse 5'AAATACAGGTTTTCATCCG 3'. |
| pGEX-4T1-TEV_AGAP1 PH Domain Short/Long | Both PH Domains were ordered in from Gene Art ready for Infusion. The pGEX-4T1-TEV backbone was digested with BamHI and XhoI and the fragments inserted via InFusion reaction. |

| | |
|---|---|
| pLX303 mNG hAGAP1_Short/Long | Both sequences were ordered in from Gene Art ready for Infusion. The pLX303 mNG C1 new MCS backbone was digested with BsrGI and BamHI and a new, custom MCS was inserted. The product was digested with ClaI and PacI, and the GeneArt fragments inserted via InFusion reaction. |
