## Supplementary Table 5 for "PTEN deficiency exposes a requirement for an ARF GTPase module in integrin-dependent invasion in ovarian cancer"

**Supplementary Table 5_Rules for Classifier**

Rules used by CellProfiler to classify objects into “In Focusr” or “Out of Focus” and then in “Smooth” or “Hyper-protrusive” spheroids.

| **Rule to Classify out of Focus objects** |
|---|
| IF (AllCysts_Texture_DifferenceEntropy_ImageAfterMath_3_00_256 > 6.4043863878471221, [0.5665469353035022, -0.5665469353035022], [-0.53784614440892686, 0.53784614440892686])<br>IF (AllCysts_AreaShape_Solidity > 0.94251824817518237, [0.69190675699170856, -0.69190675699170856], [-0.26577431521079087, 0.26577431521079087])<br>IF (AllCysts_TrackObjects_Label_50 > 77.0, [-0.33159888087644496, 0.33159888087644496], [0.36244160487953964, -0.36244160487953964])<br>IF (AllCysts_Texture_Correlation_ImageAfterMath_3_03_256 > 0.13572703064094102, [-0.42845657779270052, 0.42845657779270052], [0.23083745049667539, -0.23083745049667539])<br>IF (AllCysts_Texture_AngularSecondMoment_ImageAfterMath_3_00_256 > 0.0010291413341912 56, [-0.50416522505083872, 0.50416522505083872], [0.23372189100484261, -0.23372189100484261])<br>IF (AllCysts_Texture_Correlation_ImageAfterMath_3_01_256 > 0.083321518091978533, [-0.24171142221650957, 0.24171142221650957], [0.28837904703132244, -0.28837904703132244])<br>IF (AllCysts_AreaShape_BoundingBoxMaximum_X > 241.0, [0.10953156003655966, -0.10953156003655966], [-0.57175061784657311, 0.57175061784657311])<br>IF (AllCysts_Texture_DifferenceEntropy_ImageAfterMath_3_00_256 > 6.6258822678025453, [0.64823250692809942, -0.64823250692809942], [-0.092352493862339013, 0.092352493862339013])<br>IF (AllCysts_Texture_SumVariance_ImageAfterMath_3_03_256 > 3020.0405237456835, [-0.25035010726665768, 0.25035010726665768], [0.21009517677429224, -0.21009517677429224])<br>IF (AllCysts_Texture_AngularSecondMoment_ImageAfterMath_3_02_256 > 0.0020694214876033055, [-0.97017639815037338, 0.97017639815037338], [0.052180912502166686, -0.052180912502166686])<br>IF (AllCysts_AreaShape_Solidity > 0.95967741935483875, [0.73145893012749053, -0.73145893012749053], [-0.086856010707660122, 0.086856010707660122])<br>IF (AllCysts_AreaShape_Zernike_9_9 > 0.0083870144082904882, [-0.45284456698834519, 0.45284456698834519], [0.1075479313423431, -0.1075479313423431])<br>IF (AllCysts_AreaShape_MeanRadius > 4.9831134122130605, [0.17818700527741363, -0.17818700527741363], [-0.21231812121146984, 0.21231812121146984])<br>IF (AllCysts_Texture_Correlation_ImageAfterMath_3_02_256 > 0.13178856322488719, [-0.15982683849251481, 0.15982683849251481], [0.33851278581872885, -0.33851278581872885])<br>IF (AllCysts_TrackObjects_Linearity_50 > 0.3838685343083385, [0.22360187354030153, -0.22360187354030153], [-0.14061180040909135, 0.14061180040909135])<br>IF (AllCysts_AreaShape_Zernike_4_4 > 0.028550436433074158, [-0.40149825845338033, 0.40149825845338033], [0.094850786343932458, -0.094850786343932458])<br>IF (AllCysts_Texture_Entropy_ImageAfterMath_3_03_256 > 12.385050159115206, [0.82893855165097441, -0.82893855165097441], [-0.044500456552775368, 0.044500456552775368])<br>IF (AllCysts_Texture_AngularSecondMoment_ImageAfterMath_3_00_256 > 0.0015575917369323269, [-0.45315993622524131, 0.45315993622524131], [0.075220126955717601, -0.075220126955717601]) |

IF (AllCysts_TrackObjects_IntegratedDistance_50 > 200.2816757973807, [-0.50716744783948275, 0.50716744783948275], [0.064822107720818062, -0.064822107720818062])
IF (AllCysts_TrackObjects_DistanceTraveled_50 > 1.4142135623730951, [0.15904629011536367, -0.15904629011536367], [-0.18443468884925132, 0.18443468884925132])

**Rule to Classify Spherical or Hyper-protrusive objects**

IF (InFocus_AreaShape_FormFactor > 0.66506490209176761, [0.85001124384269344, -0.85001124384269344], [-0.85362449077637181, 0.85362449077637181])
IF (InFocus_AreaShape_Extent > 0.67647058823529416, [0.73135767203147162, -0.73135767203147162], [-0.54740918255207627, 0.54740918255207627])
IF (InFocus_AreaShape_MinFeretDiameter > 35.355339059327378, [-0.40822338564616328, 0.40822338564616328], [0.74760864075243672, -0.74760864075243672])
IF (InFocus_AreaShape_Compactness > 1.2940048322465023, [-0.15931586225468428, 0.15931586225468428], [1.0, -1.0])
IF (InFocus_TrackObjects_ParentImageNumber_50 > 1515.0, [-0.13686475865785408, 0.13686475865785408], [0.67239867124838026, -0.67239867124838026])
IF (InFocus_AreaShape_Zernike_9_1 > 0.0058132023884139615, [-0.4354682702827945, 0.4354682702827945], [0.31205765567116739, -0.31205765567116739])
IF (InFocus_AreaShape_Solidity > 0.79548769814256537, [0.10665493023933746, -0.10665493023933746], [-1.0, 1.0])
IF (InFocus_AreaShape_Zernike_8_8 > 0.0081459922788837234, [-0.52845631472023513, 0.52845631472023513], [0.16678289577202082, -0.16678289577202082])
IF (InFocus_AreaShape_Zernike_4_4 > 0.012995089883204206, [-0.19511020862935685, 0.19511020862935685], [0.49711046905820389, -0.49711046905820389])
IF (InFocus_AreaShape_Zernike_4_4 > 0.032268888877573731, [0.72275202655379978, -0.72275202655379978], [-0.12130425445048572, 0.12130425445048572])
IF (InFocus_TrackObjects_Displacement_50 > 30.413812651491099, [-0.43893468815708503, 0.43893468815708503], [0.21644927729477936, -0.21644927729477936])
IF (InFocus_AreaShape_Zernike_5_5 > 0.011385200879590688, [-0.33785688082280235, 0.33785688082280235], [0.28447788528893858, -0.28447788528893858])
IF (InFocus_AreaShape_Extent > 0.62482517482517486, [0.25710826812964371, -0.25710826812964371], [-0.40648625053633075, 0.40648625053633075])
IF (InFocus_AreaShape_Zernike_2_0 > 0.17817473232737183, [-0.63332962732467357, 0.63332962732467357], [0.18534375919929169, -0.18534375919929169])
IF (InFocus_Parent_AllCysts > 39.0, [0.8672053026371167, -0.8672053026371167], [-0.14107847494030878, 0.14107847494030878])
IF (InFocus_AreaShape_Zernike_7_7 > 0.010714407572012651, [-0.61533872931413025, 0.61533872931413025], [0.18189578180627403, -0.18189578180627403])
IF (InFocus_AreaShape_Zernike_7_3 > 0.0045495327170735822, [-0.19065255055328276, 0.19065255055328276], [0.54495021694924806, -0.54495021694924806])
IF (InFocus_AreaShape_Zernike_8_2 > 0.0019553543396043471, [0.124194780405, -0.124194780405], [-0.89548680607592313, 0.89548680607592313])
IF (InFocus_AreaShape_Compactness > 1.2940048322465023, [-0.10688374130916414, 0.10688374130916414], [1.0, -1.0])
IF (InFocus_AreaShape_Zernike_2_2 > 0.089245133670850227, [0.89159884634238096, -0.89159884634238096], [-0.061963506016290006, 0.061963506016290006])
